# Divergent ontogeny of Tissue Resident Memory and Tissue Resident Exhausted CD8^+^ T cells underlies distinct functional potential

**DOI:** 10.1101/2025.08.08.669213

**Authors:** Simone L. Park, Mark M. Painter, Sasikanth Manne, Victor Alcalde, Maura McLaughlin, Matthew A. Sullivan, Divij Mathew, Leonel Torres, Yinghui J. Huang, David B. Reeg, Naomi R. Douek, Trenton Campos, Max Klapholz, Maria A. Cardenas, Victoria Fang, Shin Foong Ngiow, KC Wumesh, Rishi R. Goel, Amy E. Baxter, Jennifer E. Wu, Melody Tan, Corbett T. Berry, Christoph T. Ellebrecht, Alexander C. Huang, Emily Papazian, Ying Liu, Karthik Rajasekaran, Robert M. Brody, Erica R. Thaler, Devraj Basu, Ahmed Diab, Josephine R. Giles, E. John Wherry

## Abstract

Persistent antigen stimulation promotes differentiation of exhausted CD8^+^ T (T_EX_) cells. T_EX_ cells are distinct from circulating memory T (T_CIRCM_) cells but share many features with tissue-resident memory (T_RM_) cells established following infection resolution. CD8^+^ T cells co-expressing residency- and exhaustion-associated molecules in chronic diseases often correlate with clinical outcomes. However, the relationship between these cells and conventional T_RM_ or T_EX_ cells remains unclear. Here, we show that chronic antigen stimulation drives development of tissue-resident T_EX_ (TR-T_EX_) cells that are ontologically and functionally distinct from T_RM_ cells generated after antigen clearance. TR-T_EX_ phenotypically resembled T_RM_ cells but were regulated by distinct transcriptional networks and were uniquely dependent on Tox for residency programming. Although T_EX_ progenitor cells acquired residency features upon entering chronically infected tissues, they failed to generate conventional T_RM_ cells after antigen withdrawal. Conversely, T_RM_ cells were able to differentiate into T_EX_ cells during chronic antigen stimulation. Deriving cell-state specific transcriptional signatures revealed a selective association of TR-T_EX_ cells with patient responses to immune checkpoint blockade, and only TR-T_EX_ but not T_RM_ cells responded to PD-1 pathway inhibition in vivo. These data suggest that TR-T_EX_ and T_RM_ cells are developmentally distinct cell types that share a tissue-residency program but have distinct roles in disease control.

## Main

CD8^+^ T cell-mediated control of infection and cancer is central to the efficacy of immunotherapies. The CD8^+^ T cell compartment includes multiple cellular subsets with specialized roles in immune defense. Tissue-resident memory CD8^+^ (T_RM_) cells develop in peripheral non-lymphoid tissues following acutely resolved infections or vaccination, where they reside long-term without recirculating in blood. T_RM_ cells express molecules that promote tissue retention including CD69 and CD103, depend on residency-promoting transcription factors (TFs) including Blimp1, Hobit and Runx3 and are transcriptionally and epigenetically distinct from circulating memory T (T_CIRCM_) cells^1–6^. Like other memory T cells (T_MEM_), canonical T_RM_ cells formed after antigen clearance are long-lived, maintained in an antigen-independent manner, highly functional, and can provide rapid immune protection^7–11^. In contrast, during chronic infection or cancer, persistent antigen stimulation drives the development of CD8^+^ T_EX_ cells. These cells are defined by sustained inhibitory receptor (IR) expression, reduced or altered effector function, inefficient antigen-independent maintenance, and compromised developmental plasticity^12^. Underlying these properties, T_EX_ cells have a distinct epigenetic landscape compared to T_CIRCM_ that is programmed by the TF Tox^13–18^. The T_EX_ cell population is also heterogeneous and includes progenitor (T_EX-PROG_, also called stem-like) cells with increased plasticity, effector-like intermediate (T_EX-INT_), and terminally differentiated (T_EX-TERM_) cells^19^. Although T_EX_ cells are suboptimal in controlling disease, these cells – particularly proliferative T_EX_-_PROG_ – can be partially invigorated by immunotherapies such as PD-1 immune checkpoint blockade (ICB), often resulting in clinical benefit^14,20^.

Whereas T_EX_ cells are distinct from T_CIRCM_ cells^12,14^, the relationship between T_EX_ and T_RM_ cells has been less clear. Profiling antigen-specific CD8^+^ T cell responses in mouse models of vaccination, acute and chronic infection, or cancer, where the kinetics and duration of T cell receptor (TCR) stimulation can be precisely defined, has highlighted considerable phenotypic similarities between T_RM_ formed after antigen clearance and T_EX_ responding to persisting antigen. After resolution of acute infection, T_RM_ cells generated in mice express IRs including PD-1, CD39, the TF Tox and other exhaustion-associated features in the absence of persisting antigen^8,21–23^. On the other hand, chronic infections and cancer drive accumulation of non-recirculating CD8^+^ T cells in non-lymphoid tissues^24–27^ and solid tumors^28^, as well as establishment of non-migratory T_EX-PROG_ and T_EX-TERM_ cell populations in secondary lymphoid organs^25,29^. In these settings, tissue-residence is not only closely associated with high IR expression but also with expression of many T_RM_-associated molecules including CD69, Blimp1 and Runx3^28,29^. These shared features between T_RM_ and T_EX_ cells raise the possibility of potential overlap in their development and regulation.

CD8^+^ T cells co-expressing the tissue retention molecules CD69 and CD103 are widely present in healthy human tissues and in many chronic diseases, including human solid cancers^1^ where the presence of such cells often correlates with improved tumor control and immunotherapy responses^30^. Most CD69^+^CD103^+^ TIL co-express the T_RM_-associated TF Hobit (*ZNF683*) and multiple inhibitory receptors (IRs) including PD-1 and CD39^31–33^. These CD69^+^CD103^+^ TIL have a range of functional capacities in the tumor microenvironment (TME) ^34,35^, where they are often enriched for tumor-reactive T cell specificities^32,36,37^. Similar CD69^+^CD103^+^ CD8^+^ T cell populations are also prevalent in chronic autoimmune conditions including colitis^38^ and inflammatory skin disorders^39^. In these chronic diseases, CD8^+^ T cells expressing tissue-residency associated molecules are often classified as T_RM_ cells^40–45^. However, whether these cells are ontologically or operationally equivalent to canonical T_RM_ cells generated after resolution of acute infection or following vaccination is unclear. Key factors distinguishing antigen-independent T_RM_ from chronically stimulated T_EX_ cells remain poorly defined. Distinguishing between T_RM_ and T_EX_ cells is particularly challenging in human tissues or tumors where antigen and inflammation often persist, since the T cell receptor (TCR) specificity and the duration of antigen stimulation experienced by T cells in these environments is often unknown. As such, whether CD8^+^ T cells expressing tissue-residency features in these settings are developmentally related to T_RM_ cells generated after antigen clearance and possess functional properties typical of memory T cells, or whether these cells are partially or fully exhausted is unclear. Addressing these questions has implications for chronic disease treatment, as T_RM_ and T_EX_ cells may respond differently to therapeutic intervention.

To explore these issues, we compared CD8^+^ T cells residing in peripheral tissues after acute-resolving versus chronic lymphocytic choriomeningitis virus (LCMV) infection in mice. Chronic infection drove the differentiation of a population of CD8^+^ tissue-resident T_EX_ (TR-T_EX_) cells that expressed residency-associated molecules but was developmentally and functionally distinct from T_RM_ cells formed after antigen clearance. Whereas both T_RM_ and TR-T_EX_ cells relied on overlapping residency-promoting TFs such as Blimp1 and Runx3, as well Hobit in certain tissues, only TR-T_EX_ cells required Tox for residency programming and survival. T_RM_ cells retained plasticity to generate T_EX_ cells, whereas committed T_EX_ cells could not give rise to T_RM_ cells following antigen clearance and only acquired residency-associated features during chronic infection. T_RM_ and TR-T_EX_ cells engaged distinct gene regulatory networks, giving rise to cell-state specific transcriptional signatures that were stably maintained during inflammation. Finally, T_RM_ and TR-T_EX_ cells made distinct contributions to immunotherapy responses, with only TR-T_EX_ cells and not T_RM_ cells responding to PD-1 inhibition. These data reveal T_RM_ and TR-T_EX_ cells as developmentally distinct cellular lineages generated after clearance or persistence of antigen, respectively. Collectively, these findings highlight different roles for these cell types in disease settings and emphasize that immunotherapeutic interventions harnessing the unique properties of either cell subset could be developed to individually target T_RM_ versus T_EX_ cells.

## Results

### Phenotypically similar but functionally distinct CD8^+^ T cells develop in tissues during acute versus chronic infection

To explore the extent of phenotypic overlap between CD8^+^ T cells localizing to peripheral tissues after infection resolution or chronic infection, we compared expression of hallmark T_RM_ and T_EX_ associated molecules by virus-specific CD8 T cells responding to either acute-resolving (Armstrong, Arm) or chronic (clone 13, Cl13) strains of LCMV. Congenically marked (CD45.1^+^) P14 CD8^+^ T cells transgenic for a TCR recognizing the H-2D^b^-restricted gp_33-41_ epitope of LCMV were adoptively transferred to naïve mice prior to Arm or Cl13 infection. Parabiosis studies have shown that following resolution of Arm infection, the majority of tissue-localized P14 cells are non-recirculating T_RM_ cells that can be distinguished from T_CIRCM_ cells by a distinct surface phenotype that includes co-expression of the retention-coordinating molecules CD69, CXCR6, CD49a (VLA-1) and, in epithelial locations, CD103^1,27,46–48^. Here we used co-expression of CD69 and CD103 in epithelial sites (small intestine epithelium (SI) and salivary gland (SG)), or co-expression of CD69 and CXCR6 in non-epithelial tissues (liver and kidney) as proxies to identify ‘Arm T_RM_ cells’, which we defined as non-migratory CD8^+^ T cells generated in tissues after antigen clearance. P14 cells recruited into non-lymphoid tissues during chronic Cl13 infection are also largely non-recirculating^25,27^. Using the LCMV system, we could therefore explore how the nature and duration of antigen stimulation impacts tissue-resident T cell programming by comparing Arm T_RM_ cells generated after viral clearance to CD8^+^ T cells from matched non-lymphoid organs during chronic Cl13 infection.

Four weeks post-infection, P14 cells had accumulated at similar or higher numbers in Cl13-infected tissues, where viral titers remained high, compared to tissues from mice that had cleared Arm infection (**Extended Data Fig 1a-b**). Across all peripheral tissues examined, both Arm and Cl13-derived P14 cells upregulated expression of the residency-associated molecules CD69 and CXCR6 compared to their splenic counterparts, although CD69 and CXCR6 were also co-expressed by a subset of terminally exhausted (T_EX-TERM_) P14 cells in the spleen of Cl13 infected mice^29^ (**Fig 1a** and **Extended Data Fig 1c**). Whereas the T_RM_-associated marker CD103 was not expressed by CD69^+^ P14 cells in the spleen and liver, CD69^+^CD103^+^ P14 cells were abundant in peripheral tissues including the small intestine epithelium (SI), salivary gland (SG), and kidney (Kid) at day 30 after both Arm and Cl13 infection (**Fig 1b-c** and **Extended Data Fig 1c**). Although CD69^+^CD103^+^ P14 cells were present at lower frequencies following chronic compared to acutely resolved infection in some tissues including the SI^26,47,49,50^, the frequencies of these cells in other organs were comparable (SG) or even higher (Kid) at day 30 of chronic infection (**Fig 1b** and **c**). Thus, similar populations of CD69^+^CXCR6^+^CD103^+/–^ cells were found in non-lymphoid tissues after either acute or chronic infection.

**Figure 1.**
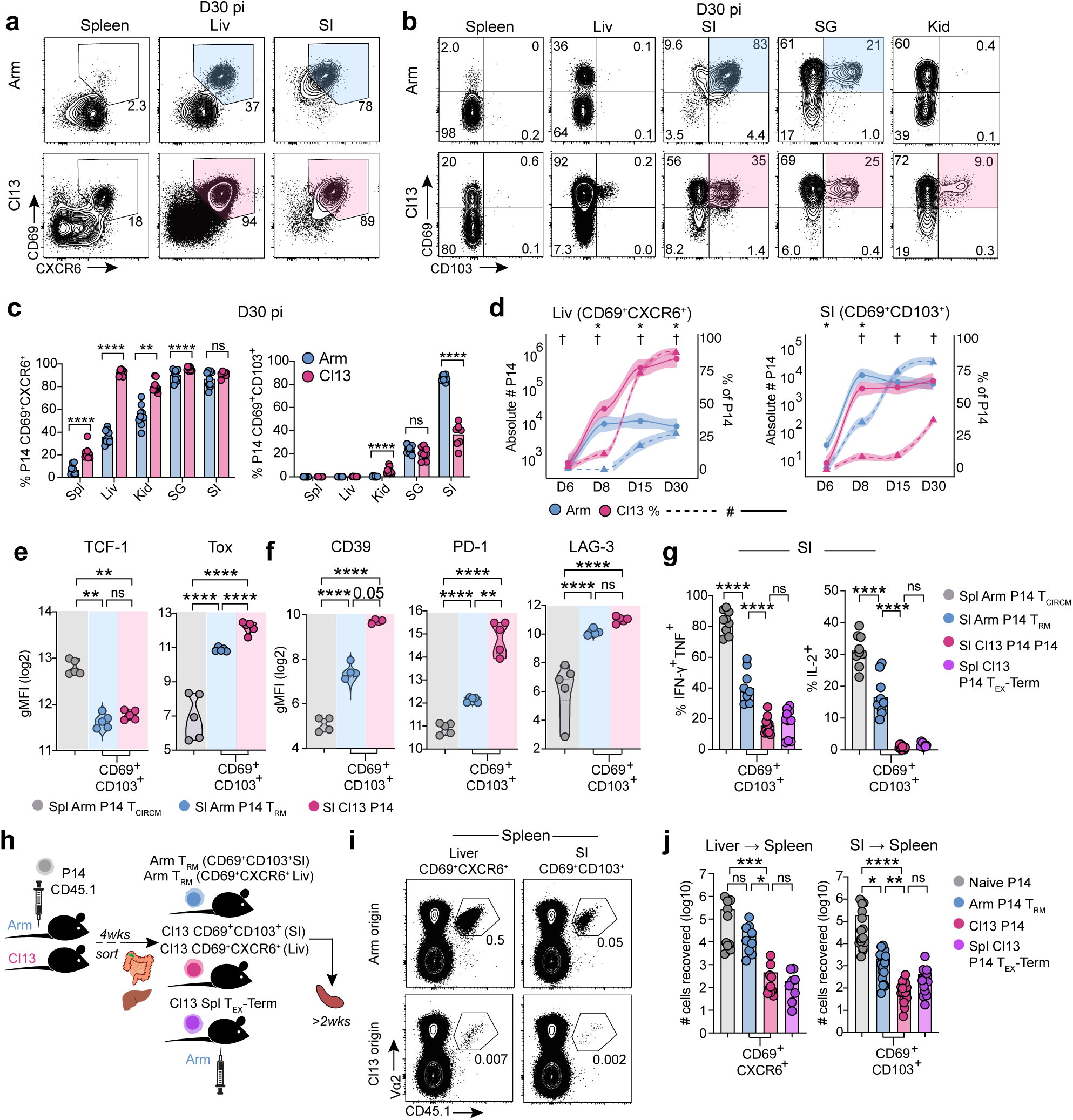
Functionally distinct CD8^+^ T cells expressing residency and exhaustion molecules populate peripheral tissues in acute and chronic infection. **a, b** Expression of CD69 and CXCR6 **(a)** or CD69 and CD103 **(b)** by P14 cells from the indicated tissues 30 days post-infection (dpi) with LCMV Arm or Cl13. Spl; spleen, Liv; liver, SG; salivary gland, SI; small intestine epithelium, Kid; kidney. Shading highlights tissue-localizing cells co-expressing both markers. **c**, Frequency of P14 cells expressing indicated molecules isolated from the indicated tissues 30 dpi with Arm or Cl13. **d**, Frequency (dashed lines, ^†^) or absolute number (solid lines, *) of CD69^+^CXCR6^+^ (Liv) or CD69^+^CD103^+^ (SI) cells at indicated dpi with Arm (blue) or Cl13 (pink). Individual points represent mean; shading represents 95% confidence interval. **e, f**, Geometric Mean Fluorescence Intensity (gMFI) of indicated molecules in T_CIRCM_ P14 cells isolated from the Spl of Arm mice (grey) or in CD69^+^CD103^+^ P14 cells isolated from the SI of Arm (T_RM_, blue) or Cl13 (pink) mice 30-40 dpi. **g**, Cytokine production by P14 T_CIRCM_ cells from the Spl of Arm mice (grey), terminally exhausted P14 cells from the Spl of Cl13 mice (T_EX-TERM_), CD69^+^CD103^+^ P14 T_RM_ cells from the SI of Arm infected mice (blue) or CD69^+^CD103^+^ P14 T cells from the SI of Cl13 infected mice (pink) 30-40 dpi following ex vivo gp_33-44_ peptide stimulation. **h**, Experimental schematic. CD45.2^+^ mice received naïve CD45.1^+^ P14 cells and were then infected with Arm or Cl13. The following populations were sort-purified 4-5 wks pi: CD69^+^CXCR6^+^ (Liv) or CD69^+^CD103^+^ (SI) P14 cells from Arm (T_RM_) or Cl13 infected mice, CD69^+^Ly108^-^T_EX_-_TERM_ cells from the Spl of Cl13 mice, and CD44^lo^ P14 naïve P14 cells from blood. Matched numbers of sorted cells (10-15,000) were adoptively transferred to separate naïve CD45.2^+^ recipients that were then rechallenged with Arm. P14 cells were re-isolated from the Spl of rechallenged mice >2 wks pi. **i, j**, Frequency **(i)** and absolute number **(j)** of total P14 cells from recovered from the Spl of Arm rechallenged recipients. Data are pooled from 2-3 independent experiments (**c, d, g, j**), or representative of 2-3 independent experiments (**a, b, e, f, i**) with n = 4-5 mice **(a-g)** or n = 5-7 **(j-k)** mice per group per experiment. * or ^†^ p < 0.05, ** p < 0.01, *** p < 0.001, **** p < 0.0001 Mann-Whitney test (**c, d**), Two-Way ANOVA **(j)** or Kruskal Wallis Test (**e, f, g**).

T_RM_ cells develop rapidly after clearance of acute viral infection, with the T_RM_ cell pool largely established by 8-14 days post-infection (dpi) and remaining numerically stable thereafter^3,6,23^. We therefore examined whether CD8^+^ T cells co-expressing residency-associated molecules emerged with similar kinetics in tissues during chronic versus acute infection. Although the relative frequencies of CD69^+^CXCR6^+^ P14 cells in the liver or CD69^+^CD103^+^ P14 cells in the SI differed in Arm compared to Cl13 infection, cells expressing residency-associated molecules initially (6-8 dpi) accumulated in similar numbers and were either numerically equivalent or present at higher numbers in tissues from chronically infected mice at later time points (30 dpi) (**Fig 1d** and **Extended Data Fig 1d**). P14 cells co-expressing CD69, CXCR6 and/or CD103 in Cl13-infected tissues also had similar induction of other residency-related molecules compared to Arm T_RM_ cells from matched tissues, including high expression of CD38, CD49a (VLA-1) and the TF Runx3 as well as downregulation of Ly6C and CD62L (**Extended Data Fig 1e**). Thus, CD8^+^ T cells expressing residency-associated molecules used to identify T_RM_ cells after infection resolution also populated peripheral tissues with comparable kinetics during chronic infection.

We next asked whether tissue-localizing P14 cells responding to acute or chronic infection differed in their expression of markers used to define T_EX_ cells and T_EX_ subsets (**Extended Data Fig 1f**). P14 cells expressing residency-associated molecules in Cl13-infected tissues phenotypically mirrored spleen-derived T_EX-TERM_ cells, as they lacked Ly108 and CX3CR1 expression (**Extended Data Fig 1g**), consistent with previous studies^29,51,52^. However, Arm T_RM_ cells also lacked expression of these markers (**Extended Data Fig 1g-i**), and both Arm T_RM_ and Cl13-derived tissue P14 cells similarly downregulated the memory-associated TF TCF1 compared to splenic T_CIRCM_ cells (**Fig 1e**). CD69^+^CD103^+^ SI and SG and CD69^+^CXCR6^+^ liver P14 cells generated after acute infection (Arm T_RM_ cells) had higher expression of the exhaustion-pioneering TF Tox and IRs including PD-1, CD39, LAG-3, TIGIT, and CD101 compared to splenic T_CIRCM_ cells. However, expression of these molecules by Arm T_RM_ cells was lower than for CD69^+^CD103^+^ and CD69^+^CXCR6^+^ P14 cells from chronically infected tissues (**Fig 1e-f** and **Extended Data Fig 1i-k**). Thus, both Arm T_RM_ cells and P14 cells expressing residency-associated molecules in Cl13 infected tissues expressed markers associated with T_EX-TERM_, with the highest expression of these molecules observed in the setting of chronic infection.

These results highlighted phenotypic similarities between T_RM_ cells generated after acute infection and tissue-localized CD69^+^CXCR6^+^CD103^+/–^ T cells in chronic infection, raising the possibility that these cell populations may be developmentally or functionally related. Unlike T_EX_ cells, T_RM_ cells are capable of robust cytokine production^53,54^, can locally expand^8,55^, and can transmigrate out of peripheral tissues following antigen-driven recall^56,57^. We therefore compared the functional capacities of T cells expressing residency-associated molecules in Cl13 infected tissues to T_RM_ cells generated after acutely resolved infection and to T_EX-TERM_ cells in the spleen of chronically infected mice. CD69^+^CD103^+^ Arm T_RM_ cells from epithelial tissues were less polyfunctional than Arm T_CIRCM_ cells (**Fig 1g** and **Extended Data Fig 1l-m**). However, chronically stimulated CD69^+^CD103^+^ P14 cells from the SI and SG and CD69^+^CXCR6^+^ P14 cells from the liver of Cl13 infected mice produced substantially lower levels of IFN-γ, TNF, and IL-2 than Arm T_RM_ cells from matched tissues, mirroring limited cytokine production by splenic Cl13 T_EX-TERM_ cells (**Fig 1g** and **Extended Data Fig 1l-m**).

We next directly compared the plasticity potential of Arm T_RM_ cells to tissue-derived P14 cells expressing residency-associated molecules from chronically infected mice by sorting these populations and adoptively transferring each population to new congenically distinct recipient mice followed by rechallenge with Arm (**Fig 1h** and **Supplementary Material 1a**). Although CD69^+^CD103^+^ T_RM_ cells from the SI were limited in their ability to differentiate into secondary effector cells compared to CD69^+^CXCR6^+^ T_RM_ cells from the liver^21,58,59^, Arm T_RM_ cells from both tissues had a greater potential for expansion compared to their phenotypically matched counterparts from chronically infected mice (**Fig 1i-j**). CD69^+^CD103^+^ P14 cells from the SI and SG of Cl13 infected mice also had a reduced Bcl2/Bim ratio compared to phenotypically matched Arm T_RM_ cells from either tissue, suggesting impaired survival capacity (**Extended Data Fig 1n**). Thus, peripherally localized CD8^+^ T cells with tissue-residency features in chronically infected mice lacked quintessential functional properties characterizing memory T cells, including T_RM_ cells generated after acutely resolved infection. Collectively, these data suggested that antigen-specific CD8^+^ T cells expressing canonical T_RM_-associated molecules including CD69, CXCR6 and CD103 develop in peripheral tissues following either acute or chronic infection and share some overlapping features of exhaustion but exhibit distinct functional potential. Notably, surface markers and TFs typically used to define T_RM_ or T_EX_ cells were insufficient to distinguish dysfunctional CD8^+^ T cells from functional memory T cells in tissues.

### Distinct transcriptional and epigenetic regulation of T_RM_ and tissue-resident exhausted T (TR-T_EX_) cells

These results highlighted a population of CD8^+^ T cells localizing to tissues during chronic viral infection that phenotypically resembled T_RM_ cells generated after antigen clearance but displayed reduced functional capacity. However, it remained unclear whether these chronically stimulated cells expressing residency-associated molecules were a subclass of dysfunctional T_RM_ cell with reduced effector and proliferative potential or a developmentally distinct T cell state. To begin to address this question, we performed single-cell trimodal Transcriptome, Epitope and Accessibility (TEA) sequencing^60^ of total P14 cells from Arm or Cl13 infected mice across multiple tissues (spleen, liver, SI and SG) at 30 dpi (**Fig 2a**). Naïve P14 cells were also analyzed for comparison. This approach allowed us to simultaneously capture protein (via oligo-barcoded antibody derived tags or ADTs), RNA-seq and ATAC-seq profiles in individual P14 cells from matched tissues following Arm and Cl13 infection and deconvolute the impacts of tissue location and duration of antigen stimulation on T cell programming (**Fig 2b**).

**Figure 2.**
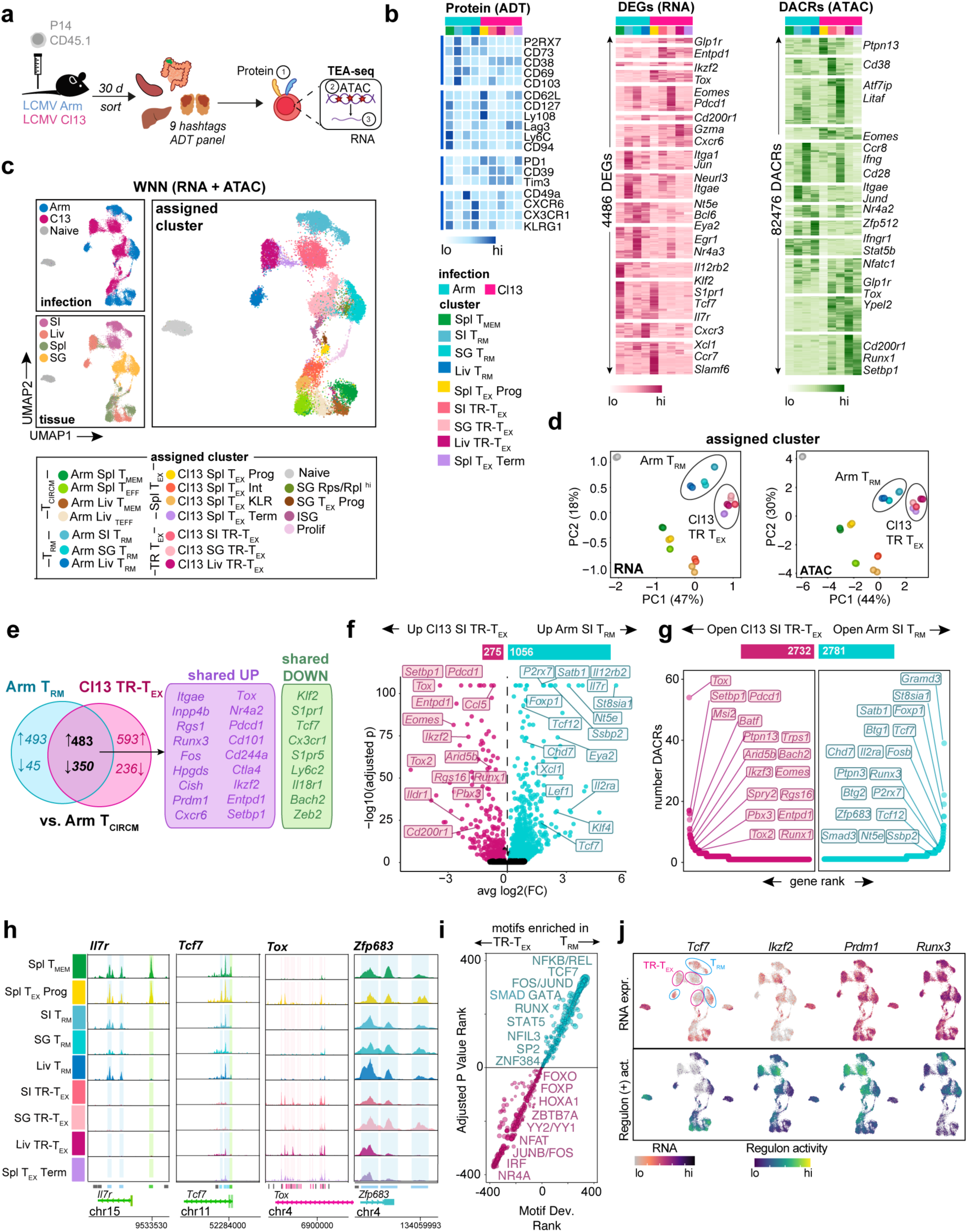
Discrete transcriptional and epigenetic regulation of T_RM_ and T_EX_ cells across tissues. **a,** Experimental schematic for single-cell TEA-seq of tissue P14 T cells. Naïve CD45.1^+^ P14 T cells were adoptively transferred to CD45.2^+^ mice that were then infected with Arm or Cl13. Total P14 cells were sort-purified from Spl, Liv, SI and SG 30 dpi or from naïve P14 blood, stained with Hashtag and Antibody-Derived-Tag (ADT) oligos then pooled and sequenced. **b,** Relative ADT intensity (blue) for all ADTs, RNA expression (pink) for the top 50 unique DEGs, or ATAC accessibility (green) for the top 5000 unique DACRs between indicated clusters. **c,** UMAP of Weighted Nearest Neighbor (WNN) analysis of RNA and ATAC modalities colored by infection (top left panel), tissue origin (bottom left panel) or assigned cluster (right panel, bottom key) based on gene expression and chromatin accessibility directed annotation of unsupervised clusters (see Supplementary Figure 2). **d,** Principal Component Analysis (PCA) of RNA and ATAC diversity between pseudobulk assigned clusters. **e,** Proportion of genes commonly up- or down-regulated by SI Arm T_RM_ (blue) or SI Cl13 TR-T_EX_ (pink) clusters compared to Arm Spl T_CIRCM_ (Arm Spl T_MEM_ and Arm Spl T_EFF_ clusters). **f,** DEGs between SI Arm T_RM_ (blue) and SI Cl13 TR-T_EX_ (pink) clusters. Bar above shows the number of DEGs in pairwise comparison. **g,** Number of DACRs per gene loci between SI Arm T_RM_ (blue) and SI Cl13 TR-T_EX_ (pink) clusters. Bar above shows the number of DACRs in pairwise comparison. **h,** ATAC coverage plots of DACRs in indicated loci for assigned clusters. Peaks called are represented by bars below. Shading on plots or coloring of individual peaks indicates DACRs enriched in Arm Spl T_MEM_ versus Arm T_RM_ (green), Arm T_RM_ versus Cl13 TR-T_EX_ (blue) or Cl13 TR-T_EX_ versus Arm T_RM_ (pink) in at least one tissue. **i,** Motifs enriched in pairwise comparisons of tissue-matched T_RM_ or TR-T_EX_ clusters from each tissue (SI, Liv, SG) rank ordered by adjusted *P* value and motif deviation (Motif Dev.). **k.** RNA expression of indicated TFs (upper panel) and of genes in regulons predicted to be controlled by that TF via network analysis (positive regulon activity). Data are pooled from n = 20-25 mice per group, with significance in pairwise comparisons determined using a Wilcox Rank Sum Test (DEGs) or Logistic Regression Framework Test (DACRs, motifs). **** p < 0.01, two-sided Wilcox test.

Uniform manifold approximation and projection (UMAP) of combined RNA-seq and ATAC-seq data from all samples revealed P14 cells clustered separately based on infection history and tissue location (**Fig 2c** and **Extended Data Fig 2a-b**). Combining RNA and protein expression with chromatin accessibility features resolved known T_EX_ cell heterogeneity in the spleen of Cl13 infected mice including progenitor (Cl13 Spl T_EX-PROG_, *Tcf7*^h*i*^*Slamf6*^hi^), effector-like (Cl13 Spl _TEX-INT_ and Cl13 Spl T_EX-KLR_, *Tcf7*^lo^*Cx3cr1^hi^*) and terminally differentiated (Cl13 Spl T_EX-TERM_; *Cxcr6*^hi^*Entpd1*^hi^) subsets. In contrast, the spleen of Arm-immune mice contained effector-(Arm Spl T_EFF_, *Klrg1*^hi^*Cx3cr1^hi^*) and memory-like (Arm Spl T_MEM_, *Il7r^hi^*) subsets, that we collectively annotated as Spl T_CIRCM_ cells (**Fig 2c** and **Extended Data Fig 2b-2c**). Re-clustering P14 cells separately for each individual organ revealed tissue-derived P14 cells from Arm and Cl13 infected mice were minimally intermixed, with each population forming several heterogeneous clusters that segregated both from splenic cells and from one another (**Extended Data Fig 2d-q**). CD103 (*Itgae)* expression (detected via ADT or RNA expression) was not a major driver of clustering between either Arm or Cl13 P14 cells in the SI and SG (**Extended Data Fig 2g** and l). However, tissue P14 cell clusters from both infections had increased expression of the residency-associated molecules *Cxcr6* and *Cd69* and reduced expression of the tissue egress drivers *Klf2* and *S1pr1* compared to splenic Arm T_CIRCM_ cells (**Extended Data Fig 2f** and k). Given that these RNA features are reliable predictors of long-term T cell tissue-residency in LCMV Arm and Cl13 parabiosis experiments^27^, we annotated *Klf2*^lo^*S1pr1*^lo^Cd69^hi^*Cxcr6^hi^Itgae^+/–^* tissue-derived clusters containing Arm-stimulated P14 cells as ‘Arm T_RM_ cells’ in our TEA-seq dataset (**Fig 2c**).

Principal Component Analysis (PCA) of global transcriptomic and epigenomic profiles revealed liver, SI and SG Arm T_RM_ cells were more closely related to each other than to their tissue-matched counterparts from Cl13 infection or to Arm T_CIRCM_ cells. In contrast, tissue-derived Cl13 P14 cells were positioned away from Arm T_RM_ cells in both transcriptomic and epigenomic PCA space, acquiring a chromatin landscape most similar to that of Spl Cl13 T_EX-TERM_ cells (**Fig 2d**). We therefore annotated clusters of chronically stimulated P14 cells in peripheral tissues as Cl13 tissue-resident T_EX_ (Cl13 TR-T_EX_) cells because these cells expressed tissue residency-associated molecules such as CD69, CXCR6 and CD103 but were molecularly distinct from Arm T_RM_ cells generated following acutely resolved infection.

Nevertheless, comparing either Arm T_RM_ or Cl13 TR-T_EX_ cells to Spl Arm T_CIRCM_ cells revealed many transcriptional similarities between both tissue-derived populations, suggesting expression of a shared set of tissue residency-associated genes linked to tissue location (**Extended Data Fig 3a**). Of the 976 genes upregulated and 395 genes downregulated in SI Arm T_RM_ cells compared to splenic T_CIRCM_ cells, 50% and 90% respectively were also concordantly regulated in SI Cl13 TR-T_EX_ compared to Spl Arm T_CIRCM_ (**Fig 2e**). These shared genes included numerous canonical residency-related molecules known to be upregulated (*Itgae*, *Fos, Jun, Runx3*) or downregulated (*Klf2, S1pr1, Tcf7, Ccr7*) in T_RM_ cells compared to T_CIRCM_ cells, as well as increased expression of several hallmark T cell exhaustion genes (*Tox, Nr4a2, Pdcd1, Setbp1* and others). Moreover, genes previously shown to distinguish T_RM_ from spleen-derived T_EX-TERM_ cells (e.g. *Inpp4b*, *Cish*, *Hpgds*) ^61^ were similarly expressed in Arm T_RM_ and their Cl13 TR-T_EX_ cell counterparts isolated from matched tissues (**Fig 2e**). Indeed, Arm T_RM_ and Cl13 TR-T_EX_ cells isolated from the same organs comparably upregulated many of the same tissue-specific genes (**Extended Data Fig 3a-b**), with tissue-derived TR-T_EX_ cells expressing elevated levels of adhesion molecules and metabolic and stress pathway components compared to splenic T_EX-TERM_ cells (**Extended Data Fig 3c**). Together, these findings highlight common patterns of gene expression adopted by tissue-localized T_RM_ and T_EX_ cells reflective of their local microenvironment. As such, many genes previously selectively associated with T_RM_ cells were similarly expressed in TR-T_EX_ cells isolated from the same tissues, suggesting they were more closely linked to tissue imprinting than to T cell differentiation state.

Therefore, to explore key differences between Arm T_RM_ and Cl13 TR-T_EX_ cell states in a manner that controls for tissue location, we performed pairwise comparisons between these populations from each individual organ. We identified >1000 differentially expressed genes (DEGs) and >2000 differentially accessible chromatin regions (DACRs) between tissue-matched Arm T_RM_ and TR-T_EX_ cells, with the majority of DEGs selectively upregulated in Arm T_RM_ cells (**Fig 2f-g**, **Extended Data Fig 3d-e** and **Extended Data Table 2** and **3**). Similar results were obtained when the analysis was restricted to CD103^+^ Arm T_RM_ and Cl13 TR-T_EX_ cells identified by ADT expression (**Extended Data Fig 3f**). Although expression and accessibility of TCF1 (*Tcf7*) was lower in Arm T_RM_ cells compared to Arm T_CIRCM_ cells^4,62^ (**Fig 1d** and **2e**), Arm T_RM_ cells across tissues had higher gene expression and chromatin accessibility at the *Tcf7* locus as well as at a variety of other canonical memory and stem-related gene loci (including *Il7r, Foxp1, Il12rb2, Satb1, Btg1*) compared to tissue-matched Cl13 TR-T_EX_ (**Fig 2g-h** and **Extended Data Fig 3d**). In addition, Arm T_RM_ cells had higher expression and accessibility at several gene loci previously associated with T_RM_ cell biology (*Xcl1, Smad3, Il2ra*) and at numerous others with uncharacterized roles (*Nt5e*, *Chd7*, *Tcf12*, *St8sia1, Klf4* and others) compared to Cl13 TR-T_EX_ cells (**Fig 2g-h**). Whereas both Arm T_RM_ and Cl13 TR-T_EX_ cells were enriched for expression of exhaustion-related genes including *Tox, Pdcd1* (PD-1) and *Entpd1* (CD39) relative to Arm T_CIRCM_ cells (**Fig 2e**), pairwise comparison of tissue-matched Arm T_RM_ and Cl13 TR-T_EX_ cells confirmed TR-T_EX_ further enriched for expression and accessibility at these shared gene loci (**Fig 2f**-**h****, Extended Data Fig 3d** and **Extended Data Fig 4a**). Arm T_RM_ and Cl13 TR-T_EX_ cells also differed in accessibility at loci encoding regulators of T_RM_ cell commitment^3,4^, including *Runx3*, *Zfp683* (Hobit) and *Prdm1*. For example, Cl13 TR-T_EX_ cells from SI and liver lacked accessibility at sites in the *Zfp683* locus that were open in T_RM_, T_CIRCM,_ and T_EX-PROG_ cells (**Fig 2h** and **Extended Data Fig 4a**).

These results suggested that despite shared expression of some residency- and exhaustion-associated molecules, tissue-matched Arm T_RM_ and Cl13 TR-T_EX_ cells were transcriptionally and epigenetically distinct, with many of the differences between these populations corresponding to memory and exhaustion lineage-associated genes. Indeed, TF motifs with increased accessibility in Arm T_RM_ cells compared to tissue-matched Cl13 TR-T_EX_ cells were highly correlated with those distinguishing Arm T_CIRCM_ from Cl13 Spl T_EX-TERM_ cells (**Extended Data Fig 4b**). Memory-associated motifs in the REL/NFKB and TCF/LEF families were the most significantly enriched in Arm T_RM_ cells compared to Cl13 TR-T_EX_ cells, whereas exhaustion-associated NR4A, NFAT and IRF family motifs were highly enriched in both Cl13 TR-T_EX_ and Cl13 Spl T_EX-TERM_ cells compared to Arm T_RM_ cells (**Fig 2i**). Thus, Arm T_RM_ and TR-T_EX_ cells were characterized by induction of a common tissue-associated transcriptional framework but distinct memory (for T_RM_) or exhaustion (for TR-T_EX_) epigenetic programs.

To explore how gene expression might be controlled in each cell type, we used Pando^63^ TF-gene correlations to infer TF activity in T_RM_ and TR-T_EX_ cells. Differential regulome analysis predicted distinct TFs to be active in Arm T_RM_ versus Cl13 TR-T_EX_ cells from matched tissues, including several shared between Arm T_RM_ and Arm T_CIRCM_ cells (Lef1, TCF1), some selectively enriched in T_RM_ cells compared to other cell states (Tcf12, Klf4) and others preferentially active in Cl13 TR-T_EX_ cells (Ikzf2, Runx1, Nr3c1) (**Fig 2j** and **Extended Data Fig 4c**). The key residency-promoting TFs Blimp1 and Runx3^3,4^ also showed differential predicted activity in Arm T_RM_ compared to Cl13 TR-T_EX_ cells, pointing to potential differences in the coordination of tissue residency-programming in either cell subset (**Fig 2j**). Together, these data suggested that T_RM_ and TR-T_EX_ cells are epigenetically distinct cellular states governed by divergent regulatory circuitry. Although both cell types acquire shared residency-driving transcriptional features linked to tissue location, T_RM_ cells generated after infection resolution have a core memory gene program, whereas TR-T_EX_ cells in chronically infected tissues adopt a canonical exhausted T cell epigenome.

### Shared and distinct transcriptional drivers of T_RM_ and TR-T_EX_ cell programming

The results above suggested that Arm T_RM_ and Cl13 TR-T_EX_ cells were governed by distinct underlying molecular regulation, including potential differences in their use of master-regulator TFs of T cell state specification. Despite differences in chromatin accessibility at *Runx3* and *Prdm1* (Blimp1) loci, both tissue-matched Arm T_RM_ and Cl13 TR-T_EX_ cells from each organ had higher expression of these TFs compared to splenic Arm T_CIRCM_ and Cl13 T_EX-TERM_ cells. Of note, whereas Arm T_RM_ cells in all tissues shared high *Zfp683* (Hobit) expression, only SG Cl13 TR-T_EX_ cells expressed this TF (**Fig 3a**). We therefore asked whether Arm T_RM_ and Cl13 TR-T_EX_ cells differed in their dependence on Blimp1, Hobit, or Runx3 for development or maintenance.

**Figure 3.**
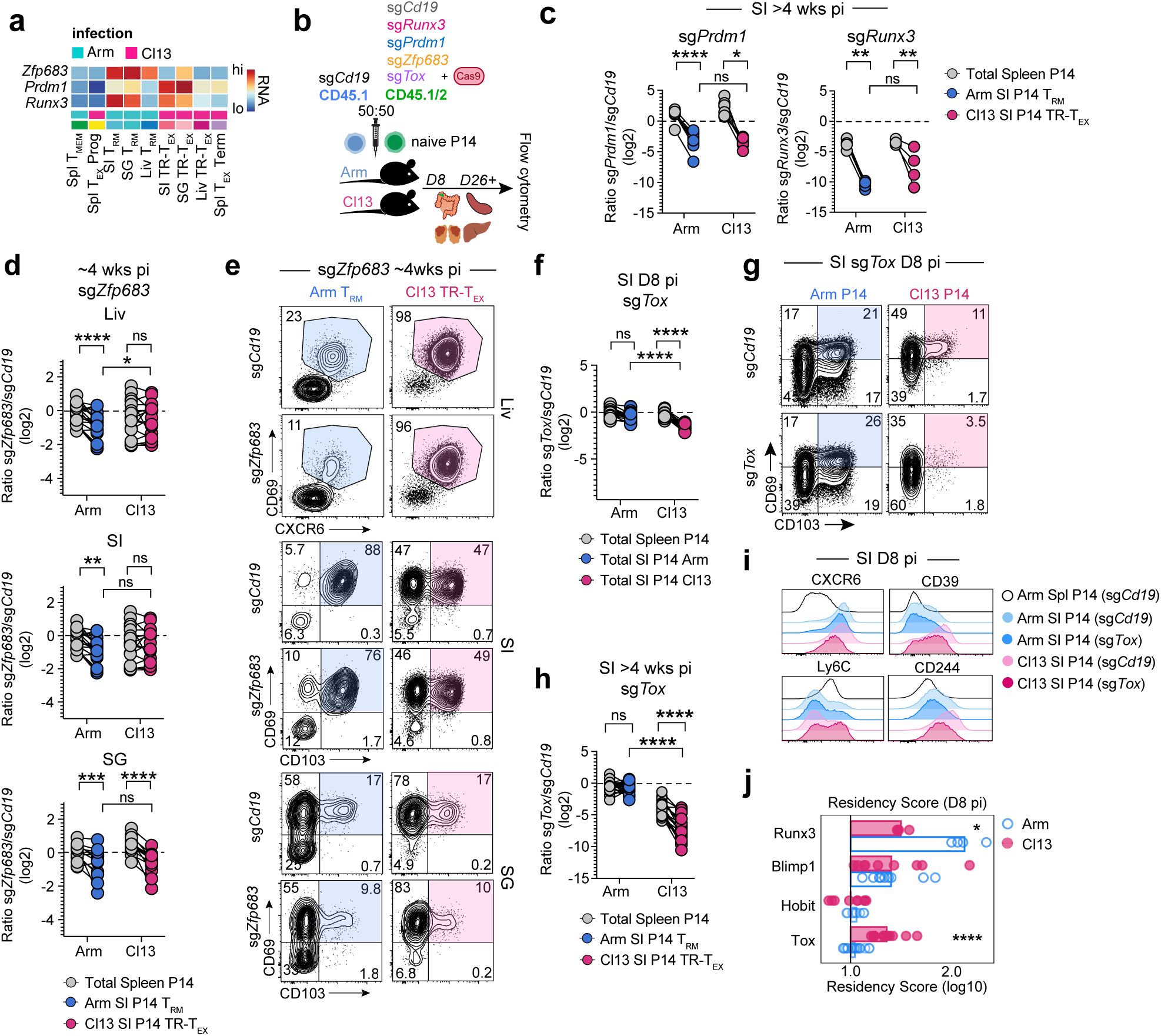
Divergent developmental requirements of T_RM_ and TR-T_EX_ cells during residency programming. **a,** RNA expression of TFs in TEA-seq clusters at 30 dpi. **b,** Experimental schematic for CRISPR-Cas9 TF editing in naïve P14 cells. Naïve CD45.1^+^CD45.1^+^ or CD45.1^+^CD45.2^+^ P14 cells were electroporated with single-guide (sg) RNAs directed towards control (*Cd19*) or target genes with Cas9 protein and adoptively co-transferred at a 50:50 ratio to CD45.2^+^ recipient mice that were then infected with Arm or Cl13. **c,** Ratio of co-transferred SI-derived CD69^+^CD103^+^ Arm T_RM_ (blue), SI-derived CD69^+^CD103^+^ Cl13 TR-T_EX_ (pink) or total Spl-derived (grey) P14 cells edited with *Prdm1* or *Runx3* sgRNAs versus control *Cd19* sgRNAs at 30-37 dpi. **d,** Ratio of co-transferred Arm T_RM_ (blue) or TR-T_EX_ (pink) P14 cells edited with *Zfp683* sgRNAs or control *Cd19* sgRNAs in indicated tissues (SI, SG P14 gated on CD69^+^CD103^+^, Liv P14 gated on CD69^+^CXCR6^+^) compared to total Spl P14 cells (grey) >26dpi. **e,** Co-expression of tissue-residency markers CD69 and CD103 (SI, SG) or CD69 and CXCR6 (Liv) by P14 T cells edited with sg*Cd19* or sg*Zfp683* >26 dpi with Arm (blue) or Cl13 (pink). **f,** Ratio of total co-transferred P14 cells edited with *Tox* sgRNAs or control *Cd19* sgRNAs from the Arm (blue) or Cl13 (pink) infected SI (colored) or spleen (grey) at 8 dpi. **g,** Co-expression of CD69 and CD103 by sg*Tox* or sg*Cd19* edited SI P14 T cells at 8 dpi. **h,** Ratio of co-transferred SI-derived CD69^+^CD103^+^ Arm T_RM_ (blue), SI-derived CD69^+^CD103^+^ Cl13 TR-T_EX_ (pink) or total Spl-derived (grey) P14 cells edited with *Tox* sgRNAs or control *Cd19* sgRNAs >30 dpi. **i,** Expression of indicated molecules in total P14 cells edited with *Tox* or control *Cd19* targeted sgRNAs in SI or Spl at 8-9 dpi. **h,** Normalized fold-change (FC) in residency score (directionally weighted normalized mean expression of CD103, CD69, CD39, CD38, CXCR6, CD49a and Ly6C compared to sg*Cd19* control) in total P14 cells edited with TF-targeting sgRNAs at 8-9 dpi with Arm (blue) or Cl13 (pink). Values indicate extent to which combined expression of markers is regulated by indicated TF. Data are pooled from or representative of 2-3 independent experiments with n = 4-5 (**c**) or n = 4-8 mice (**d-j**) per group per timepoint. * p < 0.05, ** p < 0.01, *** p < 0.001, **** p < 0.0001 paired T test (spleen versus tissue P14s) or two-tailed T test (Arm versus Cl13 tissue P14s) **(c, d, f, h)** or Mann-Whitney Test **(i, j).**

CRISPR-Cas9 electroporation targeting of *Runx3*, *Prdm1* (Blimp1) and *Zfp683* (Hobit) in naïve P14 cells prior to Arm or Cl13 infection (**Fig 3b**) revealed a clear role for Runx3 and Blimp1 in the generation (d8-9 pi) and maintenance (>4wks pi) of both Arm T_RM_ and Cl13 TR-T_EX_ cells across tissues (**Fig 3c** and **Extended Data Fig 5a-e**). Whereas Arm T_RM_ cells from all tissues examined were moderately reduced in frequency and number following CRISPR-Cas9 targeting of Hobit, Cl13 TR-T_EX_ cells in the liver and SI were unaffected (**Fig 3d-e** and **Extended Data Fig 5e**). However, both Cl13 TR-T_EX_ cells and Arm T_RM_ cells in the SG were similarly reduced following Hobit perturbation (**Fig 3d-e**), suggesting that Hobit supports TR-T_EX_ cells in some settings and that dependency on (and expression of) this TF is not an exclusive property of T_RM_ cells. A milder impact of targeting Hobit compared to Blimp1 in Arm T_RM_ cells is consistent with some redundancy between these TFs^4^. Even so, partial reliance on Hobit by SG TR-T_EX_ cells occurred despite elevated *Prdm1* expression by TR-T_EX_ in the SG relative to the SI and liver (**Fig 3a**), suggesting potentially non-redundant activities of Hobit and Blimp1^64^. Selective use of Hobit by SG TR-T_EX_ coincided with uniquely increased accessibility at the *Zfp683* locus in these cells, supporting tissue-specific coordination of TR-T_EX_ cell biology. Together, these data demonstrated that T_RM_ and TR-T_EX_ cells both enlist established regulators of residency programming, with particularly important roles for Blimp1 and Runx3. However, the observation that TR-T_EX_ cells did not require Hobit in several tissues where this TF was required by T_RM_ pointed to potential differences in the transcriptional circuits underlying residency programming in each cell type.

Given that similar residency-associated TFs drove differentiation of both T_RM_ and TR-T_EX_ cells, we next asked whether the exhaustion-associated TF Tox was required for the development and survival of each cell type. Tox is required for formation of circulating and lymphoid T_EX_ but is dispensable for T_CIRCM_ cell generation^15–18^. However, T_RM_ cells have higher expression of Tox than T_CIRCM_ cells (**Fig 1e**). As such, the role of Tox in T_RM_ cells, and their TR-T_EX_ counterparts, remained unclear. CRISPR-Cas9 targeting of *Tox* in naïve P14 cells (**Fig 3b** and **Extended Data Fig 6a**) resulted in a robust reduction in formation of TR-T_EX_ cells in the SI as early as 8 dpi in Cl13 infected mice, even before the major impact of *Tox* deletion on developing splenic T_EX_ cells was apparent (**Fig 3f**-**g** **and Extended Data Fig 6b-c**). In contrast, there was no detectable impact of *Tox* disruption on the development of early T_RM_ precursors in tissues 8d after Arm infection (**Fig 3f-g** and **Extended Data Fig 6b** and c). At later time points (> 4wks pi), the frequency and numbers of mature Arm T_RM_ cells across all tissues examined remained unaffected by *Tox* perturbation, whereas loss of Tox severely compromised generation and durability of Cl13 TR-T_EX_ cells, to either a similar or greater extent than T_EX_ in the spleen (**Fig 3h** and **Extended Data Fig 6d-e**). Similar results were obtained when gating on total CD69^+^ P14 cells in tissues without considering CD103 or CXCR6 co-expression (**Extended Data Fig 6f**). Thus, despite expressing Tox, T_RM_ cells do not require this TF for development or survival. In contrast, TR-T_EX_ cells in all tissues are highly Tox dependent.

These data suggested T_RM_ and TR-T_EX_ cells arise through independent differentiation pathways. Consistent with this idea, *Tox* perturbation selectively altered expression of many residency-associated molecules (CD103, CD69, CXCR6, CD39, CD38, Ly6C, CD244) in developing TR-T_EX_ during Cl13 infection but had little to no impact on the expression of these molecules in developing Arm T_RM_ cells (**Fig 3i** and **Extended Data Fig 6g-h**), despite early expression of Tox in both differentiating T cell populations (**Extended Data Fig 6a**). In contrast, most residency-related molecules were dysregulated in both developing Arm T_RM_ and Cl13 TR-T_EX_ cell precursors following disruption of Runx3 or Blimp1 (**Extended Data Fig 6g-h**). Examining induction of residency-associated molecules together with downregulation of circulatory-associated molecules in a combined ‘residency score’ revealed similar residency programming by Runx3 and Blimp1 in both Cl13 TR-T_EX_ and Arm T_RM_ cells, whereas Tox promoted acquisition of residency features exclusively in differentiating Cl13 TR-T_EX_ cells (**Fig 3j**). Thus, in addition to selectively promoting the survival of TR-T_EX_ cells, Tox played a central role in coordinating residency commitment within developing TR-T_EX_ cells but not in T_RM_ cells. Overall, these data supported the notion that TR-T_EX_ cells are a separate cellular lineage diverging from T_RM_ cells that develop after acutely resolved infection, with TR-T_EX_ cells navigating a distinct Tox-dependent developmental pathway to ultimately converge on a common tissue-resident biology.

### Developmental plasticity allows T_RM_ cells to give rise to T_EX_ cells during chronic antigen stimulation

The data above suggested that T_RM_ and TR-T_EX_ are distinct cell types that emerge through at least partially non-overlapping developmental pathways. To directly test the developmental relatedness of T_RM_ and TR-T_EX_ cells, we next asked whether cells committed to one differentiation state could give rise to the other. First, we asked whether established T_RM_ cells generated after acutely resolved infection could become exhausted when chronically restimulated. We used a sort-rechallenge approach whereby equal numbers of liver or SI Arm T_RM_ cells were sort-purified and adoptively transferred to naïve and congenically distinct recipient mice that were then infected with Cl13 (chronic rechallenge) or Arm (acute rechallenge) (**Fig 4a** and **Supplementary Material 1b**). For comparison, we also adoptively transferred naïve P14 cells and circulating CD127^+^CD62L^+^ central memory CD8^+^ T cells (T_CM_) to separate recipient mice that were also then infected with Arm or Cl13. Donor Arm T_RM_ and T_CM_ cells were analyzed by flow cytometry >3 weeks after Cl13 challenge and compared to: 1) in situ Arm T_RM_ cells that had not been rechallenged, 2) donor Arm T_RM_ and T_CM_ cells rechallenged with acute Arm infection, and 3) naïve P14 cells responding to Cl13, to assess potential differentiation of donor populations into T_EX_ cells during chronic infection. Following either Arm or Cl13 rechallenge, donor Arm T_RM_ cells downregulated expression of residency-associated molecules compared to in situ Arm T_RM_ cells (**Fig 4b**). In response to Arm rechallenge, both donor Arm T_RM_ and T_CM_ cells upregulated effector-associated molecules including KLRG1. In contrast, in response to Cl13 infection, donor Arm T_RM_ and T_CM_ cells robustly upregulated Tox and IRs to higher levels than in situ T_RM_ cells (**Fig 4b**). In addition, donor Arm T_RM_ and T_CM_ cells rechallenged with Cl13 had impaired cytokine production and degranulation compared to both in situ T_RM_ cells and donor cells responding to Arm rechallenge, with their defects in polyfunctionality resembling those seen in splenic T_EX_ cells derived from naïve P14 cells (**Figure 4c** and **Extended Data Fig 7a**). Thus, established T_RM_ cells generated after antigen clearance can become functionally exhausted when chronically restimulated with antigen.

**Figure 4.**
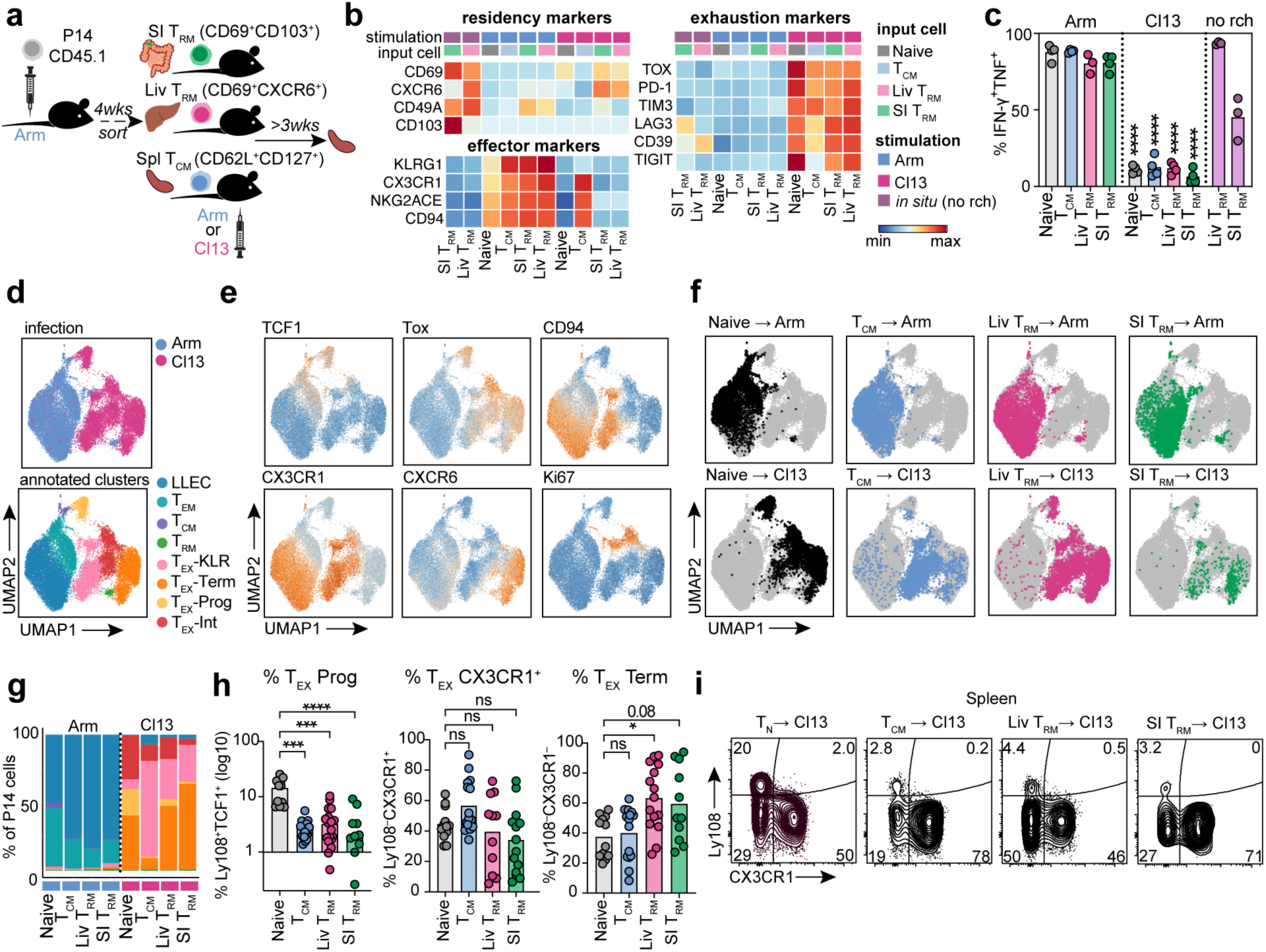
T_RM_ cells differentiate into heterogenous T_EX_ cell subsets during chronic antigen exposure. **a,** Experimental Schematic. Naïve CD45.1^+^ P14 cells were adoptively transferred to naïve CD45.2^+^ mice followed by Arm infection. Four to 6 wks pi, SI or Liv P14 T_RM_ cells (CD69^+^CD103^+^ SI P14 cells or CD69^+^CXCR6^+^CD62L^-^ Liv P14 cells), Spl T_CM_ cells (CD127^+^CD62L^+^ Spl P14 cells) or CD44^lo^ naïve P14 cells were sort-purified and 5000-9000 cells from each population were adoptively transferred to separate naïve recipient mice that were then rechallenged with Arm or Cl13. P14 cells were re-isolated from the spleen 21-28 dpi for flow cytometric analysis. Donor infection-matched SI and Liv T_RM_ cells that were not transferred to new recipients were also isolated for comparison (no rechallenge, no rch). **b,** Relative gMFI of indicated molecules expressed by rechallenged P14 T cells originating from indicated donor origin populations (input cell) following rechallenge (stimulation). **c,** Cytokine production by rechallenged P14 cells originating from indicated donor populations following stimulation with gp_33-41_ peptide in vitro. Statistics indicate pairwise comparisons between Arm and Cl13 rechallenged cells from same input population. **d-e,** UMAPs of rechallenged P14 cells colored by rechallenge virus (infection, **d** upper panel), by phenograph clusters annotated based on surface marker expression (annotated clusters, **d** lower panel) or colored by degree of surface molecule expression **(e). f,** UMAPs highlighting cells derived from indicated input origin populations (colored) following rechallenge with Arm (upper panel) or Cl13 (lower panel). **g-i,** Frequency of cells derived from indicated input origin populations (x axis) giving rise to T_EX_ subsets after Cl13 rechallenge, identified by phenograph cluster (**g**, with same legend key as in **d** lower panel) or by conventional flow cytometry gating (**h, i**). Data are representative of 3 independent experiments **(b, d-f, h)**, 2 independent experiments **(c)** or pooled from 2-3 independent experiments (**g**) with n = 3-5 **(b, d-f**), n = 4-5 **(c)**, n = 4-5 **(g-h**). * p < 0.05, ** p < 0.01, *** p < 0.001, **** p < 0.0001 Mann Whitney Test (**c**), unpaired parametric T test or Kruskal Wallis Test (**g**).

We next asked how T_EX_ cells originating from established T_RM_ cells compared to those derived from naïve T cells. Donor liver Arm T_RM_, SI Arm T_RM_, and T_CM_ cells were all less capable of expansion than naïve P14 cells upon Cl13 infection (**Extended Data Fig 7b**). However, projection of these populations in flow-cytometric UMAP space revealed that regardless of prior identity, different donor cell populations clustered together based on Arm versus Cl13 rechallenge (**Fig 4d-f**). Unsupervised phenograph^65^ clustering revealed that whereas Arm-rechallenged donor Arm T_RM_ and T_CM_ cells predominantly gave rise to long-lived effector cells (LLEC) ^56,57^, Cl13-rechallenged donor Arm T_RM_ and T_CM_ cells primarily populated clusters corresponding to T_EX-KLR_ and T_EX-TERM_ subsets (**Fig 4g** and **Extended Data Fig 7c-d**). Cl13-rechallenged donor Arm T_RM_ and T_CM_ cells also gave rise to T_EX-PROG_ and T_EX-INT_ cells during Cl13 infection, though at a lower frequency than naïve P14 cells (**Fig 4g-i**). Although entirely new clusters did not emerge from donor Arm T_RM_ and T_CM_ compared to naïve P14 cells after Cl13 infection (**Fig 4f-g**), T_EX_ cells originating from both donor Arm T_RM_ or T_CM_ were slightly enriched for expression of residency-associated molecules, including CXCR6 and CD49a, and expressed marginally lower Tox (**Extended Data Fig 7e-g**). These observations suggested that T_EX_ cells derived from previously committed T_RM_ and T_CM_ cells may retain some hallmarks of their prior identity. Nevertheless, these data collectively indicated that persistent antigen exposure could drive differentiation of established T_RM_ cells into T_EX_ cells, highlighting the context-dependent developmental plasticity of T_RM_ cells. Although T_RM_ cells could give rise to multiple conventional T_EX_ subsets including T_EX-PROG_ and T_EX-INT_ cells, T_RM_ cells were skewed towards the generation of more terminally differentiated T_EX-KLR_ and T_EX-TERM_ subsets.

### Committed T_EX_ cells are unable to generate T_RM_ cells following antigen withdrawal

Given that T_RM_ cells could give rise to T_EX_ cells following exposure to persisting antigen, we next asked whether established T_EX_ could generate T_RM_ cells after antigen clearance. We used a similar sort-rechallenge approach, this time isolating committed T_EX-PROG_, T_EX-INT_ and T_EX-_T_ERM_ cells from the spleens of mice infected with Cl13 21d prior and adoptively transferring each subset to congenically distinct naïve recipient mice. These recipients were then infected with Arm, which is cleared by day 8-10 post-infection^24^, allowing us to track the fate of donor T_EX_ cells following antigen withdrawal (**Fig 5a**). We further subdivided donor T_EX-PROG_ cells into previously described lymphoid-resident CD69^+^Ly108^+^ T_EX_-Progenitor-1 (T_EX_-Pr1) and migratory CD69^-^Ly108^+^ T_EX_-Progenitor-2 (T_EX_-Pr2) populations^29^ to examine potential differences in their ability to undergo nonlymphoid residency programming (**Supplementary Material 1c**). Matched numbers of naïve P14 T cells were also adoptively transferred to separate mice as a control for conventional non-migratory T_RM_ cell differentiation following Arm infection.

**Figure 5.**
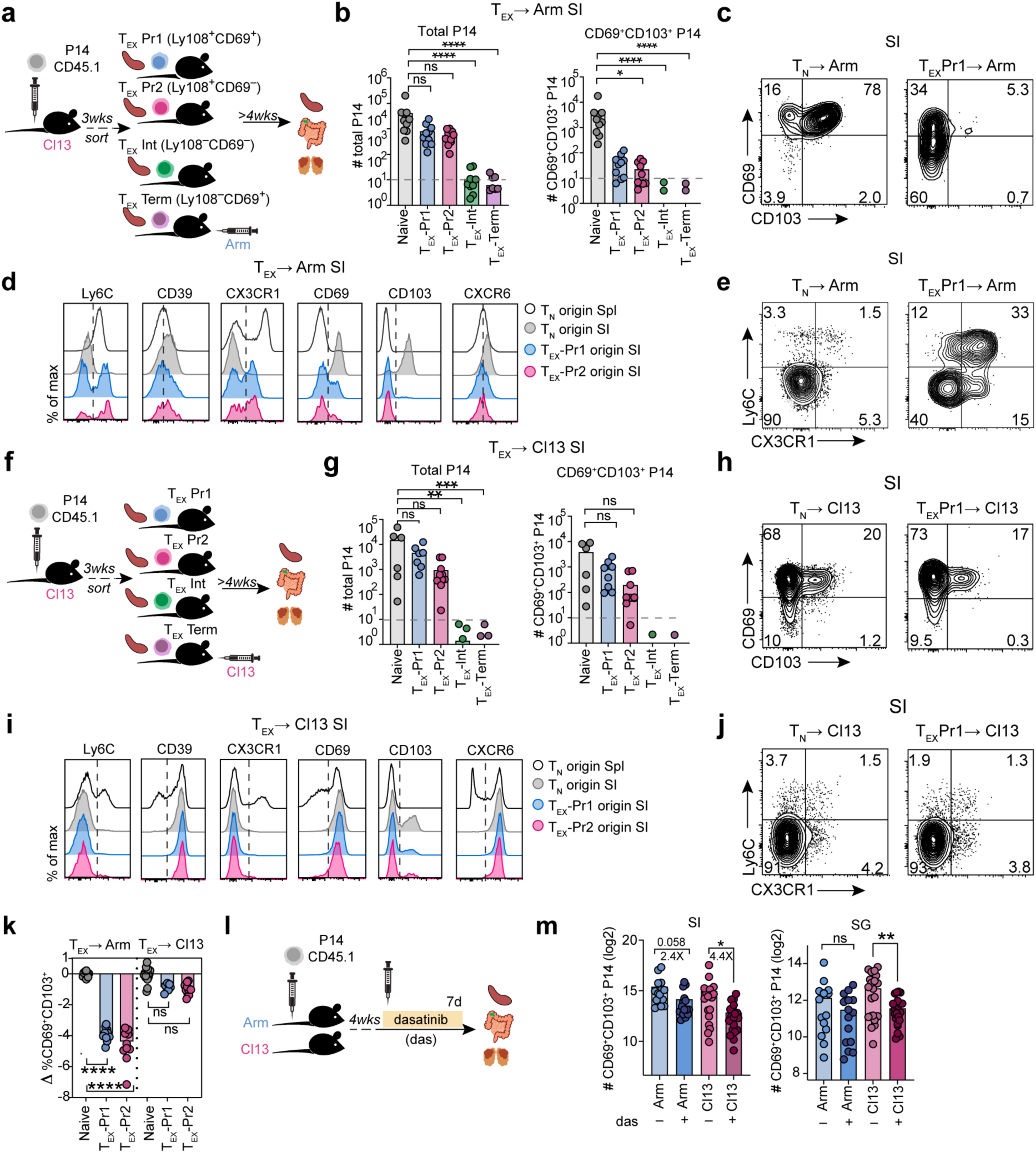
T_EX_ progenitors are unable to generate T_RM_ cells but can give rise to TR-T_EX_ cells. **a,** Experimental Schematic. Naïve CD45.1^+^ P14 cells were adoptively transferred to naïve CD45.2^+^ recipient mice followed by infection with Cl13. Ly108^+^ T_EX-PROG_ (split into CD69^+^ T_EX_-Progenitor 1 [T_EX_-Pr1] or CD69^-^ T_EX_-Progenitor 2 [T_EX_-Pr2] populations), CD69^-^Ly108^-^ T_EX-INT_, CD69^+^Ly108^-^ T_EX-TERM_ from Spl and CD44^lo^ naïve P14 cells were sort-purified 21-23 dpi and 10,000-15,000 cells from each population were adoptively transferred to separate congenic naïve mice that were then rechallenged with Arm . Sorted donor naïve or T_EX_ P14 cells were re-isolated from the Spl and tissues 25-30 days post rechallenge. **b,** Absolute number of total P14 cells (left panel) or CD69^+^CD103^+^ P14 cells (right panel) derived from sorted naïve or T_EX_ cell populations recovered from the SI of Arm rechallenged mice. **c-e,** Expression of indicated surface molecules by sorted naïve or T_EX_ P14 cells rechallenged with Arm and recovered from the SI. **f,** Experimental Schematic. Naïve CD45.1^+^ P14 cells were adoptive transferred to CD45.2^+^ naïve recipient mice followed by infection with Cl13. T_EX-PROG_, T_EX-INT_, T_EX-TERM_ from Spl and CD44^lo^ naïve P14 cells gated as in **a** were sort-purified 21-23 dpi and 5000-10,000 cells from each population were adoptively transferred to separate congenic naïve recipient mice that were then rechallenged with Cl13. **g,** Absolute number of total P14 cells (left panel) or CD69^+^CD103^+^ P14 cells (right panel) recovered from the SI of Cl13 rechallenged mice. **h-j,** Expression of indicated surface molecules by sorted naïve or T_EX_ cell populations rechallenged with Cl13 and recovered from the SI. **k,** Fold change in co-expression of CD69 and CD103 by T_EX_-derived SI P14 cells compared to naïve-derived SI P14 cells after Arm or Cl13 rechallenge. **l,** Naïve CD45.1^+^ P14 T cells were adoptively transferred to CD45.2^+^ recipient mice prior to infection with Arm or Cl13. Beginning 28-30 dpi, mice were treated with dasatinib or vehicle daily for 7d by oral gavage. Shown is the absolute number of CD69^+^CD103^+^ P14 T cells isolated from the SI or SG after treatment. Data are pooled from 2-3 independent experiments with n = 4-5 **(b-e**), n = 2-6 **(g-k**) or n = 5-8 **(l)** mice per group per experiment. * p < 0.05, ** p < 0.01, *** p < 0.001, **** p < 0.0001 unpaired parametric T test (**l**) or Kruskal Wallis Test (**b, g, k**).

Whereas donor T_EX-INT_ and T_EX-TERM_ cells survived poorly in Arm-rechallenged hosts, donor T_EX-PROG_ populated tissues and were present at only slightly reduced numbers compared to naïve P14 cells responding to Arm infection (**Fig 5b** and **Extended Data Fig 7h**). There were no appreciable differences between the expansion capacity or phenotype of Arm-rechallenged T_EX_-Pr1 or T_EX_-Pr2 cells (**Fig 5b** and **Extended Data Fig 7h-k**). Intravascular labeling confirmed that Arm-rechallenged T_EX-PROG_ cells localized to the parenchyma of epithelial tissues, mirroring the distribution of conventional T_RM_ cells derived from naïve P14 cells (**Extended Data Fig 7l-m)**.

Nevertheless, although Arm-rechallenged T_EX-PROG_ cells accumulated in peripheral tissues including the SI, these cells largely failed to upregulate hallmark surface markers defining non-migratory tissue-resident T cells^27^ – such as CD69, CD103, CD49a and CXCR6 – unlike Arm T_RM_ cells derived from naïve P14 cells (**Fig 5c-d** and **Extended Data Fig 7h-j**). Arm-rechallenged T_EX-PROG_ cells also poorly upregulated markers selectively expressed by Arm T_RM_ cells compared to Cl13 TR-T_EX_ cells (see **Fig 2j**), such as CD73 and P2RX7 (**Extended Data Fig 7j**). Instead, tissue-localizing cells derived from rechallenged T_EX-PROG_ expressed markers characteristic of circulating T cells including Ly6C and CX3CR1, and maintained higher expression of Tox, Eomes and Ly108 compared to Arm T_RM_ cells derived from naïve P14 cells (**Fig 5d-e** and **Extended Data Fig 7j** and k). In addition, compared to both Arm T_CIRCM_ and naïve P14-derived Arm T_RM_ cells, Arm-rechallenged T_EX-PROG_ cells in the SI maintained elevated expression of the tissue-homing integrin α4β7 that supports continued CD8^+^ T cell migration^49^ (**Extended Data Fig 7j**), suggesting that tissue-localizing T_EX-PROG_ cells retained trafficking potential. Altogether, these data suggested that despite infiltrating tissues, T_EX-PROG_ cells were unable to acquire canonical tissue-residency features after antigen clearance, and instead upregulated or maintained expression of molecules associated with sustained T cell migration^27,49^. Thus, whereas T_RM_ cells retain developmental flexibility and can convert into T_EX_ cells when antigen persists, committed T_EX_ cells are restrained in their differentiation potential and lose the capacity to acquire a tissue-residency program or generate conventional T_RM_ cells following antigen withdrawal.

### TR-T_EX_ cells arise from committed T_EX_ progenitors during chronic infection

These findings raised questions about the origins of TR-T_EX_ cells that upregulate residency-associated molecules like CD69 and CD103 in the steady state during chronic infection. Specifically, it was unclear whether committed T_EX_ cells could continuously generate TR-T_EX_ cells in a process separate from T_RM_ differentiation or whether T_EX-PROG_ cells lost the ability to acquire certain residency features following commitment to the T_EX_ lineage. We therefore performed a reciprocal experiment in which different T_EX_ subsets were adoptively transferred to congenically distinct mice that were rechallenged with Cl13. We then examined the ability of donor T_EX_ populations to give rise to TR-T_EX_ during chronic antigen stimulation (**Fig 5f**). Donor T_EX_ cells were compared to naïve P14 cells responding to Cl13 infection as a benchmark for typical T_EX_ and TR-T_EX_ cell differentiation. Consistent with findings after Arm rechallenge, donor T_EX-PROG_ cells persisted in the spleen and peripheral tissues following Cl13 infection, whereas donor T_EX-INT_ and T_EX-TERM_ cells were nearly undetectable in recipient mice (**Fig 5g** and **Extended Data Fig 7n**). However, in contrast to findings after Arm rechallenge, progeny of T_EX-PROG_ cells infiltrating Cl13-infected tissues efficiently upregulated residency-associated molecules like CD69, CD103 and CXCR6 and downregulated markers associated with circulating T cells including Ly6C and CX3CR1 in a manner comparable to previously naïve P14 cells responding to Cl13 (**Fig 5h-i** and **Extended Data Fig 7o-q**). Thus, established T_EX-PROG_ readily upregulated residency-associated molecules and generated TR-T_EX_ cells during chronic infection, despite inefficiently upregulating these same features after acute rechallenge (**Fig 5k** and **Extended Data Fig 7r**). Together, these data reveal a selective defect in the ability of T_EX-PROG_ cells to undergo residency programming following antigen withdrawal (after acutely resolved infection) but not in the setting of chronic antigen exposure.

These results suggested that unlike Arm T_RM_ cells^11^, Cl13 TR-T_EX_ cells depend on chronic antigen stimulation. To test this idea, we treated Arm or Cl13 infected mice with the tyrosine kinase inhibitor dasatinib for 1 week starting approximately 30 dpi, to inhibit Lck signaling downstream of the TCR^66,67^ (**Fig 5l**). Whereas Arm T_RM_ cells were minimally impacted by dasatinib treatment, Cl13 TR-T_EX_ cells were substantially reduced in the SI and SG **(Fig 5m** and **Extended Data Fig 7s-t**). Thus, TR-T_EX_ cells were more sensitive to inhibition of ongoing antigen stimulation and TCR signaling for their generation and/or maintenance compared to their Arm T_RM_ counterparts. These data further reinforce the idea that T_RM_ and TR-T_EX_ cells are distinct cell types that originate through separate memory and exhaustion developmental pathways. Whereas established T_RM_ cells retain the differentiation flexibility to become exhausted when exposed to chronic antigen, committed T_EX_ cells require persistent antigen stimulation to undergo tissue residency programming.

### Distinct T_RM_ and TR-T_EX_ cell state specific gene programs

To better understand the molecular underpinnings of T_RM_ and TR-T_EX_ cell identity, we sought to define transcriptional programs that uniquely distinguish these distinct cell states from each other and alternative cell types. To this end, we leveraged our TEA-sequencing dataset to generate a global gene regulatory network (GRN) and compare underlying transcriptional circuits active in Arm T_RM_ or Cl13 TR-T_EX_ cells to those governing other CD8^+^ T cell fates. Global GRN projection in UMAP space (**Fig 6a** and **Extended Data Fig 8a**) and construction of subnetworks specific to each T cell subset (**Fig 6b** and **Extended Data Fig 8b**) identified expected core gene modules engaged in effector (enriched in Arm Spl T_EFF_), memory (enriched in Arm Spl T_MEM_) and spleen-derived exhausted (enriched in Cl13 Spl T_EX-PROG_, Cl13 Spl T_EX-INT_ and Cl13 Spl T_EX-TERM_ clusters) T cell subsets (**Extended Data Fig 8b-d**). We also identified a collection of core residency-related genes (shared residency module) that were engaged in both Arm T_RM_ and Cl13 TR-T_EX_ cells (including *Rgs1, Itgae, Runx3, Itga1* and *Inpp4b*) (**Fig 6a-b** and **Extended Data Fig 8e**).

**Figure 6.**
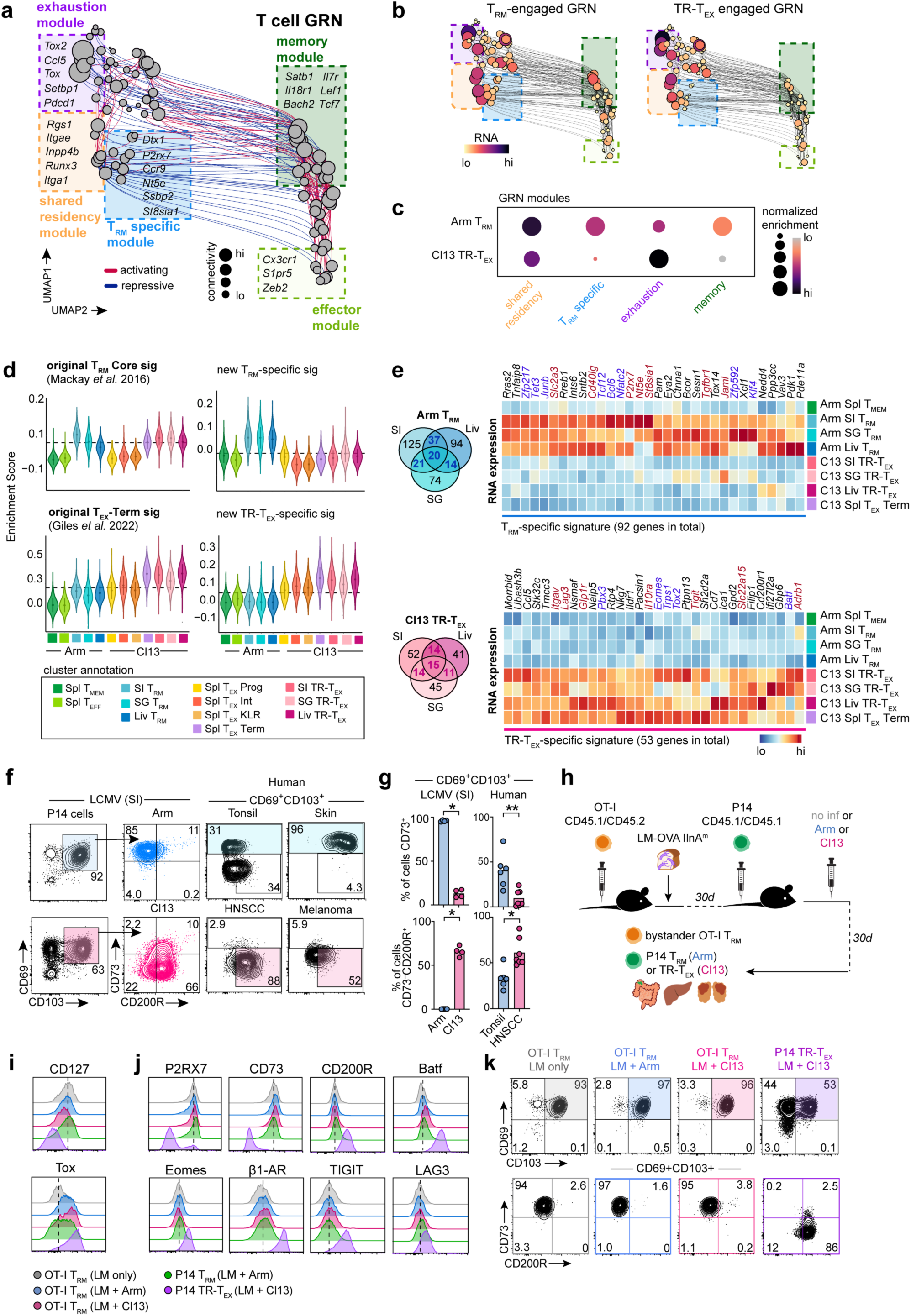
Identification of core transcriptional programs that distinguish T_RM_ from TR-T_EX_ in inflamed tissues. **a,** UMAP of inferred single-cell Gene Regulatory Network (GRN) based on combined TF expression and motif accessibility in all non-naïve P14 cells. Size of nodes (genes) represents number of connections. **b,** Individual UMAP embeddings of genes in GRN in **b** that are actively engaged in either SI T_RM_ (left panel) or SI TR-T_EX_ (right panel) cells. Node size and color scale represent RNA expression. **c,** Relative enrichment for GRN-derived modules highlighted in **a** (highly connected gene clusters) in T_RM_ or TR-T_EX_ across tissues. **d,** Enrichment score for published core T_RM_^4^ and T_EX_-_TERM_^68^ signatures compared to T_RM_-specific or TR-T_EX_-specific signatures in TEA-seq clusters. Dashed line indicates the median score in all plotted cells. **e,** RNA expression of genes selectively upregulated in T_RM_ compared to T_CIRCM_ and TR-T_EX_ cells from matched tissues (T_RM_-specific signature) or selectively upregulated in TR-T_EX_ compared to T_CIRCM_ and T_RM_ from matched tissues (TR-T_EX_-specific signature). Venn diagrams indicate genes shared between T_RM_ or TR-T_EX_ cells across tissues with genes included in each signature colored. Blue text labels: TFs; red text labels: surface proteins. **f,** Expression of CD73 and CD200R by CD69^+^CD103^+^ SI Arm T_RM_ or SI Cl13 TR-T_EX_ cells or by non-naïve CD8^+^CD69^+^CD103^+^KLRG1^-^ T cells from human tonsil or skin, or in head and neck cancer (HNSCC) or melanoma tumors. **g,** Frequency of CD69^+^CD103^+^ P14 T cells (LCMV) or non-naïve CD8^+^CD69^+^CD103^+^KLRG1^-^ T cells (human samples) expressing CD73 and CD200R. **h,** Experimental schematic. Naïve CD45.1^+^CD45.2^+^ OT-I T cells were adoptively transferred to naïve CD45.2^+^ recipient mice that were then infected with the LM-OVA InlA^m^ by oral feeding. Thirty days after LM infection, congenically distinct CD45.1^+^CD45.1^+^ naïve P14 cells were adoptively transferred to the same mice that were then infected with Arm or Cl13 or left uninfected (no inf). Tissues were analyzed 30d after LCMV infection. **i, j** Expression of molecules in SI CD69^+^CD103^+^ OT-I T_RM_ cells without LCMV infection (grey, LM only), after Arm infection (blue) or after Cl13 infection (pink) compared to SI CD69^+^CD103^+^ Arm P14 T_RM_ cells from Arm infected mice (green) or SI CD69^+^CD103^+^ Cl13 P14 TR-T_EX_ cells from LCMV Cl13 infected mice (purple). **k,** Co-expression of CD69 and CD103 by total OT-I T cells or P14 T cells isolated from LM and LCMV infected mice (top panel) and expression of CD73 and CD200R by each indicated CD69^+^CD103^+^ transgenic T cell population. Data are pooled from n = 20-25 mice per group (**a-e**), representative of at least 2 independent experiments with n = 4-5 mice per group or n = 5-7 patient samples per tissue (**f, g**) or representative of 2 independent experiments with n = 4-7 mice per group **(i-k**). * *p* < 0.05, ** *p* < 0.01, **** p < 0.001 Kruskal Wallis Test.

Arm T_RM_ cells were distinguished from other T cell fates including Cl13 TR-T_EX_ cells by combinatorial and selective engagement of: i) a core memory gene module shared with T_CIRCM_ cells (including *Il7r, Satb1*, *Lef1* and *Foxp1*) and ii) a T_RM_-specific gene module comprising several genes (including *Nt5e*, *St8sia1*, *Ssbp2* and *P2rx7*) that were selectively active in Arm T_RM_ cells compared to alternate cell types, including Arm T_CIRCM_ cells and each Cl13 T_EX_ cell subset (**Fig 6b-c** and **Extended Data Fig 8c-d**). Comparing gene expression between Arm T_RM_, Arm Spl T_CIRCM_ and Cl13 T_EX_ cells resolved a similar but expanded foundational set of memory genes expressed by all memory T cells regardless of location (Arm T_RM_ and Arm Spl T_CIRCM_) and lacking in all T_EX_ cells (**Extended Data Fig 8f**). In contrast, TR-T_EX_ cells more strongly engaged the core exhaustion gene module relative to T_RM_ cells without engaging core memory or T_RM_-specific gene modules (**Fig 6b-c** and **Extended Data Fig 8c-d**). In addition, comparative analyses identified a set of exhaustion-related genes increased in both Arm T_RM_ and Cl13 T_EX_ cells compared to Arm T_CIRCM_ cells, but more highly expressed in T_EX_ cells (*Tox*, *Stat3*, *Cd101*, *Nr4a2* and others). These genes were distinct from a separate set that was selectively enriched in all T_EX_ cells including TR-T_EX_ cells, but not expressed by T_RM_ cells (*Eomes, Pbx3, Ubash3b, Gbp6* and others) (**Extended Data Fig 8g**). Thus, although T_RM_ and TR-T_EX_ cells shared some core tissue-residency features, T_RM_ were uniquely distinguished from TR-T_EX_ cells and other cell states by selective expression of a T_RM_-specific gene program. On the other hand, although T_RM_ and TR-T_EX_ cells both engaged some exhaustion-associated genes, other genes were restricted to the T_EX_ lineage and were not upregulated by T_RM_ cells. T_RM_ and TR-T_EX_ cell identities were therefore shaped by distinct gene regulatory networks, with the integrated engagement of cell type-specific as well as shared core gene modules defining each cell type.

Despite these key differences in transcriptional specification, gene signatures typically used to identify T_RM_ or T_EX_ cells conflated these T cell populations: Arm T_RM_ and Cl13 TR-T_EX_ cells from our TEA-Seq dataset were similarly enriched for the core T_RM_ cell gene signature previously reported to distinguish T_RM_ cells from T_CIRCM_ cells^4^, and Arm T_RM_ cells substantially enriched for gene sets previously shown to distinguish T_EX-TERM_ cells from T_CIRCM_ cells^68^ (**Fig 6d**). Thus, existing transcriptional signatures lacked the resolution required to distinguish T_RM_ cells from TR-T_EX_ cells. To address this issue, we derived new transcriptional signatures based on the cell-state specific features of each subset. We identified 92 and 53 genes that were selectively upregulated in either T_RM_ or TR-T_EX_ cells from at least two tissues, respectively, compared to T_CIRCM_ cells (**Figure 6e** and **Extended Data Fig 9a** and **Extended Data Table 4**). Of note, neither cell-state specific signature included *Itgae* or *Zfp683* typically used to annotate T_RM_ in human TIL datasets^32,69^. Instead, the T_RM_ cell signature included several genes in the GRN T_RM_-selective gene module (*Ssbp2, Nt5e, St8sia1, P2rx7, Klf4, Tcf12*) as well as multiple genomic regulators (*Tet3*, *Zfp592*, *Nfatc2*), enzymes (*Pdk1*, *Ptpn3*) and signaling molecules (*Sh2b1*, *Gpr55*) not previously linked to T_RM_ cell regulation. Genes comprising the TR-T_EX_ cell-specific signature included TFs previously associated with T cell dysfunction (*Tox2*, *Eomes* and *Batf*) and T_EX_-lineage specific regulators of cell signaling and stimulation (*Ubash3b*, *Naip5*, *Sh2d2a*), but notably lacked *Tox*, *Entpd1* and other shared exhaustion features (**Figure 6e** and **Extended Data Fig 9a**). Unlike published signatures, the T_RM_ cell-specific gene signature derived here was selectively enriched in Arm T_RM_ cell clusters from our LCMV TEA-seq dataset and in healthy human intestinal tissue^70^, whereas the TR-T_EX_ cell signature preferentially enriched for Cl13 TR-T_EX_ and splenic Cl13 T_EX_-_TERM_ populations and for CD39^+^PD-1^+^ T_EX_-like cells in human blood^71^ (**Figure 6d** and **Extended Data Fig 9b-d**).

Given that these selective T_RM_ and TR-T_EX_ cell signatures included several surface molecules (**Extended Data Figure 9e**), we next asked if these cells could be distinguished using surface proteins. The ectoenzyme CD73 (*Nt5e*) was highly expressed by SI and liver Arm T_RM_ cells but not by Cl13 TR-T_EX_ cells. In contrast, the IR CD200R was selectively upregulated by Cl13 TR-T_EX_ cells but not by Arm T_RM_ cells (**Fig 6f-g** and **Extended Data Fig 9f-g**). Likewise, CD69^+^CD103^+^ CD8^+^ T cells isolated from healthy human tonsil or skin epidermis were enriched for CD73 expression, whereas CD69^+^CD103^+^ CD8^+^ TIL from either head and neck squamous cell carcinoma (HNSCC) or melanoma expressed little CD73 and had high expression of CD200R (**Fig 6g, Extended Data Fig 9g** and **Supplementary Material 1d**). Indeed, CD8^+^CD69^+^CD103^+^CD200R^+^CD73^-^ TIL had higher expression of multiple IRs compared to CD8^+^CD69^+^CD103^+^CD73^+^ cells from healthy tissues, paralleling expression patterns observed for Arm T_RM_ and Cl13 TR-T_EX_ cells after LCMV infection (**Extended Data Fig 9h-i**). Thus, we identified T_RM_ and TR-T_EX_ specific gene signatures and putative surface molecules that could potentially better deconvolute these cell states in human tissues.

Inflammatory cues can influence the biology of memory T cells^72,73^, potentially complicating efforts to distinguish T_RM_ and TR-T_EX_ cells in chronic disease. We therefore next tested whether chronic bystander inflammation impacted expression of molecules distinguishing T_RM_ from TR-T_EX_ cells in a setting where resting T_RM_ cells were exposed to an unrelated chronic infection. In these experiments, we adoptively transferred naïve CD45.1^+^CD45.2^+^ transgenic OT-I T cells specific for Ovalbumin (OVA) to mice that were then infected with recombinant *Listeria monocytogenes* (LM-OVA IlnA^mut^), generating OT-I T_RM_ cells in the SI (CD69^+^CD103^+^) and liver (CXCR6^+^CD69^+^)^74^. Thirty days post-LM-OVA infection, the same mice received naïve CD45.1^+^CD45.1^+^ P14 cells and were either left uninfected (no LCMV), infected with Arm to generate P14 T_RM_ cells or infected with Cl13 to generate P14 TR-T_EX_ cells (**Fig 6h**). This approach allowed us to directly compare the phenotype of resting OT-I T_RM_ cells exposed to acute (Arm) or chronic (Cl13) bystander infection to P14 TR-T_EX_ cells actively responding to chronic infection in the same mice.

OT-I T_RM_ cells maintained similar expression of tissue-residency associated molecules including CD69 and CD103 in Cl13 infected mice with persisting infection compared to Arm infected mice or mice with no LCMV infection (**Extended Data Fig 10a-b**). In contrast, some molecules associated with T_EX_ or T_MEM_ biology were altered by chronic bystander inflammation. For example, OT-I T_RM_ cells exposed to chronic infection downregulated expression of CD127 and upregulated expression of Tox compared to OT-I T_RM_ cells from Arm infected mice or mice with no LCMV infection (**Fig 6i**). However, the expression of key surface molecules and TFs demarcating T_RM_ from TR-T_EX_ cells in the cell-state specific gene signatures derived above (CD73, P2RX7, CD200R, Eomes, Batf and others) remained unchanged in bystander OT-I T_RM_ cells from Cl13 infected mice compared to Arm infected mice or mice with no LCMV infection (**Fig 6j-k** and **Extended Data Fig 10c**). Together, these data indicated that although some conventional memory and exhaustion-associated features were modestly impacted by bystander inflammatory cues, the T_RM_ and TR-T_EX_ cell-specific gene programs defined here remained relatively stable despite bystander chronic inflammation. These data further reinforced the idea that TR-T_EX_ and T_RM_ cells are distinct cell types, with duration of antigen exposure – rather than persistent inflammation – being the key determinant of their differentiation.

### Divergent roles for T_RM_ and TR-T_EX_ cells in chronic disease and immunotherapy

The results above indicated T_RM_ and TR-T_EX_ cells have distinct ontogeny and divergent functional capacities. These observations raised the possibility that each cell type might contribute differently to disease control or immunotherapy responses. TIL expressing CD103 and/or Hobit have been implicated in cancer control and ICB responsiveness^42,43,75^, but the extent to which this correlation reflects the presence of T_RM_ versus TR-T_EX_ cells is unclear. We therefore compared the ability of the cell-state specific T_RM_ or TR-T_EX_ signatures derived above to stratify cancer patient survival in cohorts for which protective associations with TIL displaying T_RM_-associated features had previously been reported^33,43,76,77^. Both T_RM_ and TR-T_EX_ signatures highly correlated with CD8 expression in immunotherapy-naïve patients from The Cancer Genome Atlas (TCGA) skin cutaneous melanoma (SKCM) and METABRIC triple-negative breast cancer (TNBC) cohorts^78^. As a result, either signature predicted survival when considering all patients, possibly reflecting a surrogate for overall T cell infiltration (**Extended Data Fig 10d-f**). However, pre-stratifying patients into CD8^hi^ and CD8^lo^ groups revealed differential association of distinct T_RM_ and TR-T_EX_ cell-state specific signatures with survival in each cohort. Whereas there was a trend towards improved survival in CD8^hi^ melanoma patients with higher TR-T_EX_ cell but not T_RM_ cell signature scores, only the T_RM_ cell signature was highly correlated with survival in CD8^hi^ TNBC patients (**Fig 7a-b** and **Extended Data Fig 10d-e).**

**Figure 7.**
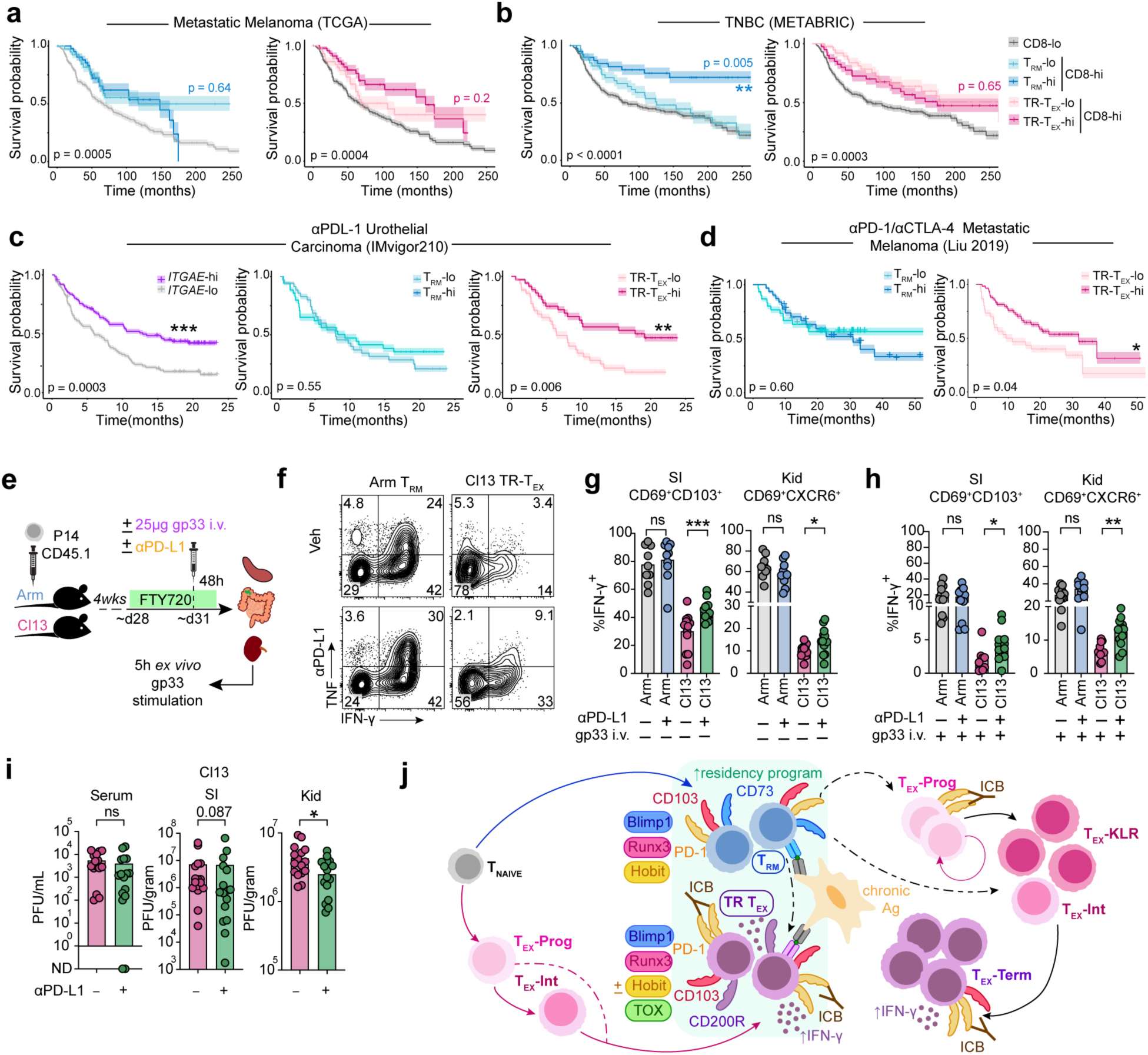
Differential contribution of T_RM_ and TR-T_EX_ cells to disease control. **a,** Kaplan-Meier survival curves for overall survival of metastatic melanoma patients from the TCGA database stratifying by T_RM_ and TR-T_EX_ signatures (^hi^ = top 25%, ^lo^ = bottom 25%) within CD8^hi^ patients compared to CD8^lo^ patients (^hi^ = top 50%, ^lo^ = bottom 50% for *CD8A* expression). **b,** Kaplan-Meier survival curves for overall survival of triple-negative breast cancer (TNBC) patients from METABRIC database stratifying by T_RM_ and TR-T_EX_ signatures (^hi^ = top 25%, ^lo^ = bottom 25%) within CD8^hi^ patients compared to CD8^lo^ patients (^hi^ = top 50%, ^lo^ = bottom 50% for *CD8A* expression). **c,** Kaplan-Meier survival curves for overall survival of urothelial carcinoma patients treated with α-PDL1 from IMvigor210 database stratifying by *ITGAE* expression (left panel; ^hi^ = top 25%, ^lo^ = bottom 25%) or T_RM_ and TR-T_EX_ signatures (^hi^ = top 10%, ^lo^ = bottom 10%). **d,** Kaplan-Meier survival curves for overall survival of melanoma patients treated with α-PD1 and/or α-CTLA4^79^ stratifying by T_RM_ and TR-T_EX_ signatures (^hi^ = top 25%, ^lo^ = bottom 25%). **e,** Experimental Schematic. Naïve CD45.1^+^ P14 cells were adoptively transferred to naïve CD45.2^+^ recipient mice prior to Arm or Cl13 infection. Daily i.p. treatments with FTY720 commenced 4 weeks p.i.. Three days after the first FTY720 treatment mice received one dose of α-PD-L1 or vehicle (PBS) i.p. with or without co-injection of 25µg gp_33-44_ peptide i.v. Tissues were collected 48h later and stimulated ex vivo with gp_33-41_ peptide. **f,** Frequency of CD69^+^CD103^+^ SI P14 T_RM_ (Arm) or TR-T_EX_ (Cl13) cells producing IFN-γ and TNF after α-PD-L1 treatment. **g, h,** Frequency of Arm T_RM_ or Cl13 TR-T_EX_ cells (CD69^+^CD103^+^ in SI or CD69^+^CXCR6^+^ in Kid) producing IFN-γ after α-PD-L1 treatment without **(g)** or with **(h**) i.v. gp_33-41_ peptide treatment. **i,** Viral titers in Cl13 infected FTY720-treated mouse tissues 48h after one dose of α-PD-L1. **j,** Model schematic. T_RM_ and TR-T_EX_ cells derive from divergent differentiation trajectories but acquire a common residency program. T_RM_ cells maintain heightened plasticity potential whereas TR-T_EX_ cells are responsive to PD-1 pathway blockade. Data are pooled from 2-3 independent experiments (**f-i**) with n = 4-8 (**f-h**) or n = 9-10 (**i**) mice per group. Shading on survival curves represents 50% confidence interval. * p < 0.05, ** p < 0.01, *** p < 0.001, **** p < 0.0001 Mann Whitney Test (**g-i**) or Kaplan-Meier estimate (**a-d**).

We then examined a cohort of urothelial carcinoma patients treated with α-PD-L1 therapy (IMvigor210), where an association between *ITGAE* (CD103) expression and improved survival was previously reported^33^ (and **Fig 7c** *left panel*). Whereas the TR-T_EX_ cell-specific signature strongly enriched for improved patient survival following α-PD-L1 treatment, the T_RM_ cell-specific signature was not associated with disease outcome (**Fig 7c**). Similarly, only the TR-T_EX_ cell-specific signature was associated with improved outcome in metastatic melanoma patients treated with α-PD-1 alone or in combination with α-CTLA4^79^, whereas the T_RM_ cell-specific signature showed no predictive association (**Fig 7d**). These associations therefore suggested that although abundance of T_RM_ cells can correlate with improved cancer outcomes, it was TR-T_EX_ cells rather than T_RM_ cells that were preferentially associated with improved disease control following PD-1-directed ICB.

These data raised questions about whether TR-T_EX_ and T_RM_ cells might be differentially modulated by ICB. We therefore asked whether T_RM_ and TR-T_EX_ cells would differ in their response to PD-1 pathway blockade (α-PD-L1) *in vivo* in the LCMV setting. To control for the presence of antigen, we devised a system where Arm T_RM_ cells could be restimulated at the time of PD-1 pathway inhibition via intravenous (i.v.) infusion of cognate gp_33-41_ peptide. We generated populations of T_RM_ and TR-T_EX_ cells by infecting mice with Arm or Cl13, treated these mice with the sphingosine-1-phosphate (S1P) inhibitor FTY720 to restrict T cell migration between blood and tissues^80^, and then administered α-PD-L1 blocking antibodies to Arm or Cl13-infected mice with or without simultaneous co-infusion of gp_33-41_ peptide i.v. (**Fig 7e**).

Delivery of gp_33-41_ peptide to Arm-infected mice triggered PD-1 upregulation in reactivated T_RM_ cells, as well as upregulation of PD-L1 on local antigen-presenting cells in tissues by 48h post-infusion (**Extended Data Fig 10g**), without prompting CD8^+^ T cell infiltration (**Extended Data Fig 10h**). After 48h of α-PD-L1 treatment, there was no change in cytokine production by Arm T_CIRCM_ or Cl13 T_EX_ cells in the spleen (**Extended Data Fig 10i-j**). However, CD69^+^CD103^+^ SI and CD69^+^CXCR6^+^ Kid Cl13 TR-T_EX_ cells from α-PD-L1 treated mice had enhanced IFN-γ production and degranulation compared to control-treated mice, both with and without gp_33-41_ peptide co-infusion (**Fig 7f-h** and **Extended Data Fig 10k-n**). In contrast, cytokine production and degranulation by both resting and peptide-reactivated Arm T_RM_ cells was unaffected by α-PD-L1 treatment (**Fig 7f-h** and **Extended Data Fig 10k-n**). Enhanced cytokine production by Cl13 TR-T_EX_ cells coincided with locally reduced Cl13 viral titers in the kidney but not serum 48h after a single dose of α-PD-L1 compared to control treated mice (**Fig 7i**), suggesting that TR-T_EX_ cells can enhance local viral control following ICB. Thus, the restrained effector functions of TR-T_EX_ cells can be partially enhanced by α-PD-L1 blockade, whereas T_RM_ cell activity is not enhanced by ICB *in vivo*. These findings underscore a fundamental functional divergence between T_RM_ and TR-T_EX_ cells, pointing to distinct roles for each subset in disease control and immunotherapy responses.

## Discussion

CD69^+^CD103^+^CD39^hi^ CD8^+^ T cells that acquire transcriptional features of T_RM_ cells are abundant in autoimmunity, chronic infections or cancer where they often associate with disease outcomes. Despite many reports identifying such cells, their ontogeny has remained unclear. A key question was whether T_RM_-like cells expressing IRs found in tissues in chronic disease settings are a subset of pre-committed T_RM_ cells experiencing dysfunction or whether these cells are a separate T cell state. The data presented here support a model where persistent antigen stimulation induces a population of CD69^+^CD103^+/–^ T_EX_ cells within peripheral tissues that are developmentally and functionally distinct from T_RM_ cells formed after antigen clearance. Despite engaging a common tissue-residency program, T_RM_ and TR-T_EX_ cells have different underlying transcriptional circuitry and arise from divergent developmental pathways. TR-T_EX_ cells acquire an epigenome most similar to splenic T_EX-TERM_ cells and are derived from committed T_EX-PROG_ cells that are unable to form conventional T_RM_ cells following antigen clearance. Moreover, whereas TR-T_EX_ cells depend on the exhaustion-driving TF Tox for both tissue-residency programming and survival, T_RM_ cells generated after infection resolution do not require Tox. T_RM_ and TR-T_EX_ cells were stable differentiation states shaped by antigen clearance or persistence, respectively, with each cell type maintaining its intrinsic identity even after exposure to bystander inflammation. Thus, these findings support a model where T_RM_ and TR-T_EX_ cells are orthogonal cellular states belonging to separate memory and exhaustion CD8^+^ T cell differentiation trajectories, respectively (**Fig 7j**). Such a model has implications for understanding T cell programming, the dynamics of CD8^+^ T cell responses in chronic disease and the therapeutic responsiveness of CD8^+^ T cell subsets.

CD8^+^ T cells in nonlymphoid tissues acquire a set of characteristic transcriptional features including elevated Hobit (*Zfp683*) and CD103 expression, initially described for cells generated after acute-resolving infections^3,4,6^. This transcriptional module is often used to identify and annotate T_RM_ cells in chronic diseases including tumors. However, our data demonstrates that this residency-associated transcriptional framework can also be co-opted by T_EX_ cells in non-lymphoid tissues. These TR-T_EX_ cells activate a foundational exhaustion transcriptional program shared with other T_EX_ cells together with this residency module, whereas T_RM_ cells are characterized by the combinatorial engagement of core memory T cell, common tissue-residency, and T_RM_-cell specific gene modules. Despite both subsets expressing a shared set of residency-associated features, T_RM_ and TR-T_EX_ cells have distinct functional capacities that reflect their underlying memory or exhaustion lineage identity. Memory T cells formed after antigen clearance – including T_RM_ cells – retain the developmental plasticity required to contribute to secondary immune responses. Indeed, not only do T_RM_ cells have the capacity to generate new effector and memory T cells upon acute rechallenge^8,55–57,81^, they can also differentiate into T_EX_ cells following chronic antigen stimulation. In contrast, T_EX_ cells – including those with residency features – are programmed to develop an epigenetic landscape that restricts their fate flexibility and differentiation plasticity. Thus, although both T_RM_ and TR-T_EX_ cells can populate tissues during chronic infection, cancer and other diseases, these are biologically distinct cell types with unique properties that may differentially impact disease trajectories and therapeutic responsiveness.

Although these new data and related studies^82^ suggest that distinguishing between T_RM_ and TR-T_EX_ may be of interest in chronic disease, identifying memory versus exhausted CD8 T cells in tissues has been challenging. Use of individual genes (or previous gene signatures) comprising a common residency module may be useful to identify cells with residency properties, but such signatures were insufficient to distinguish T_RM_ from TR-T_EX_ cells. Our analyses derived new cell-state specific transcriptional signatures and surface markers that can distinguish these cell types with improved resolution. These transcriptional programs were stably maintained during chronic inflammation, indicating that it is now possible to use these signatures to separate T_RM_ from TR-T_EX_ cells in chronic disease settings. Such an approach could be informative in human tissues, where factors such as T cell specificity and the timing or duration of antigen stimulation are often unknown. Our data suggest that T_RM_ cells with memory-like attributes arise in situations where antigen has been completely cleared or compartmentalized in a manner that limits continuous T cell stimulation. Such scenarios could include after resolution of acutely resolved infection, in some settings of latent infections^83,84^, during tumor-immune equilibrium where antigen is available intermittently and T cell stimulation is low^10^, following tumor antigen mutation or loss, or after vaccination. In contrast, sustained antigen stimulation such as occurs in chronically infected tissues or in established tumors drives the formation of T_EX_ cells that combine residency features with a backbone of exhaustion. This combination leads to the development of a different resident CD8^+^ T cell type (TR-T_EX_) with distinct regulatory circuitry that shapes cellular behavior, durability and response to immunotherapy. When the biological properties and origins of T cells in tissues cannot be explicitly defined or tested, broader less prescriptive terms – such as ‘tissue-resident T cells’ – may better capture the idea of tissue-residency biology without inferring memory versus exhaustion properties.

The distinct functional properties of T_RM_ and TR-T_EX_ cells described here may influence their respective contributions to disease control. For example, the transcriptional signatures of T_RM_ or TR-T_EX_ cells differentially correlated with patient survival in distinct cancer settings. Whether T_RM_ or TR-T_EX_ cells can directly inhibit human cancer growth, or whether protective associations between either cell type and disease outcomes reflects more broadly the presence of a productive anti-tumor CD8^+^ T cell response, remains an open question. Studies in preclinical models suggest antigen-specific T_RM_ cells generated prophylactically via infection or vaccination can directly inhibit cancer growth^10,85^. T_RM_ cells may also drive cancer-immune equilibrium and prevent tumor progression^10^. Thus, antigen-specific T_RM_ cells generated after tumor clearance or after a reduction in tumor burden may persist and reactivate to protect against cancer recurrence or metastasis. Likewise, TR-T_EX_ TIL may also be capable of contributing to cancer control, but in different settings. Although T_EX-PROG_ cells are necessary to sustain durable anti-tumor CD8^+^ T cell responses their role is to give rise to downstream T_EX_ subsets such as T_EX-INT_ and/or T_EX-TERM_ that can mediate some tumor cell killing by producing effector cytokines and cytotoxic molecules locally^86–88^. Indeed, T_EX-TERM_ and/or T_EX-INT_ cells found in the TME are capable of slowing tumor growth independently of T_EX_-_PROG_ cells in mouse models^86,88–90^. Thus, both T_RM_ and TR-T_EX_ cells could act through distinct pathways to reduce tumor burden and curtail cancer progression.

Nevertheless, the differences in T_RM_ and TR-T_EX_ functionality observed here have implications for immunotherapy, suggesting distinct approaches could be tailored to effectively harness or modulate each population. Prior work has shown T_EX-PROG_ cells are key targets of PD-1 pathway blockade, as these are the cells that are reinvigorated and expand to give rise to more effector-like T_EX_ and more terminally differentiated T_EX_ cells following ICB^91–93^. One implication of this idea is that associations between TR-T_EX_ cells and beneficial responses to immunotherapy in cancer patients might reflect increased abundance of upstream T_EX-PROG_ cells that respond to ICB, in turn amplifying TR-T_EX_ cell abundance. Our data now add to this understanding of ICB effects on T_EX_ by demonstrating that the effector functions of terminally exhausted TR-T_EX_ cells themselves are also reinvigorated by PD-1 pathway blockade *in vivo*, highlighting a secondary mechanism through which ICB may promote cancer control. Although these TR-T_EX_ cannot expand numerically, the increase in local effector function of these cells elicited following PD-1 pathway blockade was sufficient promote some disease control during chronic infection. These data are consistent with results in tumor explant cultures from human melanoma where ex vivo tumor-resident TIL function can be reinvigorated by PD-1 blockade even in tumor fragments where T_EX-PROG_ were unlikely to be present^94^. Thus, TR-T_EX_ cells have the potential to provide local benefit following PD-1 pathway blockade.

In contrast, although T_RM_ cells generated after antigen clearance express IRs including PD-1, the functions of these cells are not directly enhanced by PD-1 targeted ICB. Instead, T_RM_ cells retain higher baseline functionality and plasticity compared to T_EX_ cells, which may allow these cells to contribute to disease control in part via the generation of multiple T_EX_ subsets following chronic antigen stimulation (**Fig 7j**). Thus, T_RM_ cells might participate more effectively in cancer prevention, cancer vaccination, or adoptive cellular therapy settings that depend on flexible and durable T cell responses, whereas TR-T_EX_ cells may be more amenable to transient intervention using ICB. This latter notion is consistent with our analyses that indicated T_RM_-like TIL associated with responses to ICB across diverse tumor contexts^42,45,76,95,96^ are more closely aligned with TR-T_EX_ cells than to T_RM_ cells generated after antigen clearance. Overall, these new findings suggest a re-evaluation of how to define tissue CD8^+^ T cell states in pathological settings, with implications for understanding how to best leverage CD8^+^ T cell responses to improve treatment of chronic disease.

## Methods

### Mice

Female C57BL/6 recipient mice were purchased from Charles River or Jackson Laboratories and used at 6-8 weeks of age. CD45.1^+^ × P14 (B6.SJL.PtprcaPep3b/BoyJ) and CD45.1^+^CD45.2^+^ × P14 transgenic mice expressing a TCR specific for H2-D^b^-restricted LCMV gp_33-41_, and CD45.1^+^ × OT-I and CD45.1^+^CD45.2^+^ × OT-I transgenic mice expressing a TCR specific for H2-K^b^-restricted OVA_257-264_ were bred in-house at the University of Pennsylvania and backcrossed to either NCI C57BL/6 or Jackson Laboratory C57BL/6J mice and used at 6-18 weeks of age. Mice were maintained in a specific-pathogen-free animal facility at the University of Pennsylvania at ∼20°C with 55% humidity and a light-dark cycle of 12h/12h. All experiments and breeding were performed in accordance with the Institutional Animal Care and Use Committee (IACUC) guidelines for the University of Pennsylvania and complied with ethical US and international animal ethical guidelines.

### Human samples

All human specimens were collected with informed consent and approval from the University of Pennsylvania Institutional Review Board (IRB#844642, IRB#417200 and IRB#808244). Healthy human peripheral blood mononuclear cell (PBMCs, n = 3), skin (n = 5), tonsil (n = 6), melanoma (n = 4) and HNSCC (n = 7) samples were obtained through the University of Pennsylvania. Age, sex and tissue site information can be found in **Extended Data Table 5**.

### Adoptive T cell transfer

500-1000 CD45.1^+^ naïve P14 T cells (in LCMV Arm and LCMV Cl13 experiments) and 10,000 CD45.1^+^ naïve OT-I T cells (in LM-OVA experiments) were isolated from the peripheral blood of donor mice via density gradient centrifugation with Histopaque-1083 (Sigma-Aldrich) and adoptively transferred to recipients in PBS intraveneously (i.v.) through the tail vein or retro-orbitally (r.o.) 1-2 days prior to infection. In sort-rechallenge experiments using only Arm donors, 5000 naïve P14 T cells were adoptively transferred.

### Infections and plaque assays

LCMV Arm and Cl13 were grown in BHK cells (American Type Culture Collection (ATCC), CL-10) and titrated by performing plaque assays on Vero cells (ATCC, CCL-81) as described^97^. Recipient mice were infected intraperitonealy (i.p.) with LCMV Arm (2 × 10^5^ PFU) or i.v. with LCMV Cl13 (4 × 10^6^ PFU) diluted in 1% FBS/RPMI. *Listeria monocytogenes* infection was performed using the recombinant LM-OVA IlnA^mut^ strain expressing OVA and a mutated internalin A protein. Mice were infected with 10^9^ CFU LM via the oral feeding route in bread, as described^74^.

### Mouse tissue processing

For flow cytometric analysis, lymphocytes were isolated from the spleen and lymph nodes by pushing organs through a 70μm nylon filter and treating single-cell suspensions with ACK Lysing Buffer (Thermofisher) for 3 min at room temperature (RT) before washing cells with 5% FBS/RPMI. In experiments requiring cell-sorting (TEA-seq and sort-rechallenge experiments), lymphocytes were isolated from the spleen by pushing organs through a 70μm nylon filter then performing EasySep magnetic negative selection for CD8^+^ T cells using a Stem Cell Technologies kit (cat # 19858) according to the manufacturers’ instructions. Lymphocytes were isolated from the liver by pushing organs through a 70μm nylon filter to create single cell suspensions, resuspending cells in 44% Percoll/RPMI solution and performing density centrifugation for 20 mins at 500G at RT and then incubating cells in ACK Lysing Buffer for 5 mins at RT before washing with 5% FBS/RPMI. Lymphocytes were isolated from the salivary gland, kidney and small intestine epithelium as described^21,74^. Briefly, the salivary gland and kidney were finely chopped in 2% FBS/RPMI containing Collagenase Type III (Worthington, 180,000 U/mL) and incubated for 1 hour in a 37°C water bath. After quenching with 5% FBS/RPMI, digested samples were pushed through a 70μm nylon filter and resuspended in a 44%/70% Percoll density gradient before centrifuging for 20 min at 500G at RT and washing cells with 5% FBS/RPMI. Lymphocytes were isolated from the small intestine epithelium by submerging the small intestine in ice cold CMF solution (10%FBS/Hank’s Balanced Salt Solution (HBSS)/Hepes Bicarbonate Buffer), removing Peyer’s Patches, cutting the intestine open lengthwise, and removing mucus and intestinal content before cutting into ∼0.5inch pieces and placing in ice cold CMF. Intestinal pieces were incubated in CMF containing dithioerythritol (DTE, 0.155mg/mL) for 30 mins at 37°C with shaking at 220rpm. Single cell suspensions were vortexed, filtered through a 70μm nylon filter and washed with 5% FBS/RPMI before being overlaid on a 44%/70% Percoll/RPMI density gradient and centrifuged for 20 min at 500G at 18°C prior to washing cells with 5% FBS/RPMI. To perform *in vivo* intravenous fluorescent antibody labeling, mice were injected with 3µg CD8α-BV650 antibody in PBS i.v. 5 minutes prior to sacrifice as described^98^.

### Human tissue processing

Lymphocyte isolation from blood was performed as described^99^. For lymphocyte isolation from tonsil and HNSCC tumor samples, tissue was finely chopped then incubated at 37°C for 60 minutes in DMEM supplemented with DNase I (Sigma Aldrich, 50mg/mL) and Liberase (Roche, 25mg/mL) then passed through a 70mm cell strainer. For lymphocyte isolation from healthy human skin, subcutaneous fat was removed using scissors and the epidermal layer of skin gently scored using a razor blade before incubating samples overnight in Dispase II/PBS solution (Roche, 2.5mg/mL) at 4°C. Dermal and epidermal fractions were separated, finely chopped and incubated separately in Collagenase Type III solution (Worthington, 180,000U/mL) for 90 min in a 37°C water bath. Following incubation, samples were agitated by mixing with a transfer pipette in 10%FBS/RPMI solution and single cell suspensions filtered through a 70μm nylon filter prior to cryopreservation in 10%DMSO/FBS. Melanoma samples were processed as described^100^. For analysis, cryopreserved samples were thawed at 37°C and washed with 10% FBS/RPMI prior to staining.

### Flow cytometry

Single cell suspensions were stained with amine-reactive Zombie dyes (Biolegend) diluted in PBS for 10 mins at RT before staining with surface antibodies (see **Extended Data Table 6**) diluted in a 1:1 mixture of Brilliant Stain Buffer (BD) and 2%FBS/PBS for 60 mins on ice. Samples were then fixed and permeabilized with an eBioscience Foxp3 Staining Kit (Thermofisher Scientific) according to the manufacturers’ instructions before staining with intracellular antibodies for 60-90 mins at RT. To determine absolute cell counts, SPHERO Blank Calibration Particles (6-6.4µm, BD) were added to each sample before acquisition on BD Symphony A5 or Cytek Aurora flow cytometers. Samples were analyzed in Flowjo v10 or in OMIQ. For mean fluorescence intensity calculations in Fig 1 and Extended Data Fig 1, Arm and Cl13 P14 cells from tissue or spleen were stained separately with anti-CD45.1-Biotin antibodies, barcoded with unique streptavidin tags then mixed prior to cell surface staining to control for staining intensity variability between tissue single-cell suspensions. P14 cell ratios in CRISPR-Cas9 experiments were normalized back to the exact input ratio of naïve cells injected at the time of adoptive cotransfer. Combined residency scores were determined by summing the fold-change in MFI of markers typically upregulated in T_RM_ cells (CD69, CD103, CD49a, CXCR6, CD38, CD39) in control sg*Cd19* T cells compared to test guide perturbed T cells, subtracting the fold-change in MFI Ly6C and normalizing back to the average score in the sg*Cd19* versus sg*Cd19* control group.

### In vitro stimulation of CD8^+^ T cells

For *in vitro* restimulation experiments, mice were injected with 50μg of anti-mouse ARTC2 nanobody (Treg protector, Biolegend, clone s+16a) r.o. 10-30 mins prior to euthanasia to limit P2RX7-mediated cell death. Following tissue processing, cells were resuspended in cRPMI (10%FBS/RPMI containing 1% Penn/Step, 10mM HEPES, 1% MEM Nonessential Amino Acids, 1% L-glutamine, 1mM Sodium Pyruvate, 0.05mM β-mercaptoethanol), CD107a AF647 (Biolegend), GolgiStop (1:250, BD, 554724), GolgiPlug (1:500, BD, 555029) and 2 × 10^−4^ µg/mL gp_33-41_ peptide (KAVYNFATM) and incubated at 37°C for 5 hours. Cells were washed, stained with surface antibodies and fixed and permeabilized with a BD Biosciences Cytofix/Cytoperm kit according to the manufacturers’ instructions prior to intracellular antibody staining.

### Flow cytometric sorting and viral rechallenge

For sort-retransfer and TEA-seq experiments, mice were injected with 50μg of anti-mouse ARTC2 nanobody (Treg protector, Biolegend, clone s+16a) r.o. 10-30 mins prior to euthanasia to limit P2RX7-mediated cell death. Following tissue processing, for sort-retransfer experiments cells were resuspended in PBS containing amine-reactive Zombie dyes (Biolegend) and surface antibodies and incubated on ice for 30 mins before being resuspended in 2%FBS/PBS containing 2mM EDTA. For TEA-sequencing experiments, cells were first stained with surface antibodies on ice then resuspended in 2% FBS/PBS containing 7-aminoactinomycin D (7AAD) for discrimination of live cells. Cells were sorted into 50% FBS/RPMI on a BD FACS Aria III using a 100µm nozzle with the sorting chamber maintained at 4°C. For sort-retransfer experiments, cells were washed twice with PBS prior to adoptive transfer to recipient mice via i.v. injection. Mice were either rechallenged with LCMV Arm i.p. 1 hour later or with LCMV Cl13 i.v. 12 hours later.

### Transcriptomic, Epitope and Accessibility (TEA) sequencing

TEA-sequencing was performed using a modified version of the standard 10x Genomics Chromium Single Cell Multiome ATAC + Gene Expression Kit protocol, as described^60^. Briefly, live CD8β^+^CD44^hi^CD45.1^+^Vα2^+^ P14 cells from each sample (pooled organs from 20 Arm-infected mice or 20 Cl13-infected mice) were sorted into separate FBS-coated FACS tubes containing 50%RPMI/FBS and washed once with Cell Staining Buffer (Biolegend) before staining samples with individual TotalSeq-A Hashtag Oligo (HTO) antibodies and a custom pool of TotalSeq-A Antibody Derived Tag (ADT) antibodies (Biolegend; see **Extended Data Table 6**). Samples were washed with Cell Staining Buffer (850G, 7 mins, 4°C) before resuspending cells in 2%BSA/DPBS and pooling cells. Cells were resuspended in custom TEA-seq Wash Buffer (20mM Tris-HCl, 150mM NaCl, 2mM MgCl_2_ and 1% BSA in H_2_O) containing RNase Inhibitor (1U/mL, Roche) and 0.01% Digitonin (Thermofisher) and incubated for 5 minutes on ice to permeabilize cell and nuclear membranes. Cells were topped with TEA-seq Wash Buffer and spun for 7 mins at 850G at 4°C before washing with TEA-seq Tagmentation Buffer (20mM Tris-HCl, 150mM NaCl, 2mM MgCl_2_ in H_2_O). 30,000 cells per reaction were removed and subjected to transposition and GEM generation as described in the 10x Chromium Single Cell Multiome ATAC + Gene Expression Kit. Library preparation was performed as described^60^, with the following customizations. Indexing of ADT and HTO libraries was performed using Biolegend-recommended Illumina D70x (i7 long; HTO) or RNA PCR Index (RPIx short; ADT) adapter sequences (IDT), with three additional rounds of amplification performed on purified HTO/ADT libraries using P5/P7 primers specific for generic Illumina adapter sequences (IDT) prior to sequencing (see **Extended Data Table 7**). Libraries were assessed using D1000 or D5000 High Sensitivity DNA ScreenTape (Agilent) on an Agilent Tapestation and quantified using a KAPA Illumina Library Quantification Kit (KK4824, Roche) prior to sequencing on S1 and S4 Flow Cells (Illumina) using a NovaSeq 6000 Instrument (Illumina).

### Single-cell RNA and ATAC sequencing processing and quality control

BCL files were demultiplexed using cellranger-atac-2.0.0 for ATAC-seq data, bcl2fastq for HTO and ADT data and cellranger-arc for RNA-seq data. Alignment and quantification of reads was performed using cellranger_arc (v2.0.2) for RNA-seq and ATAC-seq data and in Barcounter (v1.0) for HTO and ADT data. Analysis was then performed in R (v4.2) using ArchR (v1.0.2), Seurat (v5.1.0) and Signac (v1.13) using custom R scripts. Quantified HTO and ADT reads were combined with gene expression reads derived from cellranger to create Seurat objects. Cells were filtered out based on the following metrics: less than 500 RNA features, less than 2000 ATAC counts, less than 1500 ATAC features, a nucleosome signal greater than 5, a Transcription Site Enrichment (TSS) enrichment score greater than 4, more than 30% mitochondrial reads, and outliers with high numbers of HTO reads. HTOs were normalized using the Seurat CLR method and manually demultiplexed in using the multimode package (v1.5) by inspecting the distribution of each HTO signal against all others profiled. Negative droplets and doublets identified during HTO calling were removed. ATAC peaks were then called in ArchR using all cells that had passed quality control in Seurat and Signac. Briefly, additional QC was first performed by executing AddDoubletScores() then removing doublets with a filter ratio of 0.5 and filtering additional cells with a TSS Enrichment Score less than 10. Clustering based on ATAC fragments was performed in ArchR using the function addIterativeLSI() with the default parameters except for use of 25,000 variable features and a resolution of 2 with 2 iterations and then executing the addClusters() function with a resolution of 3.4. Pseudobulk replicates were created using the function addGroupCoverages() with a minimum cell cutoff of 30, maximum cell cutoff of 774 and the default parameters. Peaks were then called on clusters using macs2 and the ArchR function addReproduciblePeakSet() using the default parameters. This ArchR-derived peak set was then added to a Seurat object containing filtered cells merged from both wells using FeatureMatrix() and CreateChromatinAssay() and peaks were annotated against the mm10 genome using EnsDb.Mmusculus.v79 (v2.99) and the ClosestFeature() function. Barcounter derived ADT counts from filtered cells were processed and normalized using dsb (v1.0.4) using the default parameters and isotype control antibodies (see **Extended Data Table 6**) then added to the Seurat object.

### Single-cell RNA and ATAC sequencing analysis

Following quality control and peak calling, combined RNA and ATAC analysis was performed in Seurat (v5.1.0) and Signac (v1.13) on 1) the entire object containing all samples (23,338 cells) and 2) individual subsetted objects containing cells from each tissue and spleen in both infections. For RNA normalization and integration, SCTransform (SCT) v2 was performed on each well individually using 3500 variable features with regression of mitochondrial read proportions. Supervised reciprocal PCA (RPCA) integration of wells was performed on SCTransformed RNA using a k.anchor of 20, an anchor matrix disallowing matches between inconsistent HTOs and by excluding a subset of genes shown to be modulated by heat treatment during sample digestion^101^, heatshock genes (beginning with ‘Hsp’ and ‘Dnaj’) and ribosomal-related and mitochondrial genes (beginning with ‘Rps’, ‘Rpl’ or ‘mt’) from the AnchorFeatures. For ATAC normalization and integration, standard processing was performed using FindTopFeatures(), RunTFIDF() and RunSVD() and integration performed using PCA dimensions 2:50 and FindIntegrationAnchors() by subsetting the anchor matrix to disallow matches between inconsistent HTOs then performing IntegrateEmbeddings() using the default settings. Weighted Nearest Neighbor (WNN) analysis was performed using FindMultimodalNeighbors() using the integrated RNA (SCT dimensions 1:30) and ATAC (dimensions 2:50) reductions followed by RunUMAP(). Clustering on WNN within each object was then performed with the following resolutions: 2.3 (whole object), 0.7 (SI and spleen), 0.6 (SG and spleen) and 1.2 (liver and spleen). Clusters were either annotated within the whole object (splenic and naïve P14 cells) or within individual organ subsetted objects, with annotations from the latter then projected back onto the whole object for tissue-derived P14 cells. Principal Component Analysis was performed in edgeR (v4.2.1) and DEseq2 (v1.44) by pseudobulking each cluster according 10x lane origin. CD103-ADT^+^ cells were annotated as cells with normalized expression of CD103-ADT >1 from each tissue. Most RNA-seq and ATAC-seq based clusters comprised cells from a single tissue and infection origin, but clusters defined by proliferation (Prolif) interferon-stimulated genes (ISG) or by high expression of Rps/Rpl stress-related genes (SG Rps/Rpl^hi^) contained a mixture of cells from acutely and chronically infected mice and so were excluded from downstream analysis.

Differentially expressed genes (DEGs) were calculated in Seurat using the SCT assay and functions FindAllMakers() and FindMarkers() for pairwise comparisons using a Wilcoxon Test and log_2_ fold-change cutoff of 0.125 and adjusted *P* value of less than 0.05 with a min.pct of 0.05. Gene signature enrichment was determined using the function AddModuleScore(), except for analysis of bulk RNA-seq human PBMC samples in Extended Data Figure 11b, where single-sample gene set enrichment analysis (ssGSEA) was used (GSVA_1.50.5) to calculate enrichment scores. Cliff’s delta values to assess effect size of enrichment were calculated using the effsize package. The established core T_RM_ cell signature^4^ was defined as all genes upregulated in T_RM_ cells across the SI, skin and liver. The T_EX_-_TERM_ signature^68^ was defined as the top 300 genes enriched in T_EX_-_TERM_ cells (by average log_2_ fold-change) isolated 30 dpi with Cl13 compared to splenic T cells isolated from Arm infected mice. DACRs were calculated in Signac using the functions FindAllMarkers() and FindMarkers() for pairwise comparisons using the LR test and a log_2_ fold-change cutoff of 0.125 and adjusted *P* value of less than 0.05 with a min.pct of 0.01, including the number of counts as a latent variable. Differentially motif enrichment (motif deviation) was calculated in Seurat using the JASPAR2024 motif set (for all vertebrates, extracted from the JASPAR2024 package v0.99.6 and added as a custom pfm) and the function RunChromvar() with BSgenome.Mmusculus.UCSC.mm10, using FindMarkers() for pairwise comparisons using the LR test and a log_2_ fold-change cutoff of 0.125 and adjusted *P* value of less than 0.05 with a min.pct of 0.05, including the number of counts as a latent variable. All heatmaps were generated using pheatmap (v1.0.12) by calculating average feature expression for each cluster using the AverageExpression() function in Seurat. Venn Diagrams and upset plots were generated using VennDetail (v1.20). Feature Plots and Joint Density Plots were generated in Seurat or with scCustomize (v2.1). Genome coverage tracks were generated using the Signac functions CoveragePlot(), PeakPlot(), TilePlot() and AnnotationPlot(). Peak-gene linkages for coverage tracks were calculated in Seurat using the default parameters. Adjusted *P* values < 5^−150^ were mutated to 5^−150^ for display in volcano plots to permit visualization.

Gene regulatory network (GRN) analysis was performed on all non-naïve cells in Pando^63^ (v1.1.1) using the top 3500 variable SCTransform normalized RNA features excluding heat-modulated, heatshock and Rsp/Rpl genes (as above) and using a union of selected candidate regulatory chromatin regions including conserved regulatory elements from ENCODE^102^, phastCons elements conserved across vertebrates^103^ (Vert 35 El) and additional domains of regulatory chromatin (DORCs) identified through peak-gene correlation for all P14 cells performed in FigR^104^ (v0.1.0). Motif identification was performed using the JASPAR2024^105^ motif set (for all vertebrates) and TFs identified using TFDB^106^ and the genome BSgenome.Mmusculus.UCSC.mm10. The GRN was inferred using the function infer_grn() using the xgb method with the peak to gene method Signac, a TF correlation threshold of 0.05 and extending 10^5^ bases up and downstream of the transcription start site. GRN UMAP visualization was performed in Pando and included all genes inferred in the GRN. Cell-specific gene regulatory networks were generated using tidygraph (v1.3.1) using the Pando coefficient matrix, UMAP coordinates from the global GRN and by aggregating RNA expression determined by SCTransform in Seurat for each cluster. Regulatory modules were identified using the function find_modules() with a *P* threshold of 0.05, R^2^ threshold of 0.05 and an nvar_threshold of 6. Regulons were extracted from the Pando SeuratPlus object using the function NetworkModules(), filtered to modules containing more than 7 genes and regulon activity was determined using the AddModuleScore() function in Seurat and assigned to cells using the CreateAssayObject() function. Differential regulon activity was calculated using FindMarkers() in Seurat using a Wilcoxon Test and log_2_ fold-change cutoff of 0.125 and adjusted *P* value of less than 0.05.

### CRISPR-Cas9 electroporation of naïve CD8^+^ T cells

Electroporation of naïve CD8^+^ transgenic T cells was performed as described^107^. Briefly, CD45.1^+^ and CD45.1^+^CD45.2^+^ P14 T cells were isolated from the spleen or LN and subjected to EasySep magnetic CD8^+^ T cell enrichment (StemCell Technologies, kit 19858) according to the manufacturers’ instructions. RNP complexes were generated by incubating 0.3nmol sgRNA (2 guides per target obtained from IDT Technologies, see **Extended Data Table 7**) with 0.6µl Cas9 protein (IDT, 1081059) for 10 mins at RT. 2-10 × 10^6^ CD8^+^ T cells were resuspended in supplemented P3 Buffer (Lonza P3 Primary Cell 4D-Nucleofector X electroporation kit, Lonza, V4XP-3032), mixed with RNP complexes and electroporated in a Lonza 4D-Nucleofector TM 4 Core Unit (Lonza, AAF-1002B) using the program DN100. Cells were quenched with cRPMI and rested for 10 mins at 37°C. 1000 congenically distinct control-targeted and test-targeted cells were then mixed at a 1:1 ratio and adoptively co-transferred i.v. to CD45.2^+^ recipient mice that were infected 48h post-cell infusion. Exact co-transfer ratios were recorded and used to normalize ratios of test to sg*Cd19* control cells observed in each organ back to the starting input ratio for analysis.

### *In vivo* immune modulator, antibody and gp33 peptide administration

For Dasatinib treatment, mice were delivered 50mg/kg Dasatinib (MedChem Express) in 0.9%saline/10%DMSO/20%Sulfobutylether-β-Cyclodextrin (MedChem Express) solution daily for 7 days by oral gavage. For FTY720 treatment, mice were injected i.p. with 1mg/kg Fingolimod (Cayman Chemical) in 2%(2-hydroxypropyl)-beta-Cyclodextrin/PBS (Sigma Aldrich). For α-PD-L1 treatment, mice were injected once i.p. with 200ug of α-PD-L1 antibody (10F.9G2, BioXCell, BE0101) diluted in PBS and analyzed 48 hours later. For i.v. gp_33-41_ peptide administration, mice were injected i.v. with 25µg gp_33-41_ peptide diluted in 10% DMSO/PBS. In all cases, control mice received the vehicle solution only.

### Construction of cell state-specific gene signatures

The T_RM_ cell-specific signature was derived by taking the union of genes significantly upregulated in T_RM_ cells from at least two tissues compared to TR-T_EX_ cells from matched tissue sites and compared to T_CIRCM_ cells (Arm Spl T_MEM_ and Arm Spl T_EFF_ clusters) from the spleen with a log_2_ fold-change threshold of 0.5 and adjusted *P* value cutoff of 0.05, excluding genes that were also upregulated in TR-T_EX_ cells compared to T_CIRCM_ cells. The TR-T_EX_ cell-specific signature was derived by taking the union of genes significantly upregulated in TR-T_EX_ cells from at least two tissues compared to T_RM_ cells from matched tissue sites and compared to T_CIRCM_ cells from the spleen with a log_2_ fold-change threshold of 0.5 and adjusted *P* value cutoff of 0.05, excluding genes that were also upregulated in T_RM_ cells compared to T_CIRCM_ cells.

### Survival probability analysis in tumor patient cohort

Clinical metadata and gene expression count files were obtained from cBioPortal (TCGA SKCM and METABRIC datasets), the R package IMvigor210CoreBiologies (v1.0.1, IMvigor210 dataset) or from Liu *et al.* 2019^79^ (α-PD-L1/α-CTLA4 metastatic melanoma dataset). The METABRIC dataset was pre-filtered to select for triple negative breast cancer (TNBC) patients. Kaplan Meier survival analysis was performed in R using the packages survminer (v0.4.9), survival (v3.7.0) and GSVA (v1.48.3). Where indicated, samples were divided into the top and bottom 50% (CD8^hi^ and CD8^lo^) based on ranked expression of the gene *CD8A*, before the CD8^hi^ fraction was further divided into signature^hi^ (top 10 or 25% as indicated), signature^med^ (middle 50 or 80% as indicated) or signature^lo^ (bottom 10 or 25% as indicated) fractions based on ranked scores for each signature. In other analyses, all samples were split into signature^hi^, signature^med^ or signature^lo^ fractions without *CD8A* pre-stratification. Where relevant, the IMvigor210 cohort was stratified based on clinical response using RECIST criteria. Signature score correlation analyses with *CD8A* expression were performed using the Pearson correlation method in R.

### Statistical analysis

Statistical analysis was performed in Graphpad Prism (v10.2) or in R (v4.2) using a Mann Whitney Test, Wilcoxon Paired Sum Rank Test, Two Way ANOVA or Kaplan Meier Test where indicated in the figure legends and Methods. All statistical tests performed were two-sided tests. Where parametric tests were used, a Kolmogorov-Smirnov test was used to confirm normal distribution of samples. In all other cases, non-parametric tests were used. Except where indicated, individual data points are indicated with symbols and bars on plots indicate the mean. For box plots, the center line indicates the median, the box limits represent the upper 75^th^ and lower 25^th^ percentile and interquartile range and whiskers extend to 1.5x the interquartile range. Samples with fewer than 10 P14 cells were excluded from flow cytometric frequency calculations. Samples for which no or low titers of LCMV Cl13 could be detected in the serum or kidney >15 dpi were excluded from chronically infected groups in rechallenge experiments and as sort donors for rechallenge experiments and TEA-sequencing experiments. Mice were allocated to groups randomly or by quantifying LCMV Cl13 titers in serum by plaque assay and evenly distributing mice between groups based on viral titers.

## Data availability

Multimodal ADT, HTO, RNA and ATAC raw sequencing data generated in this study will be deposited in the National Center for Biotechnology Information Gene Expression Omnibus prior to publication. Raw differential gene expression, differential accessibility data and unique gene signatures generated in this study are available as Extended Data Tables. All other raw and processed data are available from the authors upon reasonable request. Publicly available datasets used for analysis in this manuscript are available at the following accession numbers: GSE70813, GSE179613, GSE199565, or via gutcellatlas.org.

## Acknowledgments

We thank all members of the Wherry Laboratory for helpful discussions and critical analysis of this manuscript. We thank the Penn Cytomics and Cell Sorting Shared Resource Laboratory and Children’s Hospital of Philadelphia Flow Cytometry Core for providing technical support and instrumentation. We thank the Penn Dermatology Skin Biology and Diseases Resource-based Center (SBDRC) for providing human skin samples. We thank Brian Sheridan for providing the IlnA^mut^ strain of LM-OVA.

## Author Contributions

S.L.P. and E.J.W. conceived the study and designed experiments. S.L.P., M.M.P, V.A., M.M., M.S., D.M., L.T., D.R., N.D., T.C., M.K., Y.J.H., M.A.C., V.F., W.K., S.F.N, A.E.B., J.E.W, M.T. and J.R.G. carried out experiments. C.T.B., E.P., Y.L., K.R., R.M.B, E.R.T., D.B., A.C.H, C.T.E and A.D. acquired and provided samples for investigation. S.L.P., M.M.P., S.M., R.R.G. and J.R.G analyzed data. S.L.P. and M.M.P. prepared visualizations. S.L.P. and E.J.W. wrote the manuscript.

## Extended Data Figure Legends

**Extended Data Figure 1.**
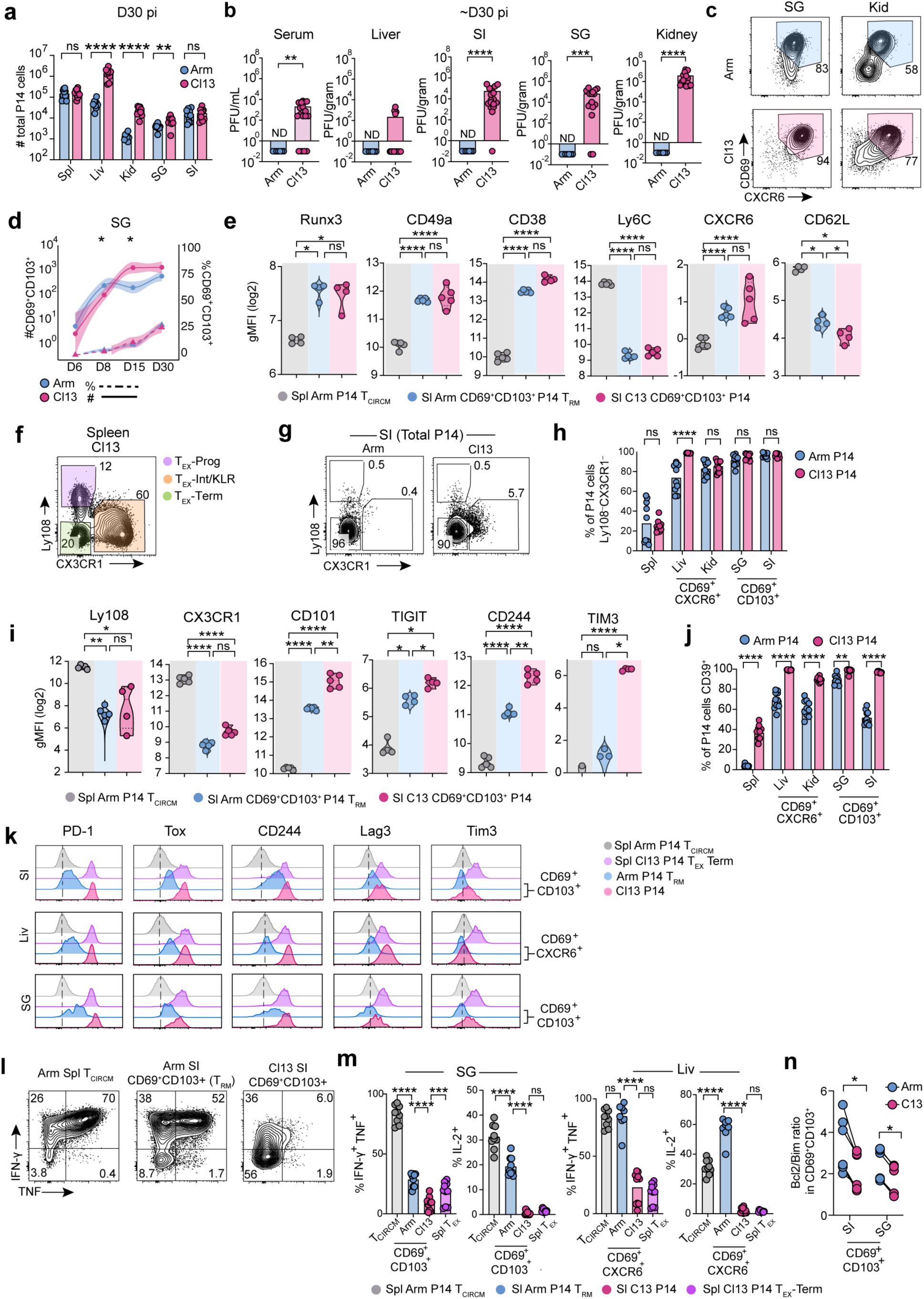
Phenotype and function of tissue-infiltrating CD8^+^ T cells during acute and chronic infection. **a,** Absolute number of Vα2^+^CD45.1^+^ P14 T cells isolated from spleen (Spl), liver (Liv), kidney (Kid), salivary gland (SG) or small intestine epithelium (SI) 30 dpi with acute LCMV Arm or chronic LCMV Cl13 infection. **b,** Viral titers detected by plaque assay in serum or indicated tissues 28-32 dpi. **c,** Expression of CD69 and CXCR6 by P14 cells in indicated tissues 30 dpi with Arm or Cl13. **d,** Frequency (dashed lines, ^†^) or absolute number (solid lines, *) of CD69^+^CD103^+^ cells in SG at indicated dpi with Arm (blue) or Cl13 (pink). Individual data points represent mean; shading represents 95% confidence interval. **e,** Geometric mean-fluorescence intensity (gMFI) of residency-associated molecules by CD69^+^CD103^+^ P14 cells isolated 30 dpi from the SI of Arm (T_RM_) or Cl13 infected mice or T_CIRCM_ cells from Arm Spl. **f,g,** Expression of Ly108 and CX3CR1 by P14 T cells isolated from Spl (**f**) or SI (**g**) 30 dpi. T_EX_-Prog; T_EX_ Progenitor, T_EX_-Int; T_EX_-Intermediate, T_EX_-KLR; T_EX_-Killer cell lectin-like receptor, T_EX_-Term; T_EX_ Terminal. **h,** Frequency of Ly108^-^CX3CR1^-^ cells within total P14 cells (Spl), CD69^+^CXCR6^+^ P14 cells (Liv or Kid) or CD69^+^CD103^+^ P14 cells (SG, SI) 30d post-infection with Arm or Cl13. **i,** Geometric mean-fluorescence intensity (gMFI) of exhaustion-associated molecules by CD69^+^CD103^+^ P14 cells isolated 30 dpi from the SI of Arm (T_RM_) or Cl13 infected mice or T_CIRCM_ cells from Arm Spl. **j,** Expression of CD39 by P14 cells with indicated surface phenotype 30 dpi with Arm or Cl13 across tissues. **k,** Expression of indicated surface markers by CD69^+^CD103^+^ (SI, SG) or CD69^+^CXCR6^+^ (Liv) P14 T_RM_ cells (Arm, blue) or P14 T cells (Cl13, pink) compared to Arm Spl T_CIRCM_ cells (grey) and Cl13 Spl T_EX_-Term cells (purple) 30-40 dpi. Dashed line indicates average expression in Arm T_CIRCM_ P14 cells. **l,** Cytokine production by T_CIRCM_ P14 cells from Arm Spl or by CD69^+^CD103^+^ T_RM_ cells from Arm SI or CD69^+^CD103^+^ P14 cells from Cl13 SI 30-40 dpi following *in vitro* gp_33-44_ peptide stimulation. **m,** Cytokine production by P14 T_CIRCM_ cells isolated from the Spl of Arm infected mice (grey), CD69^+^CXCR6^+^ terminally exhausted P14 cells from the Spl of Cl13 infected mice (T_EX_-_TERM_), or by CD69^+^CD103^+^ (SG) or CD69^+^CXCR6^+^ (Liv) P14 cells from Arm (T_RM_, blue) or Cl13 (pink) infected mice 30-40 dpi following *in vitro* gp_33-44_ peptide stimulation. **n,** Ratio of Bim and Bcl2 gMFI in CD69^+^CD103^+^ P14 T cells isolated from the SI or SG of Arm (T_RM_, blue) or Cl13 (pink) infected mice 15-25 dpi. Data are pooled from 2-3 independent experiments (**a, b, d, h, j, m, n**), or representative of 2-3 independent experiments (**c, e, f, g, i, k, l**) with n = 4-5 mice **(a-m)** or n = 3 mice per group per experiment (**n**). * or ^†^ p < 0.05, ** p < 0.01, *** p < 0.001, **** p < 0.0001 Mann-Whitney test (**a, c, d, e, h, i, j, m**) or Wilcoxon signed-rank test (**n**).

**Extended Data Figure 2.**
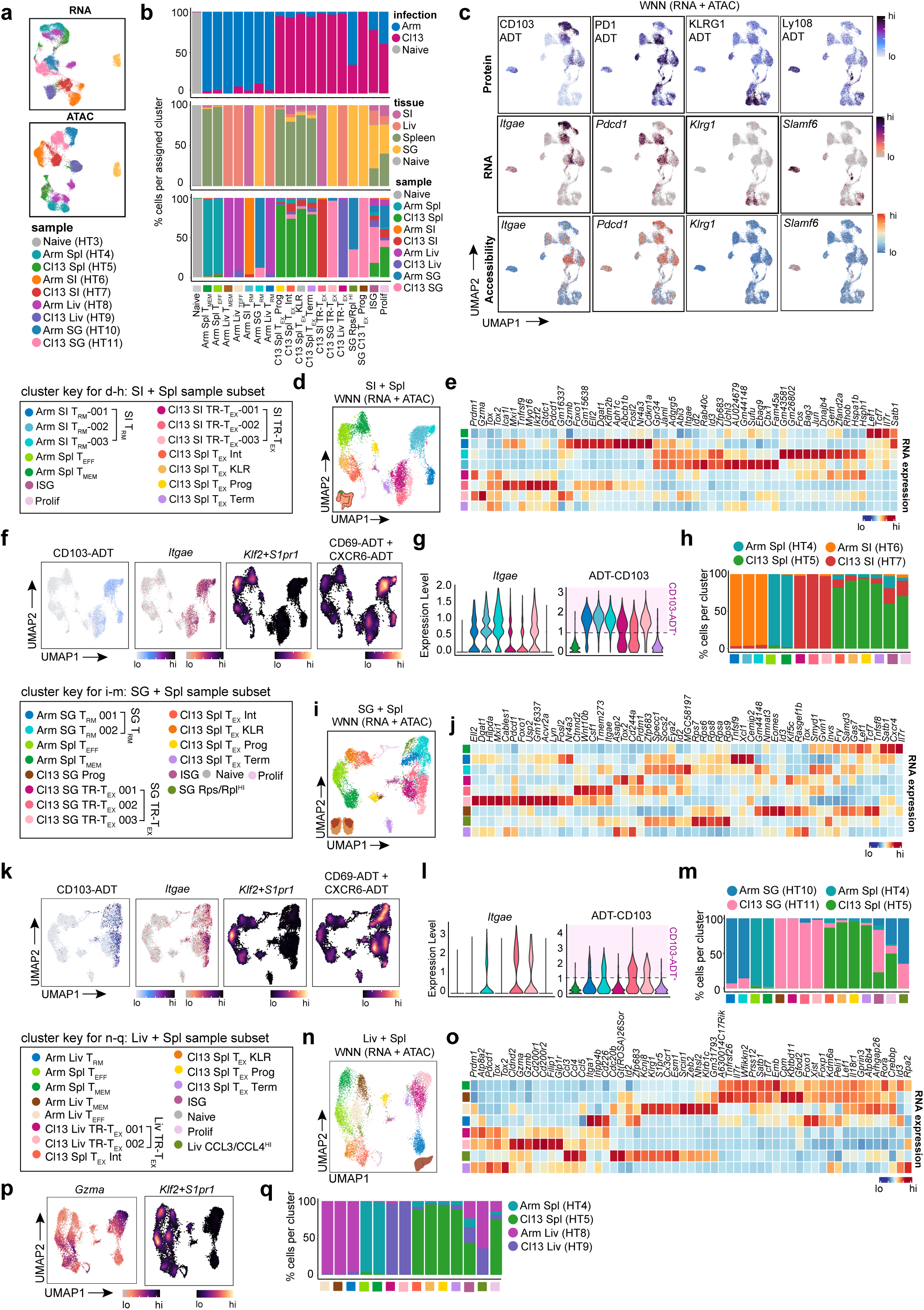
Resolution of T cell states across tissues in acute and chronic infection. **a,** Projection of all P14 cells analyzed by TEA-seq in UMAP space based on RNA expression only (upper panel) or chromatin only (lower panel) colored by sample (Hashtag; HT). **b,** Distribution of P14 cells assigned to each annotated cluster derived from each infection (upper panel), tissue (middle panel) or sorted sample (lower panel). **c,** Weighted-nearest neighbor (WNN) UMAP based on combined RNA and ATAC expression in P14 cells colored by ADT (protein, top row), RNA expression (middle row) or by gene activity (bottom row, chromatin accessibility) for indicated markers. **d,** Re-clustered WNN UMAP based on combined gene expression and chromatin in P14 cells from SI and Spl samples only, colored by annotated Seurat subcluster (cluster key d-h). **e** RNA expression of top 10 marker genes and selected key genes by each indicated SI-derived subcluster and Spl-derived cluster. **f,g** Expression of CD103 protein (ADT) and RNA (*Itgae*), joint RNA expression of *Klf2* and *S1pr1* or joint CD69 and CXCR6 protein (ADT) expression in SI and Spl derived P14 cells. Purple shaded area in **g** indicates cells annotated as CD103-ADT^+^. **h,** Distribution of cells from SI and Spl assigned to each subcluster. **i,** Re-clustered WNN UMAP based on combined gene expression and chromatin in P14 cells from SG and Spl samples only, colored by annotated Seurat subcluster (cluster key i-m). **j,** RNA expression of top 10 marker genes and selected key genes by each indicated SG-derived subcluster and Spl-derived cluster. **k, l,** Expression of CD103 protein (ADT) and RNA (*Itgae*), joint RNA expression of *Klf2* and *S1pr1* or joint CD69 and CXCR6 protein (ADT) expression in SI and Spl derived P14 cells. Purple shaded area in **l** indicates cells annotated as CD103-ADT^+^. **m,** Distribution of cells from SG and Spl assigned to each subcluster. **n,** Re-clustered WNN UMAP based on combined gene expression and chromatin in P14 cells from Liv and Spl samples only, colored by annotated Seurat subcluster (cluster key n-q). **o,** RNA expression of top 10 marker genes and selected key genes by each indicated Liv subcluster and Spl-derived cluster. **p,** Expression of *Gzma* RNA or joint RNA expression of *Klf2* and *S1pr1* in Liv and Spl derived P14 cells. **q,** Distribution of cells from Liv and Spl assigned to subcluster. Data are pooled from 20-25 mice per infection per tissue.

**Extended Data Figure 3.**
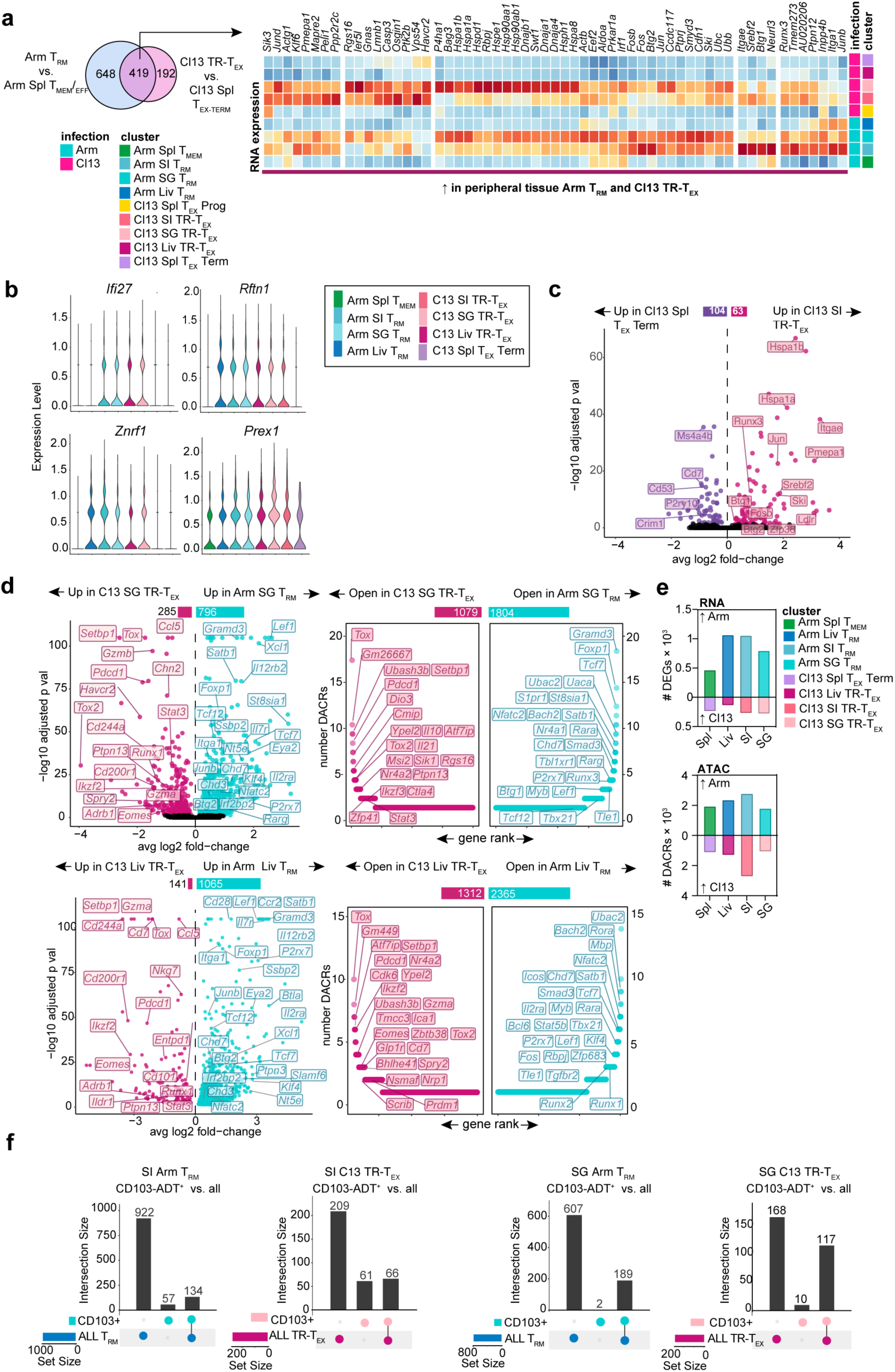
Transcriptional and epigenetic differences between T_RM_ and TR-T_EX_ cells. **a,** Heatmap of RNA expression of genes that are upregulated in Arm T_RM_ cells versus Arm Spl T_CIRCM_ cells and are also upregulated in Cl13 TR-T_EX_ cells versus Cl13 Spl T_EX_-_TERM_ cells. **b,** Violin plots displaying gene expression in indicated Seurat clusters. **c,** Volcano plot of differentially expressed genes (DEGs) between Arm SI TR-T_EX_ and Cl13 Spl T_EX_-_TERM_ cells. **d,** Volcano plots displaying pairwise DEGs (left column) and rank-ordered plots displaying pairwise differentially accessible chromatin regions (DACRs) (right column) between Arm T_RM_ cells (blue) and Cl13 TR-T_EX_ cells (pink) from SG (top panel) or Liv (bottom panel). **e,** Number of DEGs (left panel) and DACRs (right panel) between Arm Spl T_MEM_ and Cl13 Spl T_EX_-Term or tissue-matched Arm T_RM_ (blue) and Cl13 TR-T_EX_ (pink) P14 cells. **e,** Proportion of genes upregulated or downregulated in CD103-ADT^+^ Arm T_RM_ or CD103-ADT^+^ Cl13 TR-T_EX_ cells from the SI or SG that are also up- or down-regulated by total Arm T_RM_ or Cl13 TR-T_EX_ cells from the same tissue. Data are pooled from 20-25 mice per infection per tissue.

**Extended Data Figure 4.**
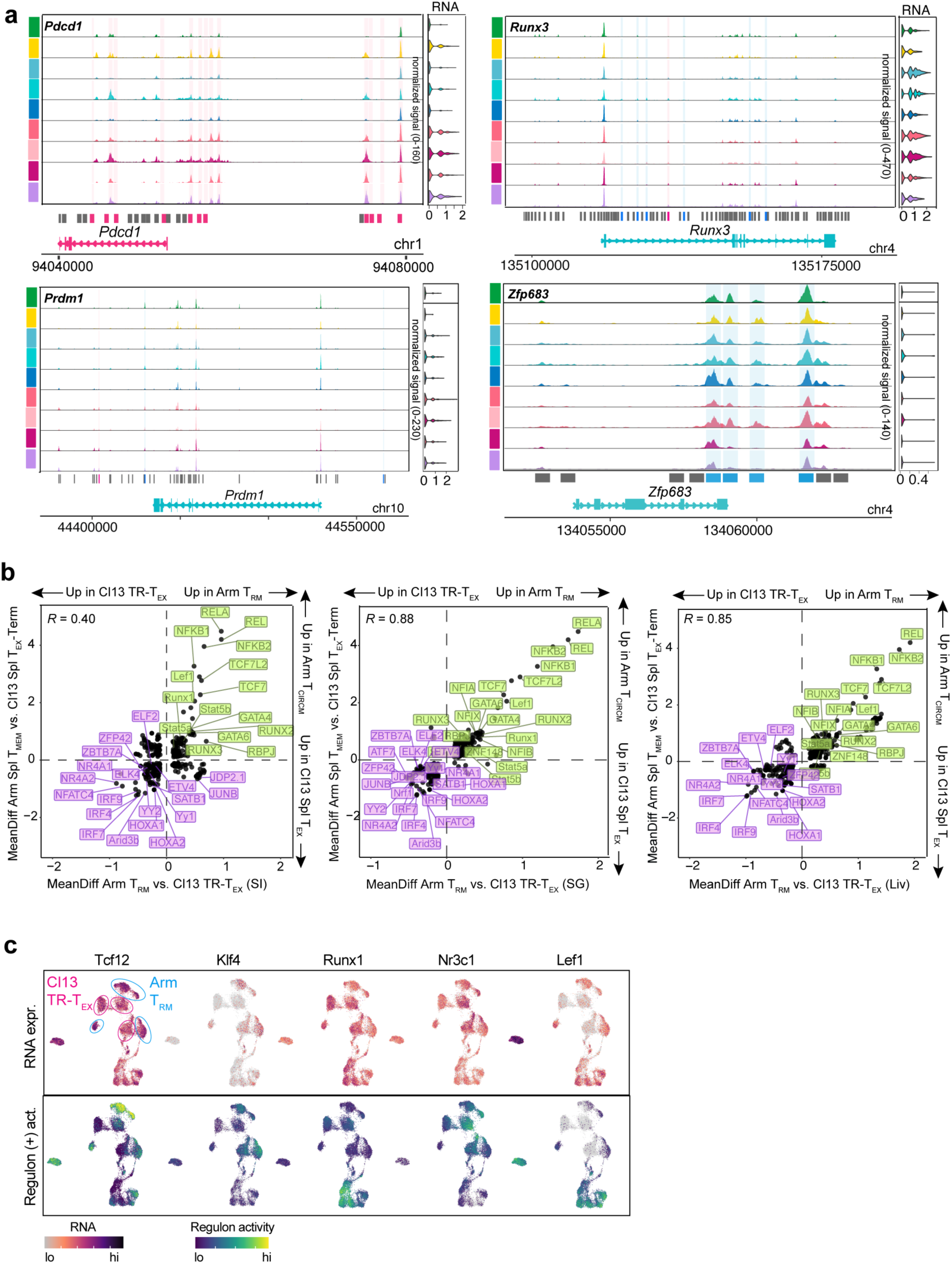
Differential regulation of exhaustion, memory and tissue-residency associated factors in T_RM_ and TR-T_EX_ cells. **a,** ATAC coverage plots of DACRs in indicated gene loci for each cluster. SI and SG Arm T_RM_ and Cl13 TR-T_EX_ are subsetted to CD103-ADT^+^ cells. Green bars indicate DACRs enriched in Arm Spl T_MEM_ compared to Arm T_RM_ cells, blue bars indicate DACRs enriched in Arm T_RM_ cells compared to location-matched Cl13 TR-T_EX_ cells from at least one tissue, pink bars indicate DACRs enriched in Cl13 TR-T_EX_ compared to location-matched Arm T_RM_ cells from at least one tissue. **b,** Pairwise transcription factor (TF) motif enrichment in SI (left panel), SG (middle panel) or Liv (right panel) Arm T_RM_ cells compared to tissue-matched Cl13 TR-T_EX_ cells (x axis) plotted against pairwise TF motif enrichment in Arm Spl T_MEM_ cells compared to Cl13 Spl T_EX_-_TERM_ cells (y axis). MeanDiff = mean difference determined by chromvar motif deviation analysis. **c,** RNA expression of indicated TFs (upper panel) and of genes in Pando-defined regulons predicted to be controlled by each TF (positive regulon activity). Data are pooled from 20-25 mice per infection per tissue.

**Extended Data Figure 5.**
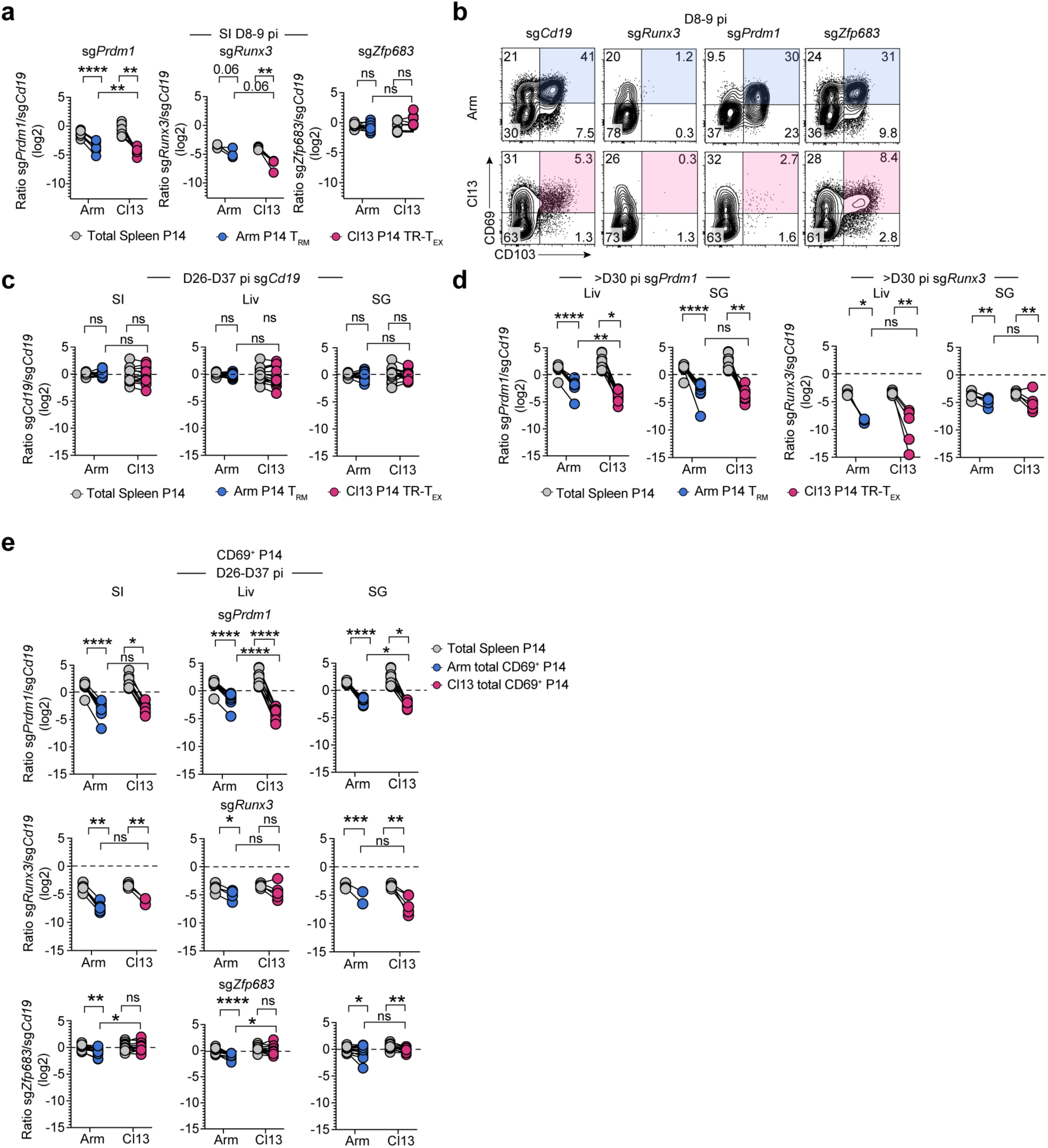
Requirements for Runx3, Blimp1 and Hobit for residency programming across tissues. **a,** Ratio of total P14 cells electroporated with indicated sgRNAs versus control *Cd19* sgRNAs from Arm (blue) or Cl13 (pink) infected SI (colored) or spleen (grey) at 8-9 dpi..**b,** Co-expression of CD69 and CD103 by SI P14 T cells electroporated with control sg*Cd19* or indicated TF-targeting sgRNAs at 8-9 dpi with Arm (blue) or Cl13 (pink). **c,** Ratio of congenically distinct and co-transferred P14 Arm T_RM_ (blue), TR-T_EX_ (pink) cells electroporated with identical control *Cd19* sgRNAs (same guide in each congenic population) in indicated tissues (SI, SG P14 gated on CD69^+^CD103^+^, Liv P14 gated on CD69^+^CXCR6^+^) compared to co-transferred total Spl-derived P14 cells (grey) at 26-37 dpi. **d,** Ratio of co-transferred Arm T_RM_ (blue), Cl13 TR-T_EX_ (pink) or total Spl-derived (grey) P14 cells electroporated with *Prdm1* or *Runx3* sgRNAs versus control *Cd19* sgRNAs at 30-37 dpi. Arm T_RM_ and Cl13 TR-T_EX_ were gated as CD69^+^CD103^+^ in the SG and as CD69^+^CXCR6^+^ in the Liv. **e,** Ratio of total CD69^+^ P14 cells electroporated with indicated sgRNAs versus control *Cd19* sgRNAs from Arm (blue) or Cl13 (pink) infected tissues (SI, Liv, SG; colored) compared to ratio of total Spl-derived P14 cells (grey) at 26-37 dpi. Data are pooled from 2 independent experiments with 4-8 mice per group per experiment. * p < 0.05, ** p < 0.01, *** p < 0.001, **** p < 0.0001 paired T test (spleen versus tissue P14s) or two-tailed T test (Arm versus Cl13 tissue P14s) **(a, c-e).**

**Extended Data Figure 6.**
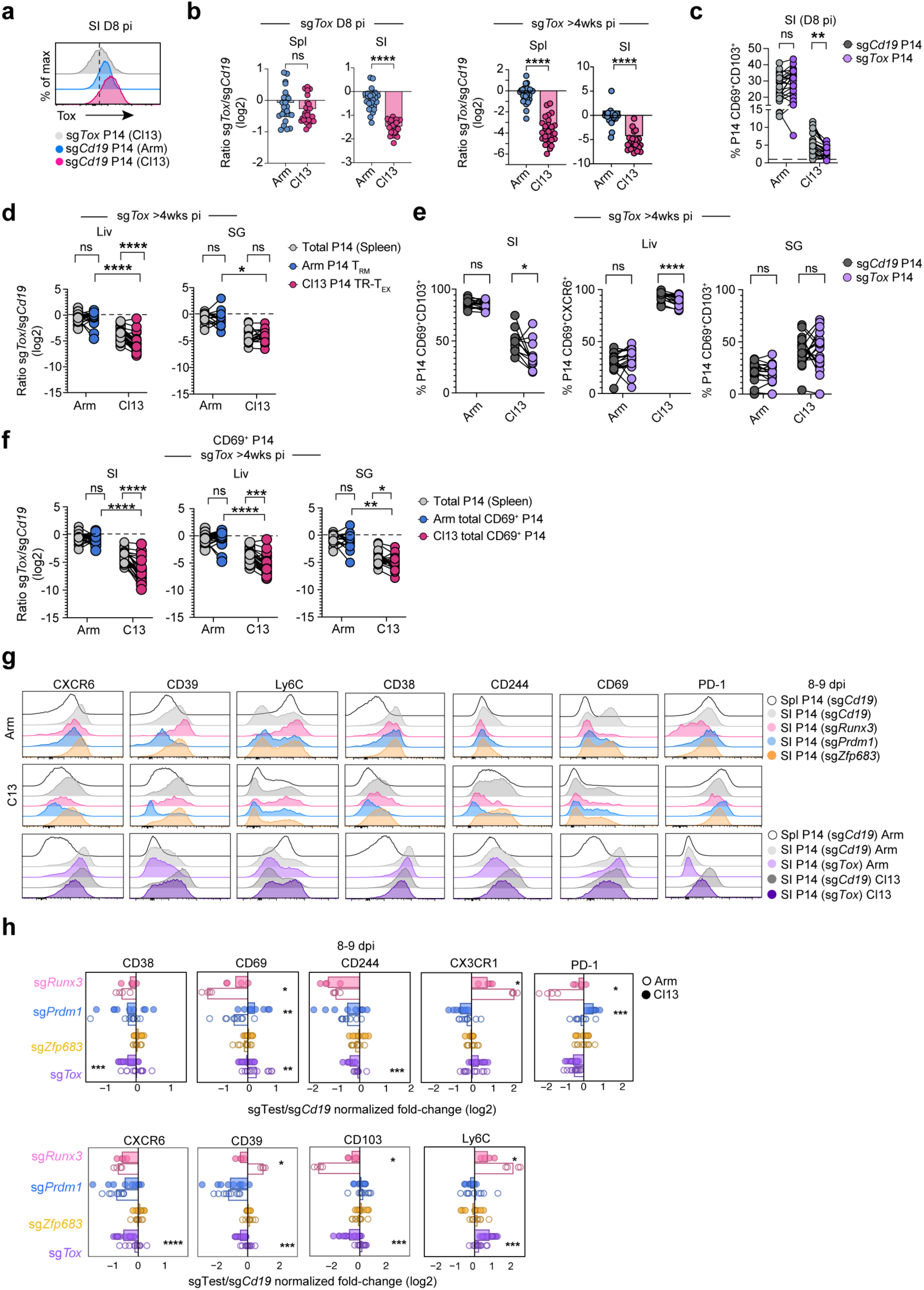
Tox coordinates TR-T_EX_ cell residency programming but is not required for T_RM_ cell development. **a,** Expression of Tox in total sg*Cd19* and sg*Tox* electroporated P14 cells isolated from the SI at 8-9 dpi with Arm or Cl13. **b,** Ratio of total P14 cells electroporated with *Cd19* control or *Tox* targeting sgRNAs isolated from the Spl or SI at 8d (left panel) or >4wks (right panel) pi with Arm or Cl13. **c,** Frequency of CD69^+^CD103^+^ sg*Cd19* (grey) or sg*Tox* (purple) electroporated P14 cells isolated from the SI at 8 dpi. **d,** Ratio of co-transferred Arm T_RM_ (blue), Cl13 TR-T_EX_ (pink) or total Spl-derived (grey) P14 cells electroporated with *Tox* sgRNAs versus control *Cd19* sgRNAs >4wks pi. Arm T_RM_ and Cl13 TR-T_EX_ were both gated as CD69^+^CD103^+^ in the SG or as CD69^+^CXCR6^+^ in the Liv. **e**, Frequency of P14 cells that were CD69^+^CD103^+^ (SI, SG) or CD69^+^CXCR6^+^ (Liv) following electroporation with *Tox* (purple) or *Cd19* (grey) targeting sgRNAs isolated from indicated tissues >4wks pi. **f,** Ratio of total CD69^+^ P14 cells electroporated with *Tox* sgRNAs versus control *Cd19* sgRNAs from Arm (blue) or Cl13 (pink) infected tissues (SI, Liv, SG; colored) compared to ratio of total P14 cells in the spleen (grey) >4wks pi. **g,** Expression of indicated surface molecules by total P14 cells electroporated with indicated sgRNAs and isolated from the SI 8-9 dpi with Arm or Cl13. **h,** Fold change (FC) in expression of indicated surface molecules in total P14 cells electroporated with sgRNAs directed towards genes encoding indicated TFs compared to control cells electroporated with sgRNAs directed towards *Cd19* at 8-9 dpi with Arm or Cl13. Data are pooled from 2-3 experiments with 4-8 mice per group per experiment. * p < 0.05, ** p < 0.01, *** p < 0.001, **** p < 0.0001 paired T test (spleen versus tissue P14s) or two-tailed T test (Arm versus Cl13 tissue P14s) **(b-f)** or Mann Whitney test (**h**).

**Extended Data Figure 7.**
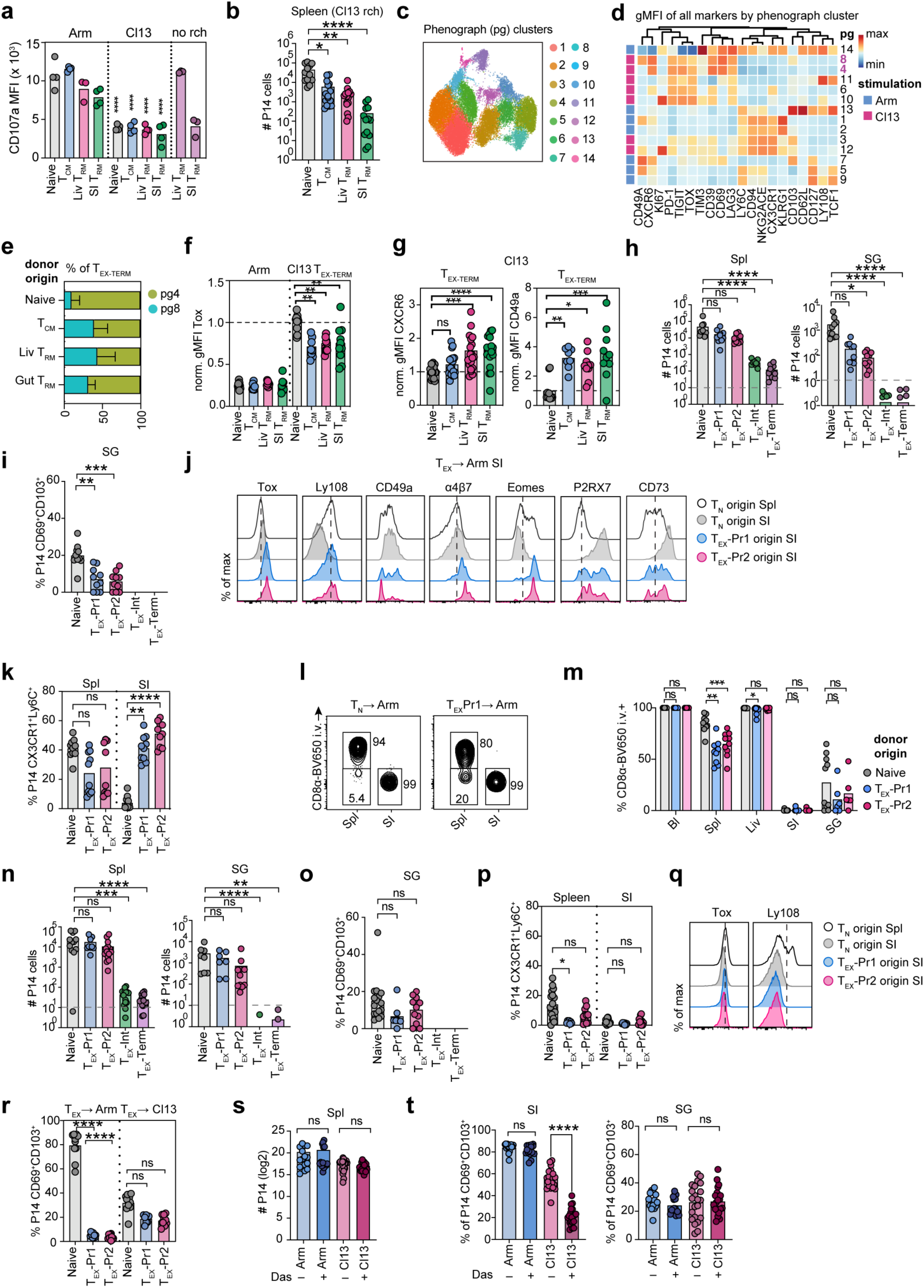
Developmental plasticity and lineage relationships between T_RM_ and T_EX_ cells. **a,** Expression of CD107a by sort-transferred donor P14 cells rechallenged with Arm or Cl13 and re-isolated from the Spl 21-28 dpi following *ex vivo* gp_33-44_ stimulation, compared to T_RM_ cells that were not transferred or rechallenged (no rch). Statistics indicate comparison between Arm and Cl13 rechallenged cells of same origin. **b,** Absolute number of total P14 cells recovered from the Spl of mice receiving donor cells from indicated origin following Cl13 rechallenge. **c,** UMAP of unsupervised phenograph (pg) clustering of donor P14 cells from Arm or Cl13 rechallenged mice; numbers in key indicate pg cluster determined by kmeans clustering. **d,** Heatmap of marker expression in pg clusters. Rechallenge infection giving rise to each cluster is indicated by pink (Cl13) or blue (Arm). T_EX_-_TERM_ clusters are highlighted in purple (pg 8 and pg 4). **e,** Proportion of Spl T_EX_-_TERM_ cells derived from donor P14 cells of indicated origin in pg clusters after Cl13 rechallenge. **f, g,** Expression of markers by Spl T_EX_-_TERM_ cells from indicated donor origin after Cl13 rechallenge compared to Arm rechallenge. **h,** Absolute number of total P14 cells (Spl) or CD69^+^CD103^+^ P14 cells (SG) derived from donor P14 cells following Arm rechallenge. **i,** Proportion of Arm-rechallenged P14 cells co-expressing CD69 and CD103 in SG. **j,** Expression of indicated surface molecules by Arm-rechallenged donor P14 cells of indicated origin isolated from the Spl or SI. **k**, Proportion of Arm-rechallenged P14 cells in the Spl or SI expressing CX3CR1 and Ly6C. **l, m,** Proportion of Arm-rechallenged donor P14 cells staining positive for CD8a-BV650 during intravascular labelling. Bl; blood. **n,** Absolute number of total P14 cells (Spl) or CD69^+^CD103^+^ P14 cells (SG) derived from donor P14 cells following Cl13 rechallenge. **o,** Proportion of Cl13-rechallenged donor P14 cells co-expressing CD69 and CD103 in SG. **p,** Expression of Ly6C and CX3CR1 by Cl13-rechallenged donor P14 cells recovered from the SI. **q,** Expression of indicated surface molecules by Cl13 rechallenged donor P14 cells isolated from the Spl or SI. **r**, Proportion of donor P14 cells expressing CD69 and CD103 in the SI after Arm or Cl13 rechallenge. **s,** Absolute number of total P14 cells isolated from the Spl of Arm or Cl13 infected mice following 1wk dasatinib treatment. **t,** Frequency of P14 cells co-expressing CD69 and CD103 in SI and SG of Arm or Cl13 infected mice following 1wk dasatinib treatment. Dashed lines on absolute number plots (**h, n**) indicate threshold limit of detection for plotting in frequency plots. Dashed lines in histograms (**l, q**) indicate average expression in total naïve P14-derived cells in Spl. Data are pooled from or representative of 2 independent experiments **(a, h-r)** or 3 independent experiments **(b-g, s-t)**, with n = 3-5 mice per group per experiment. T_N_; naïve P14 cells. * p < 0.05, ** p < 0.01, *** p < 0.001, **** p < 0.0001 Mann Whitney Test.

**Extended Data Figure 8.**
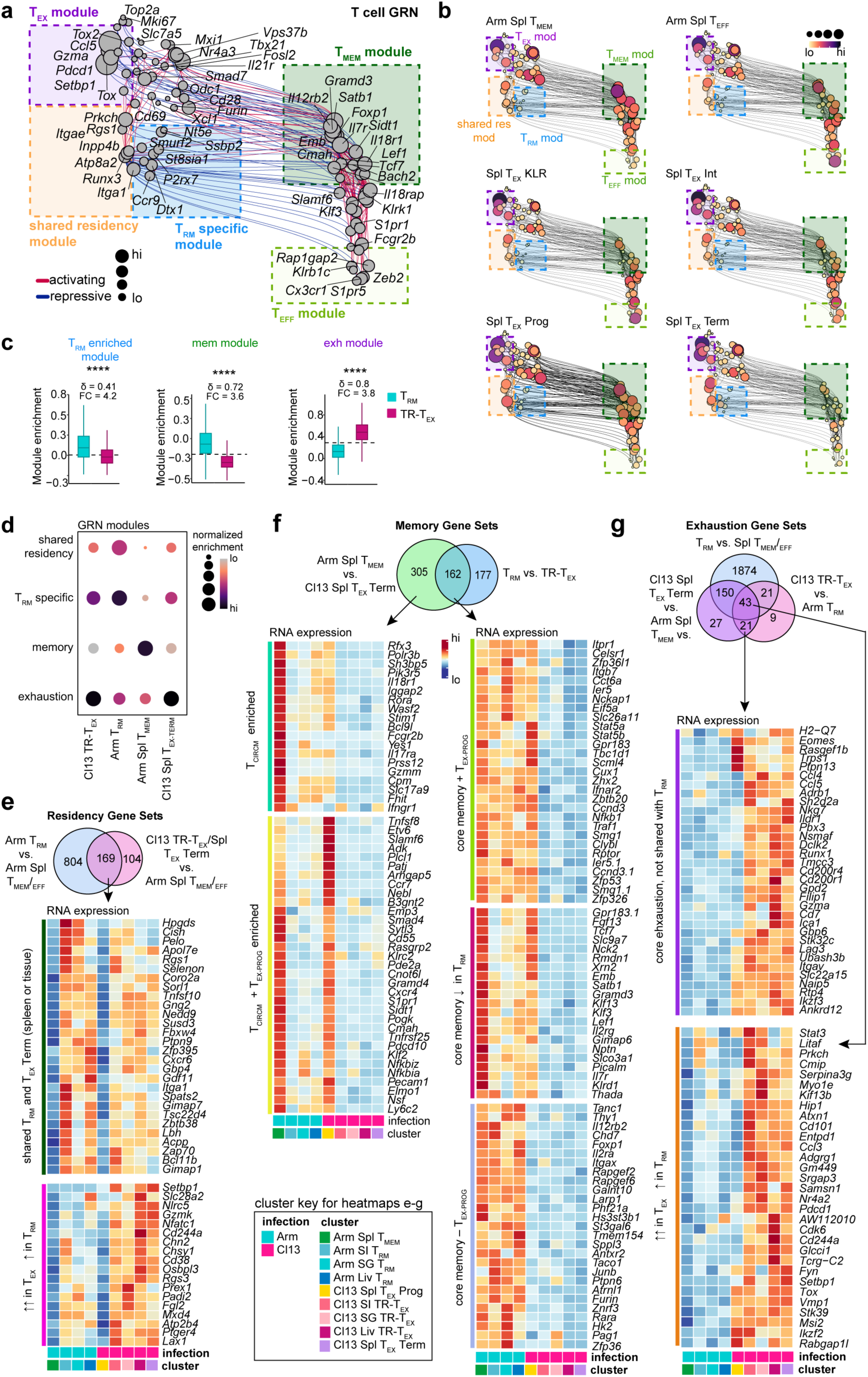
Cell-state specification of T_RM_ and TR-T_EX_ cells. **a,** Detailed UMAP embedding of inferred single-cell Gene Regulatory Network (GRN) based on combined transcription factor (TF) expression and motif accessibility in all non-naïve P14 cells analyzed by TEA-sequencing. Size of nodes (genes) represents the number of connections in network. **b,** Individual UMAP embeddings of genes and gene modules from inferred single-cell Gene Regulatory Networks (GRNs) engaged in each T cell subset. Node size and color scale represent degree of RNA expression for each gene in network. **c, d** Enrichment for GRN modules in merged T_RM_ and TR-T_EX_ cell Seurat clusters from all tissues, or in Arm Spl T_MEM_ and Cl13 Spl T_EX_-_TERM_ clusters. FC = fold change, δ = Cliff’s delta effect size. Circle size and heat scale indicate relative enrichment per cluster. **e,** Heatmap of RNA expression of genes commonly upregulated in Arm T_RM_ and Cl13 TR-T_EX_ cells versus Arm Spl T_CIRCM_ cells (pooled Arm Spl T_MEM_ and Arm Spl T_EFF_ clusters). **f,** Heatmap of RNA expression of genes uniquely upregulated in Arm Spl T_CIRCM_ cells versus Cl13 Spl T_EX_-_TERM_ cells but not in Arm T_RM_ cells versus Cl13 TR-T_EX_ cells (left panel), or commonly upregulated in Arm Spl T_CIRCM_ cells versus Cl13 Spl T_EX_-_TERM_ cells and Arm T_RM_ cells versus Cl13 TR-T_EX_ cells (right panel). **g,** Comparative analysis of RNA expression of genes upregulated in 1) Cl13 Spl T_EX_-T_ERM_ cells versus Arm Spl T_CIRCM_ cells, 2) Cl13 Spl TR-T_EX_ cells versus Arm T_RM_ cells from each tissue and 3) T_RM_ from each tissue versus Arm Spl T_CIRCM_ cells. Heatmaps display genes that are uniquely enriched in T_EX_ (top panel) or are shared with Arm T_RM_ cells (bottom heatmap) compared to Spl T_CIRCM_ cells. Data are pooled from 20-25 mice per infection per tissue.

**Extended Data Figure 9.**
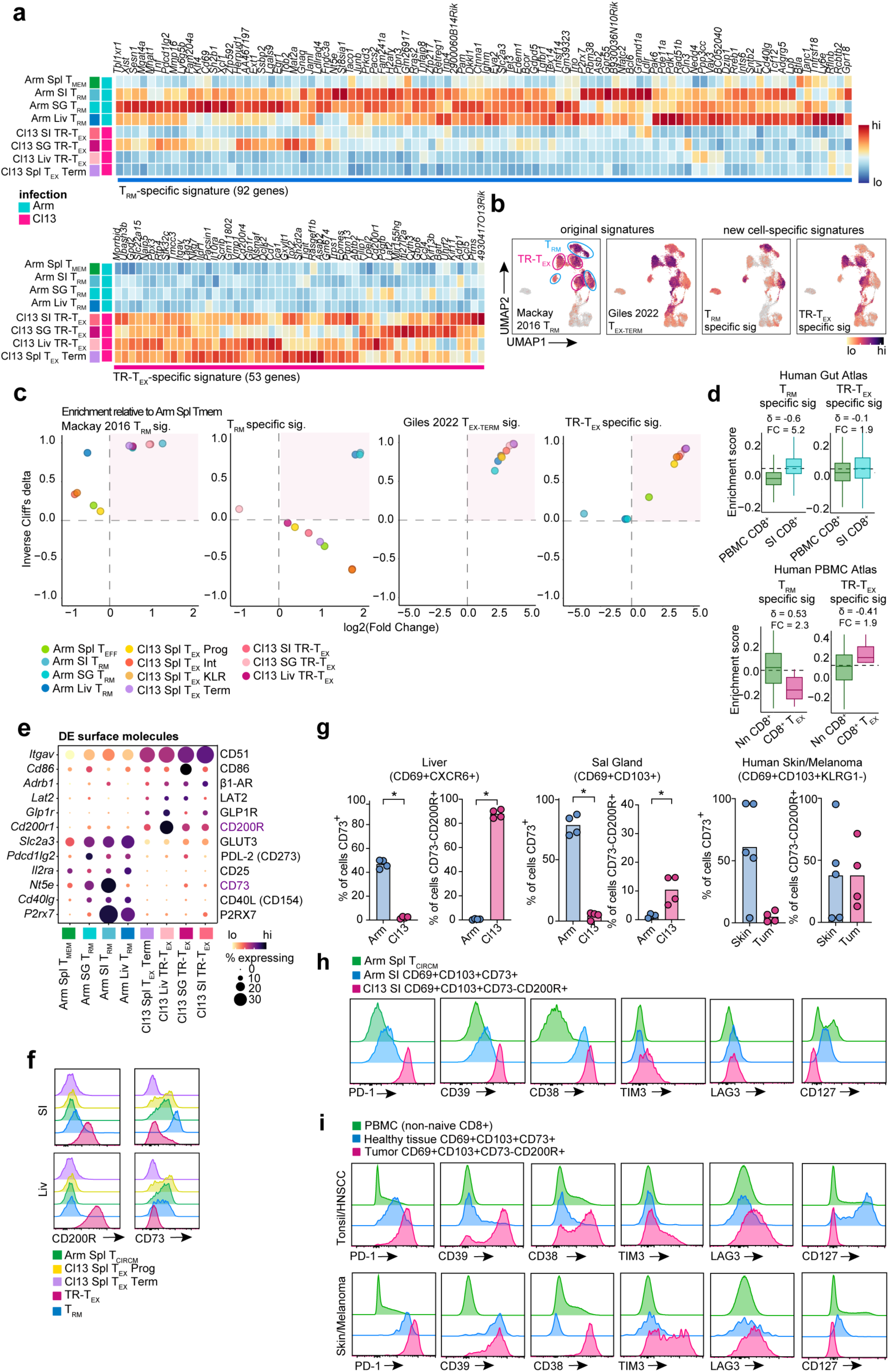
Defining cell-state specific gene and surface protein signatures of T_RM_ and TR-T_EX_ cells. **a,** Heatmaps displaying relative RNA expression of indicated genes comprising T_RM_ cell-specific (upper panel) or TR-T_EX_ cell-specific (lower panel) signatures identified through TEA-seq. **b,** UMAP projections showing Seurat enrichment for published Mackay 2016 T_RM_ core and Giles 2022 T_EX-TERM_ signatures in T_RM_ (blue circled clusters) and TR-T_EX_ (pink circled clusters) compared to newly derived T_RM_ and TR-T_EX_ cell-state specific signatures in TEA-seq dataset. **c,** Cliff’s delta effect size and fold-change of Seurat enrichment for published Mackay 2016 T_RM_ core and Giles 2022 T_EX-TERM_ signatures and newly derived T_RM_ and TR-T_EX_ cell-state specific signatures in indicated P14 cell populations from TEA-seq dataset. **d,** Seurat enrichment for cell-state specific T_RM_ or TR-T_EX_ signature scores within CD8^+^ T cells isolated from human SI epithelium or non-naïve CD8^+^ T cells from healthy donor peripheral blood^60^ or GSEA enrichment for cell-state specific signatures within CD39^+^PD-1^+^ T_EX_ cells compared to all non-naïve CD8^+^ T cells isolated from healthy donor peripheral blood^61^. δ = Cliff’s delta, FC = fold-change. **e,** RNA expression of genes encoding indicated surface proteins in LCMV-generated P14 cell clusters identified via TEA-seq. Circle size indicates proportion of cells expressing each gene; scale bar indicates relative expression. **f,** Expression of indicated molecules in Arm Spl T_CIRCM,_ Cl13 Spl T_EX_-_PROG_ or Cl13 Spl T_EX_-_TERM_ cells and in T_RM_ or TR-T_EX_ cells isolated from SI (CD69^+^CD103^+^) or Liv (CD69^+^CXCR6^+^) 30-40 dpi with Arm or Cl13, respectively. **g,** Proportion of T_RM_ (blue) or TR-T_EX_ cells (pink) expressing CD73 and CD200R in Liv (CD69^+^CXCR6^+^) or SG (CD69^+^CD103^+^) after Arm or Cl13 infection, respectively (left and middle panel) or proportion of T_RM_-like cells in human epidermal or melanoma samples (KLRG1^-^CD69^+^CD103^+^) expressing CD73 and CD200R (right panel). **h,** IR expression by total P14 T_CIRCM_ from the Spl of Arm-infected mice or by CD69^+^CD103^+^CD73^+^ T_RM_ cells (blue) or CD69^+^CD103^+^CD73^-^CD200R^+^ TR-T_EX_ cells (pink) 30-40 dpi with Arm or Cl13, respectively. **i,** IR expression by KLRG1^-^CD69^+^CD103^+^CD73^+^ T_RM_-like cells from human tonsil or epidermal skin or KLRG1^-^CD69^+^CD103^+^CD73^-^CD200R^+^ T_RM_-like TIL from HNSCC or melanoma. Data are pooled from 3-6 human donors (**g, i**) or representative of at least 2 experiments with 4-5 mice per group per experiment (**e, f, g**). TEA-seq data are pooled from 20-25 mice per infection per tissue. * p < 0.05, Mann Whitney test.

**Extended Data Figure 10.**
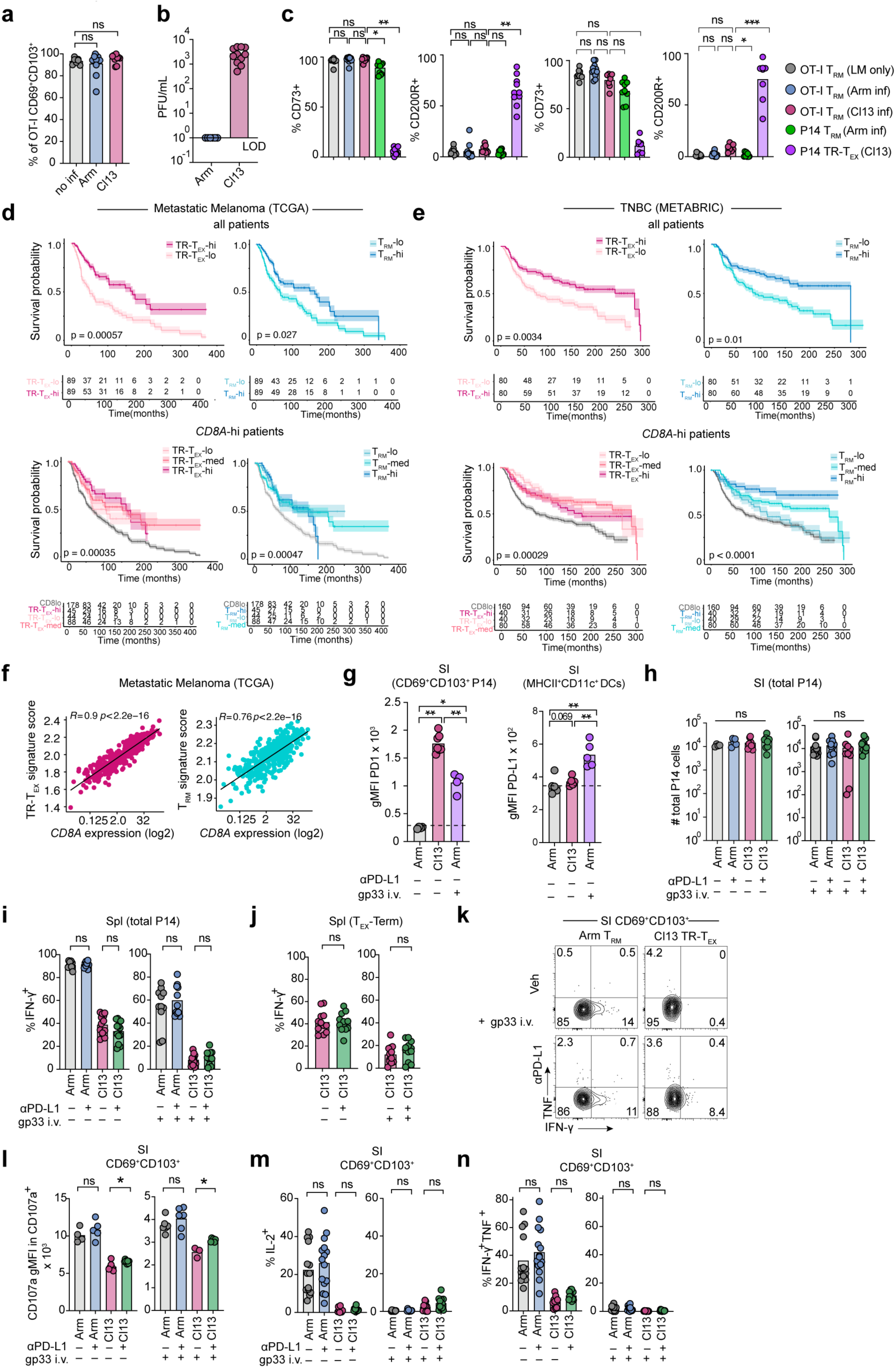
T_RM_ and TR-T_EX_ cells differentially contribute to immune responses. **a,** Frequency of LM-OVA generated OT-I cells in the SI co-expressing CD69 and CD103 following LCMV Arm or Cl13 reinfection, or with no LCMV infection (no inf). **b,** Viral titers in the serum of LM-OVA immune mice 30d after infection with Arm or Cl13. LOD = limit of detection. **c,** Proportion of CD69^+^CD103^+^ OT-I T_RM_ (LM-derived), P14 T_RM_ (Arm-derived) or P14 TR-T_EX_ (Cl13-derived) cells expressing CD73 or CD200R following LCMV infection or without LCMV infection (LM only). **d, e** Kaplan-Meier survival curves for overall survival of metastatic melanoma patients from the TCGA database (**d**) or TNBC patients from the METABRIC database (**e**) displaying stratification by T_RM_ and TR-T_EX_ specific signatures (^hi^ = top 25%, ^lo^ = bottom 25%) within all patients (upper panel) or within the top 50% of CD8^hi^ patients compared to CD8^lo^ patients (lower panel). **f,** Correlation analysis comparing T_RM_ (blue) and TR-T_EX_ (pink) signature scores and *CD8A* expression in TCGA melanoma patients. **g,** Geometric mean fluorescence intensity (gMFI) of PD-1 in CD69^+^CD103^+^ Arm SI P14 T_RM_ or Cl13 TR-T_EX_ cells (left panel) and of PD-L1 in MHCII^+^CD11c^+^ SI-derived dendritic cells (DCs) at baseline (no peptide treatment) and 48h after treatment with gp_33-44_ peptide i.v.. **h,** Number of total P14 cells isolated from the SI (right panel) of Arm or Cl13 infected mice treated with FTY720 and α-PD-L1 with or without i.v. gp33_33-41_ peptide administration 48h earlier. **i, j,** Cytokine production by total Spl P14 cells (**i**) or by CD69^+^CXCR6^+^ Cl13 Spl T_EX_-_TERM_ cells (**j**) treated with α-PD-L1 with or without i.v. gp33_33-41_ peptide treatment. **k,** Frequency of CD69^+^CD103^+^ SI P14 T_RM_ (Arm) or TR-T_EX_ (Cl13) cells producing IFN-γ and TNF following α-PD-L1 treatment with gp_33-44_ peptide i.v. co-infusion in all mice. **l-n,** Degranulation (CD107a expression, l) or cytokine production by SI CD69^+^CD103^+^ P14 Arm T_RM_ cells or Cl13 TR-T_EX_ cells from mice treated with α-PD-L1 with or without gp33_33-41_ peptide i.v. . Data are pooled from and representative of 3 independent experiments with n = 4-7 mice per group per experiment (**a-c**) and 2-3 independent experiments with 3-8 mice per group per experiment (**g-n**). * p < 0.05, ** p < 0.01, Kruskal Wallist Test (**a-c, g**), Mann Whitney test (**h-n**), Kaplan-Meier estimate (**d, e**) or Pearson correlation test (**f**).

