## Supplementary Material 1 for "Divergent ontogeny of Tissue Resident Memory and Tissue Resident Exhausted CD8^+^ T cells underlies distinct functional potential"

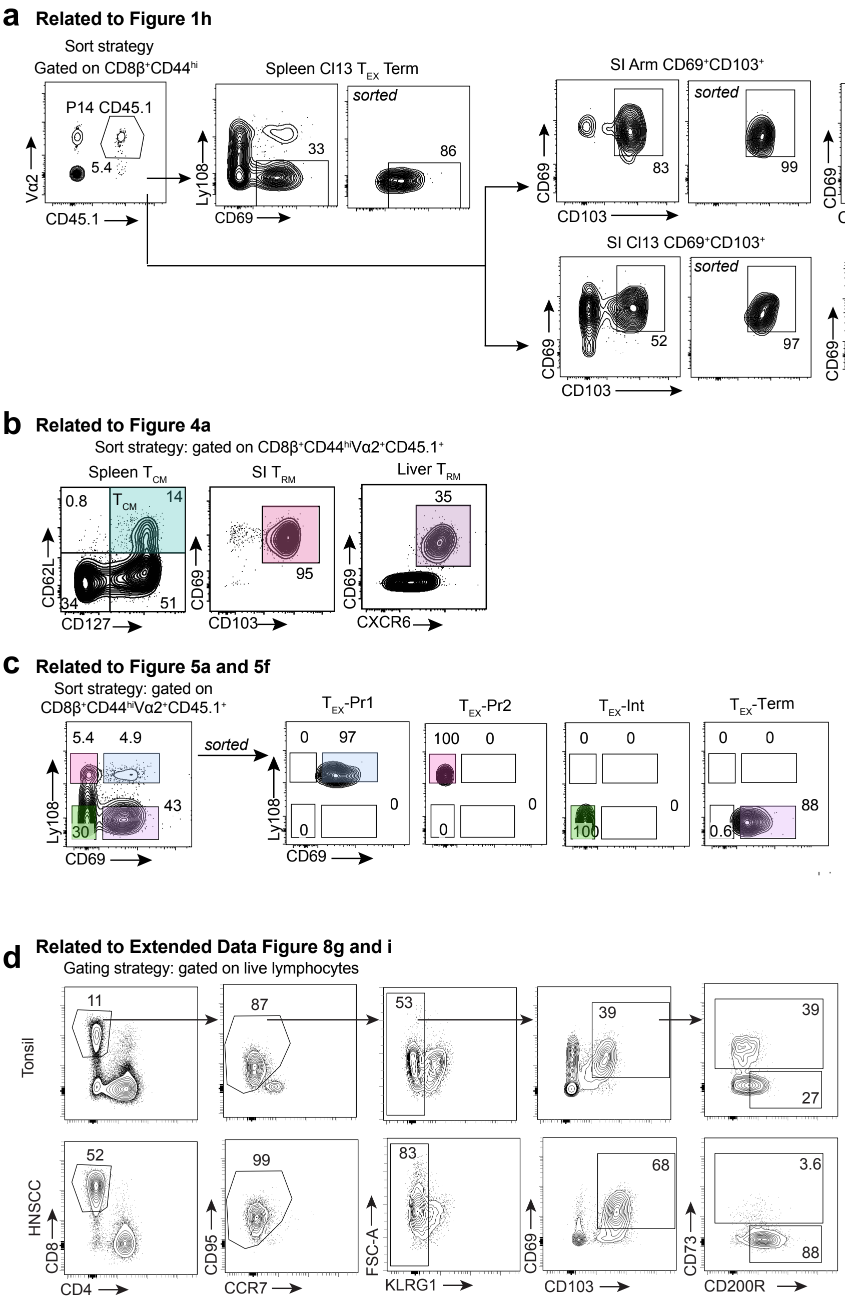


**Supplementary Material 1. a**, Gating strategy for sort-retransfer experiments outlined in Figure 1h. Cells were pre-gated on CD8β$^+$CD44$^{hi}$CD45.1$^+$Vα2$^+$ P14 cells and then sorted as: Ly108$^-$CD69$^+$ Cl13 Spleen P14 (Cl13 Spl T$_{EX-TERM}$), CD69$^+$CD103$^+$ Arm SI P14 (SI T$_{RM}$) or CD69$^+$CD103$^+$ Cl13 SI P14 and CD69$^+$CXCR6$^+$CD62L$^-$ Arm Liv P14 (Liv T$_{RM}$) or CD69$^+$CXCR6$^+$CD62L$^-$ Liv Cl13 P14. **b,** Gating strategy for sort-retransfer experiments outlined in Figure 4a. Cells were pre-gated on CD8β$^+$CD44$^{hi}$CD45.1$^+$Vα2$^+$ P14 cells and then sorted as: CD62L$^+$CD127$^+$ Arm Spleen P14 (T$_{CM}$), CD69$^+$CD103$^+$ Arm SI P14 (SI T$_{RM}$), CD69$^+$CXCR6$^+$CD62L$^-$ Arm Liv P14 (Liv T$_{RM}$). **c,** Gating strategy for sort-retransfer experiments outlined in Figure 5a and 5f. Cells were pre-gated on CD8β$^+$CD44$^{hi}$CD45.1$^+$Vα2$^+$ P14 cells and then sorted as: Ly108$^+$CD69$^+$ Cl13 Spleen P14 (Cl13 Spl T$_{EX}$-Progenitor 1, T$_{EX}$-Pr1), Ly108$^+$CD69$^-$ Cl13 Spleen P14 (Cl13 Spl T$_{EX}$-Progenitor 2, T$_{EX}$-Pr1), Ly108-CD69- Cl13 Spleen P14 (Cl13 Spl T$_{EX}$-Intermediate, T$_{EX-INT}$), Ly108$^-$CD69$^+$ Cl13 Spleen P14 (Cl13 Spl T$_{EX-TERM}$). **d,** Gating strategy for human flow cytometry experiments shown in Extended Data Figure 8g and i. Cells were pre-gated on live lymphocytes then gated as shown to assess CD73 and CD200R co-expression in CD8$^+$ T cells expressing residency-associated molecules.
